# Analysis of Small Signaling Peptides in *Sorghum bicolor*: Integrating Phylogeny and Gene Expression to Characterize Roles in Stem Development

**DOI:** 10.64898/2026.03.25.714272

**Authors:** Evan Kurtz, Brian McKinley, John Mullet

## Abstract

Small signaling peptides (SSPs) are critical regulators of plant growth, development, and responses to biotic and abiotic stress, yet their role in the C4 grass *Sorghum bicolor* is largely uncharacterized. To help fill this knowledge gap, 219 *S. bicolor* genes that encode SSPs were identified based on SSP sequences previously identified in *Arabidopsis thaliana*, *Oryza sativa*, *Zea mays*, *Triticum aestivum*, and *Brachypodium distachyon*. The 219 sorghum genes were assigned to 19 gene families, analyzed for the presence of motifs, and aligned with genes that encode SSPs in other plants using phylogenetic analysis. Expression of the 219 SSP encoding genes in sorghum organs, during stem development, and in stem tissues and cell types revealed distinct spatial, temporal and developmental patterns of expression. Genes associated with the *SbCEP* and *SbRGF* families were preferentially expressed in roots, whereas *SbEPF* genes were expressed in stems and panicles. The expression of genes during bioenergy sorghum stem growth and development was investigated because stems account for ∼80% of harvested biomass and serve as conduits for water and nutrient transport between leaves and roots. During stem development, 28 SSP encoding sorghum genes in several families (*CLE, EPF, CEP, GASS, PSY, ES, PSK, CAPE, POE*) were expressed at higher levels in zones of cell proliferation. For example, the TDIF homologs S*bCLE41* and *SbCLE42* were expressed at high levels in nascent stem nodes where they may regulate cambial activity and vascular bundle cell differentiation. A different set of 15 genes in the *CIF, POE, CAPE, PSY, CEP, RALF,* and *CLE* families were expressed at higher levels in zones of stem tissue differentiation highlighted by elevated expression of 5 *SbRALF*s in the stem nodal plexus. Cell type specific expression of many SSP encoding sorghum genes was also observed in fully elongated internodes indicating gene expression is regulated with high spatial resolution. Overall, the results provide a foundation of information for analysis of SSP functions in sorghum that can be integrated with knowledge of sorghum gene regulatory networks to modulate traits important for production of sorghum crops.

## Introduction

Bioenergy sorghum (*Sorghum bicolor*) is a highly productive drought tolerant C4 grass that is well adapted to annual U.S. cropland designated for bioenergy crops (Rooney, Blumenthal et al. 2007, Mullet, Morishige et al. 2014, Langholtz, Davis et al. 2024). Sorghum’s C4 photosynthesis enhances its productivity, reduces nitrogen requirements and water use (Byrt, Grof et al. 2011) while its deep and extensive root system contributes to nutrient uptake and deposition of soil organic carbon (Lamb, Weers et al. 2022). Bioenergy sorghum’s productivity, low carbon intensity (Olson, Ritter et al. 2012, Gautam, Mishra et al. 2020), low input requirements (Olson, Ritter et al. 2013) and resilience make it a promising bioenergy crop for annual bioenergy cropland (Rooney, Blumenthal et al. 2007, Maw, Houx et al. 2017).

Bioenergy sorghum hybrids are very photoperiod sensitive which delays flowering in long days increasing the duration of vegetative phase growth and biomass yield. At the end of the growing season bioenergy sorghum stems are often 4-5 m long and account for ∼80% of harvested biomass (Olson, Ritter et al. 2012). Stem biomass is comprised of cell walls composed primarily of cellulose, glucuronoarabinoxylan (GAX), and lignin and non-structural carbohydrates such as sucrose, glucose, fructose, starch, and mixed-linkage glucans (MLGs) (McKinley, Rooney et al. 2016, McKinley, Olson et al. 2018). Sugars derived from cellulose and non-structural carbohydrates can be readily converted into biofuels such as ethanol, butanol, and sustainable aviation fuel (SAF) and other bioproducts.

Grass stems are composed of nodes and internodes. Stem nodes span the pulvinus, a tissue that produces tiller buds (Kebrom, McKinley et al. 2017) and nodal root buds (Lamb, Kurtz et al. 2024), and the nodal plexus, stem tissue located below the pulvinus where the leaf sheath joins the stem. Stem internode tissue develops between nodes from an intercalary meristem located at the base of the internode just above the pulvinus (Serrano-Mislata and Sablowski 2018, McKim 2019, Yu, McKinley et al. 2021). During the vegetative phase, a new stem node is formed below the shoot apical meristem approximately every 3-4 days. Therefore, at the end of bioenergy sorghum’s long vegetative growing season, plants contain >30 node-internode stem segments (Olson, Ritter et al. 2012). The formation, growth, and development of stem node-internode segments is a key determinant of bioenergy sorghum biomass accumulation since longer, thicker and more dense stems contribute to greater stem biomass accumulation (Kebrom, McKinley et al. 2017, McKinley, Olson et al. 2018, Dos Santos, Fernandes et al. 2020).

Stem internode growth is regulated by genetic, hormonal, and environmental factors. Sorghum *dwarfing* loci that are used to reduce grain sorghum stem length provided early insights into the regulation of stem internode growth. Sorghum *Dw3* encodes an ABCB19 IAA export transporter (Multani, Briggs et al. 2003), *Dw1* encodes a protein involved in brassinosteriod signaling (Hilley, Truong et al. 2016, Yamaguchi, Fujimoto et al. 2016) and *Dw2* encodes an AGCVIII kinase (Hilley, Weers et al. 2017) that modulates endomembrane trafficking (Oliver, Fan et al. 2021). Shade avoidance responses mediated by PhyB-signaling increases the length of sorghum internodes in part by increasing expression of *GA3ox2* and through gibberellin (GA) enhanced stem internode elongation ((Beall, Morgan et al. 1991, Yu, McKinley et al. 2021). In addition, brassinosteroids (Mantilla Perez, Zhao et al. 2014), cytokinins (Amzallag, Lerner et al. 1992), and ethylene (Finlayson, Lee et al. 1999) also modulate stem growth.

Studies of sorghum stem development provided further insight into the molecular basis of stem growth regulation. Stem node-internode segments formed during the juvenile phase contain minimal internode tissue. The crown root system grows out from this initial stack of below ground stem nodes. In contrast, above ground stem nodes produced during the adult vegetative phase are separated from each other by internodes of varying length. The early stage of internode tissue development is characterized by active cell proliferation and elongation. This is followed by secondary cell wall formation, cell wall lignification and inactivation of the intercalary meristem (Kebrom, McKinley et al. 2017). Genes involved in IAA, GA, BR and CK hormone metabolism and signaling are differentially regulated during internode development. Analysis of gene expression in the stem internode intercalary meristem during sorghum internode development enabled the identification of a gene regulatory network that modulates the expression of genes involved in cell proliferation in response to changes in IAA, GA, BR and CK (Yu, Oliver et al. 2022).

Sorghum homologs of GA responsive *AtGASA4* and *AtGASA14* genes that encode SSPs were part of the gene regulatory network that regulated stem intercalary meristem activity (Yu, Oliver et al. 2022). GASA SSPs, characterized by their conserved cysteine-rich motifs, are regulated by gibberellins and influence cell proliferation, elongation, and stress adaptation (Herzog, Dorne et al. 1995, Bouteraa, Ben Romdhane et al. 2023). They mediate hormonal crosstalk between GA, brassinosteroids, and ABA (Chen, Sun et al. 2021), with roles in stem elongation (AtGASA2; (Bouteraa, Ben Romdhane et al. 2023), grain size regulation (OsGASR9; (Li, Shi et al. 2019), and stress tolerance (i.e., AtGASA11 in ROS scavenging) (Chen, Sun et al. 2021). Additional functions include zinc homeostasis (OsGASR10; (Nanda, Pujol et al. 2017), pathogen defense (OsGASR7;(Boonpa, Tantong et al. 2018), and reproductive development (AtGASA13; (Fan, Zhang et al. 2017).

A study of sorghum nodal root buds that form on the stem pulvinus identified several genes encoding RGF SSPs (Lamb, Kurtz et al. 2024). The RGF genes were part of a gene regulatory network that modulates root tissue formation prior to outgrowth (Lamb, Kurtz et al. 2024). RGF SSPs, also known as GOLVEN (GLV), regulate root meristem maintenance, differentiation, and hormonal signaling through interactions with RGFR receptor kinases (Stuhrwohldt, Scholl et al. 2020). They modulate auxin distribution, root gravitropism, and stem cell proliferation (AtRGF9 and AtRGF4;(Matsuzaki, Ogawa-Ohnishi et al. 2010, Whitford, Fernandez et al. 2012) and influence lateral root formation (AtRGF7; (Matsuzaki, Ogawa-Ohnishi et al. 2010, Zhong, Xie et al. 2020). Crosstalk with auxin and cytokinin further integrates RGFs into root patterning and stress responses (i.e., AtRGF6) (Matsuzaki, Ogawa-Ohnishi et al. 2010, Stuhrwohldt, Scholl et al. 2020).

SSPs, such as the sorghum RGFs and GASAs noted above, are critical regulators of plant growth and development, mediating intercellular communication, and modulation of meristem activity, vascular patterning, cell elongation, and stress responses. However, in our studies of sorghum development we found that genes encoding SSPs were incompletely annotated in Phytozome. For example, (Campbell and Turner 2017) identified genes encoding RALF SSPs in 51 plant species including sorghum. While that study identified 32 sorghum genes that encode RALFs, the analysis was done prior to the generation of a more complete and better annotated sorghum reference genome (McCormick, Truong et al. 2018). Given the potential importance of SSPs in bioenergy sorghum, we decided that further analysis of sorghum genes that encode SSPs was warranted. The goal of the current study is to identify and characterize the expression of sorghum genes that encode SSPs and to elucidate their relationships with gene homologs in *Arabidopsis* and other grasses. A more complete characterization of signaling pe genes, gene expression, and function in sorghum may elucidate useful ways to enhance bioenergy sorghum performance and sustainability.

## Materials and Methods

### Identification of sorghum genes that encode SSPs

SSPs, and the genes that encode them, have been identified in *Arabidopsis thaliana*, *Zea mays*, *Oryza sativa*, *Sorghum bicolor*, *Triticum aestivum*, and *Brachypodium distachyon*. These studies identified SSPs named CAPE (Chien, Nam et al. 2015), CEP (Roberts, Smith et al. 2013, Ogilvie, Imin et al. 2014, Aggarwal, Kumar et al. 2020), CIF (Okuda, Fujita et al. 2020, Fujita 2021), CLE (Goad, Zhu et al. 2017), DVL (Narita, Moore et al. 2004, Wen, Lease et al. 2004), ECL (Sprunck, Rademacher et al. 2012), EPF (Takata, Yokota et al. 2013, Li, Dai et al. 2014, Caine, Chater et al. 2016), ES (Woriedh, Merkl et al. 2015), GASA (Muhammad, Li et al. 2019), IDA (Vie, Najafi et al. 2015), MEG (Xiong, Mei et al. 2014), PIP (Hou, Wang et al. 2014, Zhou, Xiao et al. 2022), PNP (Turek and Gehring 2016), POE (Jiménez-López, Rodríguez García et al. 2011), PSK (Matsubayashi, Ogawa et al. 2006, Stührwohldt, Bühler et al. 2021), PSY (Tost, Kristensen et al. 2021), RALF (Sharma, Hussain et al. 2016, Campbell and Turner 2017, Abarca, Franck et al. 2020), RGF (Fang, Chang et al. 2021), and TPD (Huang, Zhang et al. 2016). Additionally, *Triticum aestivum* sequences for these SSP families were extracted from a large-scale analysis (Tian, Xie et al. 2022).

The published SSP sequences were used to identify sorghum genes in the BTx623 reference genome v3.1.1 (Phytozome v13) using BLASTP analysis. The searches utilized published SSP sequences from *Arabidopsis thaliana* TAIR10, *Zea mays* RefGen_V4, *Oryza sativa* v7.0, *Sorghum bicolor* v3.1.1, *Triticum aestivum* v2.2, and *Brachypodium distachyon* v3.2 as queries. The resulting *Sorghum bicolor* putative SSP encoding genes are reported for CLE (**S-Table 2**), GASA (**S-Table 3**), RALF (**S-Table 4**), and RGF (**S-Table 5**). Among these four families, only the RALF family contained SSPs not previously reported in *Sorghum bicolor*. The nine putative RALF encoding sorghum genes exhibited some regions of sequence similarity (E-value < 1×10⁻⁴) to RALF SSPs identified in other species, but further confirmation will be required to determine if these genes encode functional RALFs. Identifying genes that encode signaling SSPs can be challenging due to their short open reading frames and highly variable central regions (Hu, Lu et al. 2021, Liang, Zhu et al. 2021, Ren, Zhang et al. 2024).

Some SSP families contained conserved domain annotations, which were used to further validate the identity of genes encoding these SSPs. The domain annotations included: RALF (PF05498), EPF (PTHR33109), PSK (PF06404), GASA (PF02704), POE (PF01190), and DVL (PF08137).

To identify conserved motifs, SSP sequences from each family were analyzed using MEME Suite version 5.5.7 in Classic mode. The motif distribution was set to “Zero or One Occurrence Per Sequence” (ZOOPS), and the minimum motif width was fixed at 6 amino acids for all families. For the CLE and RGF families, the maximum motif width was set to 50 amino acids, while for the GASA and RALF families, it was increased to 60 amino acids to accommodate their longer open reading frames and extended SSP lengths. The number of motifs to be identified was set to three for the CLE and RGF families and four for the GASA and RALF families, reflecting the increased sequence complexity of the latter. All input sequences were retained in the MEME output, and the resulting motif images were arranged in tabular format according to phylogenetic order to facilitate visual comparison of motif patterns.

### Phylogenetic Analysis

Full pre-processed SSP sequences and their variants encoded by each sorghum gene described above were collected from BioMart on Phytozome v13. Phylogenetic analysis was conducted separately for each SSP family. Multiple sequence alignments were performed using MUSCLE, and the best-fit protein substitution model was determined using ModelTest-NG v0.1.7 based on the lowest AICc value. All SSP families followed a gamma-distributed model; however, the CAPE, ES, PIP, and PSY families additionally required invariant sites. The WAG model was selected for CAPE, ES, PIP, and PNP; variable-rate models for CEP, CIF, GASA, MEG, POE, and RGF; PMB for CLE, EPF, and RALF; and JTT for DVL, ECL, IDA, PSK, PSY, and TPD. Maximum likelihood phylogenetic trees were inferred using RAxML v8.2.12 with 1,000 bootstrap replicates, running on the Terra and Grace supercomputers at the Texas A&M High Performance Research Computing (HPRC) facility. Trees were visualized and annotated using iTOL v7(Letunic and Bork 2024) and Dendroscope3, enabling interactive exploration and comparative analysis. The best homolog for a sorghum gene encoding a SSP was extracted from the phylogenetic tree based on proximity of a gene encoding a SSP in *Zea mays*, *Oryza sativa*, and *Arabidopsis thaliana* in the phylogenetic tree topology (**S-Table 6-9**). Phylogenetic trees were also generated for *Sorghum bicolor* SSPs independently to determine their relative order and assign gene names accordingly.

### Collection of stem tissues from TX08001 internodes for transcriptome analysis

Sorghum genotype TX08001 was grown in the Automated Precision Phenotyping Greenhouse at Texas A&M University during the fall under long-day conditions until 52 days after emergence (DAE) in the vegetative stage. Plants were cultivated in rhizotrons filled with a mixture of 30% clay and 70% sand. Fertilization was performed using two tablespoons of Osmocote 14-14-14, and watering was provided every 2-3 days. Shoots were harvested in biological quadruplicates, and tissues were collected from phytomers 1-4 and 6-7. Full stem node-internode samples were taken from phytomers 1-4, the youngest phytomers below the shoot apical meristem. For phytomers 6 and 7, which had partially elongated stem internodes, internode tissue samples were collected from the intercalary meristem, located at the base of the internode, to the top of the internode adjacent to the nodal plexus. The pulvinus and nodal plexus were excised separately as 5-10 mm sections, while the intercalary meristem was sampled as a 5 mm section. Additionally, 10 mm sections above the intercalary meristem were collected to capture internode tissues at various stages of elongation and maturation. The upper and oldest portion of each stem internode was divided into 20 mm sections for analysis of later stages of internode maturation. Diagrams showing the pattern of stem dissection are included in **Figure 4b** and **Figure 5b**.

### RNA purification, sequencing and RNA-seq analysis

Methods used for RNA purification and RNA-seq data collection and analysis were the same as described in previous publications (Lamb, Kurtz et al. 2024, Yu, Weers et al. 2025). Tissues were ground to a fine powder with a heat sterilized mortar and pestle filled with liquid nitrogen then transferred into liquid-nitrogen chilled sterile 1.5 mL c centrifuge tubes. RNA was extracted using the Zymo RNA Mini-Prep kit. Purity and the concentration of the RNA was analyzed using a Thermo ScientificTM NanoDrop One Microvolume UV-Vis Spectrophotometer. The integrity of sample RNA was assessed with a Agilent 5300 Fragment Analyzer using software version 3.1.0.12. RNA that passed QC was sent to the Joint Genome Institute for sequencing to a depth of 30–50 million reads. Sequenced reads were aligned to the Sorghum bicolor V3.1 genome using HISAT2 aligner (Kim et al., 2015). Transcriptome assembly and TPM normalization were conducted using StringTie version 1.3 (Pertea et al., 2015). The script prepDE.py https://github.com/gpertea/stringtie/blob/master/prepDE.py and https://ccb.jhu.edu/software/stringtie/index.shtml?t=manual was used to convert nucleotide coverage data from StringTie into read counts that were readable by differential expression statistical packages using the formula: reads_per_transcript = coverage * transcript_length/read_length. Sequence read length was 151 bp. Functional annotations of the transcripts were obtained from the *Sorghum bicolor* V3.1 genome which is available from Phytozome 13 (McCormick, Truong et al. 2018). RNA-seq data was obtained from three biological replicates of all tissues analyzed. Differential gene expression analysis was conducted using the ‘limma’ package in R. TPM normalized transcript data was log2-transformed. A linear model was fitted to the data using the log2-transformed expression values. An empirical Bayes method was applied to the model to assess the standard errors of the estimated log-fold changes. Differentially expressed genes were identified, and the results were adjusted for multiple testing using the False Discovery Rate (FDR) method. Genes with an adjusted p-value < 0.05 were used in the analysis. Fold changes were converted from log2 scale to linear scale for ease of interpretation. TAU analysis was conducted on portions of the data set to provide a gauge of organ and cell type specificity (Yanai, Benjamin et al. 2005). Some figures contain results of analyses of genes expressed at 5 TPM or > in at least one sample of the selected dataset to provide a simplified survey of genes with relatively high expression. Data on all genes was included in Supplemental Tables. Some tables and figures excluded genes that showed differential expression of <5-fold to show patterns of expression focused on genes with higher variation in expression during development. Expression data on SSP-genes with max TPM<5 is included in supplementary tables.

### Analysis of SSP encoding gene expression in plant organs

Some RNA-seq data used in this study was obtained from prior analyses of leaves, stems, roots and panicles of several sorghum genotypes at different stages of developmental (McKinley, Rooney et al. 2016, Kebrom, McKinley et al. 2017, McCormick, Truong et al. 2018, McKinley, Casto et al. 2018, McKinley, Olson et al. 2018, Kebrom, McKinley et al. 2020, Lamb, Weers et al. 2022, Koleva, Liu et al. 2025). This dataset was analyzed to estimate relative organ-level expression of genes that encode SSPs by averaging the expression of genes in tissues that comprise each organ (average of TPM values). The tissues analyzed for each organ were as follows. Leaf organ expression was based on RNA-seq data from the whole leaf, leaf blade, leaf sheath, and leaf whorl. Stem organ expression was based on data from the juvenile stem, stem apex, apical dome, full internode, nodal plexus, pulvinus, and intercalary meristem. Root organ expression was based on RNA-seq analysis of mature roots, elongating roots, meristematic roots, and roots furthest from the stem and deeper in the soil profile. Reproductive stage organ expression was based on RNA-seq data from the peduncle, upper and lower panicle, and seeds.

### Expression of SSP encoding genes in developing stem internodes

RNA-seq data from a prior study of R07020 internode development (Kebrom, McKinley et al. 2017) was used in the current study to characterize the expression of SSP encoding genes in internodes that were beginning to elongate (internode 1), in the process of elongation (internode 2, 3) and a fully elongated internode (internode 4).

### Stem cell type expression of SSP encoding genes

Laser Capture Microdissection (LCM) RNA-seq data from a prior study (Fu, McKinley et al. 2024) was utilized to characterize the expression of genes that encode SSPs in different stem internode cell types of phytomer 8 from the Wray genotype grown to 74 days after emergence (DAE). The selectivity of the expression of genes that encode SSPs in stem cell types was characterized using TAU analysis (Yanai, Benjamin et al. 2005).

## Results

### Identification of sorghum genes that encode SSPs

Sorghum genes that encode SSPs in 19 gene families were identified starting from SSP sequences identified in other plants. Pre-processed SSP sequences were obtained from the genomes of *Arabidopsis thaliana, Brachypodium distachyon*, *Sorghum bicolor*, *Triticum aestivum*, Ze*a mays*, and *Oryza sativa* and used to identify 219 sorghum genes that encode SSPs through homology searches on Phytozome v13 (*Sorghum bicolor*, v3.1.1). Sorghum genes encoding 42 CLEs, 13 GASSs, 23 RALFs, 13 RGFs, 9 CAPEs, 9 CEPs, 4 CIFs, 26 DVLs, 2 ECLs, 11 EPFs, 3 ESs, 4 IDAs, 7 MEGs, 4 PIPs, 3 PNPs, 30 POEs, 6 PSKs, 6 PSYs, and 4 TPD SSPs were identified (**S-Table 1**). Phylogenetic analysis of SSP families was carried out utilizing sequence information from sorghum and the species listed above (**S-Figures 5-23**). In addition, phylogenetic analysis using only sorghum sequences was carried out and the sorghum genes/SSPs in each family were named (numbered) based on their phylogenetic order (**S-Figures 20-37**).

### Sorghum gene families encoding RGF, CLE, GASS, and RALF SSPs

The identification of genes that encode SSPs is often challenging because the conserved sequences that encode the SSPs are short. Therefore, additional information about the SSPs encoded by the sorghum genes predicted to encode RGF, CLE, GASS, and RALF SSPs was collected to help validate the methods used for gene identification.

#### Sorghum RGF/GLV/CLEL gene family

The 13 sorghum genes that encode SbRGF SSPs identified in the current study were annotated in Phytozome and encode proteins that exhibit sequence similarity (E-value < 1 x 10^−16^) to previously published monocot RGFs (Fang, Chang et al. 2021). RGFs play a role in the maintenance and regulation of root meristem cell proliferation capacity (Keerthana, Ramakrishnan et al. 2025). RGFs are typically 13-18 amino acids long and post-translationally modified. RGF SSPs exhibit a conserved DY motif at the N-terminus and an H/(H/N) residue at the C-terminus (Fang, Chang et al. 2021, Shinohara 2021). Motif analysis confirmed the presence of a conserved C-terminal RGF motif and an N-terminal signal peptide sequence in all 13 sorghum RGF SSPs. Sequence analysis of mature RGF SSP sequences confirmed that each sorghum RGF contained a conserved N-terminal DY motif and a C-terminal HH/HN motif (**S-Table 5**). Sobic.006G11200 (*SbRGF3*) is highly related to maize Zm00001d025645, with a bootstrap support of 91(**S-Table 9**).

#### Sorghum GASS SSP gene family

Genes encoding <u>GA</u>-<u>S</u>timulated in <u>S</u>orghum (SbGASS) SSPs, originally reported by (Muhammad, Li et al. 2019), were all annotated in Phytozome and showed sequence similarity to GA-stimulated SSPs in other monocots (E-value < 1 x 10^−34^) (Muhammad, Li et al. 2019). In the *SbGASS* gene family (**S-Table 7**), Sobic.009G119800 (*SbGASS5*) shows a high sequence similarity to the well-characterized maize gene Zm00001d38056 (*ZmGSL1*), with a bootstrap support of 96 (Zhang, Yue et al. 2018). GASA SSPs are characterized by a cysteine-rich motif that stabilizes the SSP’s helix-turn-helix structure. Variation resulting in the loss of cysteine residues impacts structural integrity (Muhammad, Li et al. 2019, Bouteraa, Ben Romdhane et al. 2023, Shang, Ye et al. 2023). Each of the SbGASS SSPs contained two C-terminal cysteine-rich motif regions and a predicted N-terminal signal peptide. Sequence analysis confirmed that each SbGASS SSP contained a strongly conserved cysteine rich region containing 10 well conserved cysteine residues (**S-Table 3**).

#### Sorghum CLE SSP gene family

Forty two *Sorghum bicolor* genes were previously reported to encode CLE (SbCLE) SSPs (Goad, Zhu et al. 2017) although only *28 SbCLE* genes are annotated in Phytozome 13. All but one of the 42 *SbCLEs* exhibited sequence similarity to monocot CLE-encoding genes (E-value< 1 x 10^−5^) as previously reported (Goad, Zhu et al. 2017, Tian, Xie et al. 2022). *SbCLE17* (Sobic.002G361800) was not identified in maize or rice suggesting that this sorghum gene is highly diverged and possibly non-functional. Sobic.004G041700 (*SbCLE32*) is closely related to maize *ZmCLE18* (Zm00001d015282), with a bootstrap support of 92 (Je, Gruel et al. 2016) (**S-Table 6**).

Mature CLE/CLV SSPs are less than 20 amino acids in length and contain numerous post-translational modifications (Keerthana, Ramakrishnan et al. 2025). Pre-processed CLE proteins contain an N-terminal signal peptide and a C-terminal signaling peptide that is cleaved off for biological activity. All SbCLE proteins except SbCLE29 possessed a predicted N-terminal peptide sequence (**S-Table 2**). Biologically active CLE SSPs typically contain a dibasic KR motif, an N-terminal H/N and central prolines or glycines (Fiers, Golemiec et al. 2005, Ito, Nakanomyo et al. 2006, Ohyama, Shinohara et al. 2009, Katsir, Davies et al. 2011). Motif analysis confirmed that each SbCLE SSP harbored a single conserved C-terminal CLE motif whereas SbCLE37, SbCLE38, and SbCLE40, contained multiple CLE motifs within their pre-processed sequences. Sequence analysis showed that each mature CLE SSP contained an N-terminal R/K except for SbCLE16, 39, 41, and 32. The SSPs with multiple CLE motifs were not analyzed further since it is not clear which sequence represents the mature peptide sequence. All mature SbCLE SSPs contained a conserved P-X-G-P-X-P central motif or related sequence. All SSPs also contained the C-terminal H/N on the mature peptide except for SbCLE16 (**S-Table 2**).

#### Sorghum RALF SSP gene family

Sixteen of the 23 sorghum genes that encode RALFs identified in this study were previously identified by (Campbell and Turner 2017) and 18 were annotated in Phytozome. Of these, Sobic.003G415500 (*SbRALF6*) shares a high sequence similarity with the maize gene Zm00001d011921 (*ZmRALF10*) (**S-Table 8**), with a bootstrap support of 98 (Zhou, Wang et al. 2024). SbRALF genes that were not annotated in Phytozome exhibited some sequence similarity to previously published monocot RALF SSPs that lacked or contained modified RALF motifs (Sharma, Hussain et al. 2016, Tian, Xie et al. 2022). These genes/SSPs were labeled RALF-related (RALFR#). All of the SbRALFs exhibited homology (E-value <10^−6^) to previously identified monocot RALFs (Sharma, Hussain et al. 2016, Campbell and Turner 2017).

RALF SSPs possess the conserved YISY motif and a dibasic RR cleavage site. Sequence variation is mainly observed in the YISY motif and cleavage site, which are conserved between monocots and dicots (Matos, Fiori et al. 2008, Srivastava, Liu et al. 2009, Sharma, Hussain et al. 2016, Abarca, Franck et al. 2020). Motif and sequence analysis revealed that SbRALFR1, SbRALFR2, SbRALF3, 5, 6, 7, 8, 9, 11, 12, 13, 14, SbRALFR17, SbRALFR18, and SbRALF21 each had an identifiable RRXL cleavage site whereas the other RALF or RALFR SSPs did not contain an easily identifiable cleavage site. SbRALFR2, SbRALF3, 4, 5, 6, 7, 9, 10, 11, 12, 13, 14, RALFR17, SbRALFR20, and SbRALFR23 all contained a YISY motif or similar sequence (i.e. SIGY) (**S-Table 4**).

SbRALFR1, 2, and 17-23 displayed divergent motifs compared to canonical RALFs. SbRALFR22 showed sequence similarity to SbRALF20, however, SbRALFR22 lacked recognizable RALF motifs and is not homologous to any previously identified RALF, suggesting it may represent a different class of SSP. Our results are consistent with a prior study that identified a divergent class of clade IV RALF SSPs that lack the YISY motif and RRXL cleavage site, however these retain some sequence similarity to other RALF SSPs within their cystine rich region (Campbell and Turner 2017). Functional characterization will be required to determine if these SSPs function as RALFs (**S-Table 4**).

### Expression of SSP encoding genes in sorghum organs

Expression of the sorghum genes that encode SSPs in leaves, stems, roots and panicles was examined using RNA-seq data derived from those organs/tissues (**S-Table 10**). The expression of the 219 genes that encode the 19 families of SSPs in sorghum organs is summarized in **S-Figure 38**. The average expression of each gene in the four organs was used to calculate TAU values, a gauge of tissue specificity, with values closer to 1 indicating high tissue specificity (Yanai, Benjamin et al. 2005). Forty three of the 219 sorghum genes that encode SSPs were expressed in an organ specific manner (Tau >0.95). The relative expression of sorghum genes expressed at >5 TPM that encode SbCLE, SbGASS, and SbRALF genes in leaves, stems, roots and panicles is shown in **Figure 1a**. A max TPM >5 across the multiple tissue samples was used for average calculation across an organ and TAU calculation on organ averages to identify SSPs with relatively high expression in each organ type and determine their organ specificity. 158 SSP encoding genes had max TPM>5. The other 61 SSPs with max TPM<5 in each tissue sample is also included in (**S-Table 10**). While these gene families are expressed in all four organs, some genes show relatively high organ specific expression whereas others are expressed in more than one organ. For example, *SbCLE1*, *3*, *6*, *7*, *9*, *10*, *11*, *12*, *15*, *24*, 25, and *26* exhibited relatively high expression in roots whereas *SbCLE31* displayed relatively high expression in leaves and *SbCLE23* showed relatively high expression in the stem. *SbCLE37* and 38 were specifically expressed in the panicle, while *SbCLE18*, *20*, *23*, *27*, *28*, *41*, and *42* showed relatively high expression in the stem. *SbCLE18*, *20*, *24*, *25*, *26*, and *27* also had relatively high expression in the panicle. Among the sorghum genes encoding ‘GA-stimulated in sorghum’ SSPs (*SbGASS*), *SbGASS3*, *4*, *5*, *6*, *7*, *8*, *9*, and *13* exhibited relatively high expression in stems and *SbGASS2*, *4*, *6* were expressed in leaves. *SbGASS2* was also expressed in roots and *SbGASS1*, 2, 3, 4, 5, 7, 9, *10*, *11, 12, and 13* were expressed in the panicle. *SbRALFR1*, *2, 17*, *19*, and *23* and *SbRALF5, 7*, *9*, *11*, *12*, were expressed in the stem, while *SbRALFR2, 21*, and *23* and *SbRALF9* were expressed in the root. Several *RALF* genes, including *SbRALFR1*, *20*, and *22* were expressed in the leaf, whereas *SbRALF6*, *10*, *13*, *14*, *15*, and *16* exhibited relatively high expression in the panicle. Expression of *SbRALF5*, *7*, 9 *11*, *12* and *SbRALFR1, 2, 17, 19, 22, and 23* appeared to be the most stem-specific, while *SbRALF21* was root-specific (τ = 0.92).

**Figure 1.**
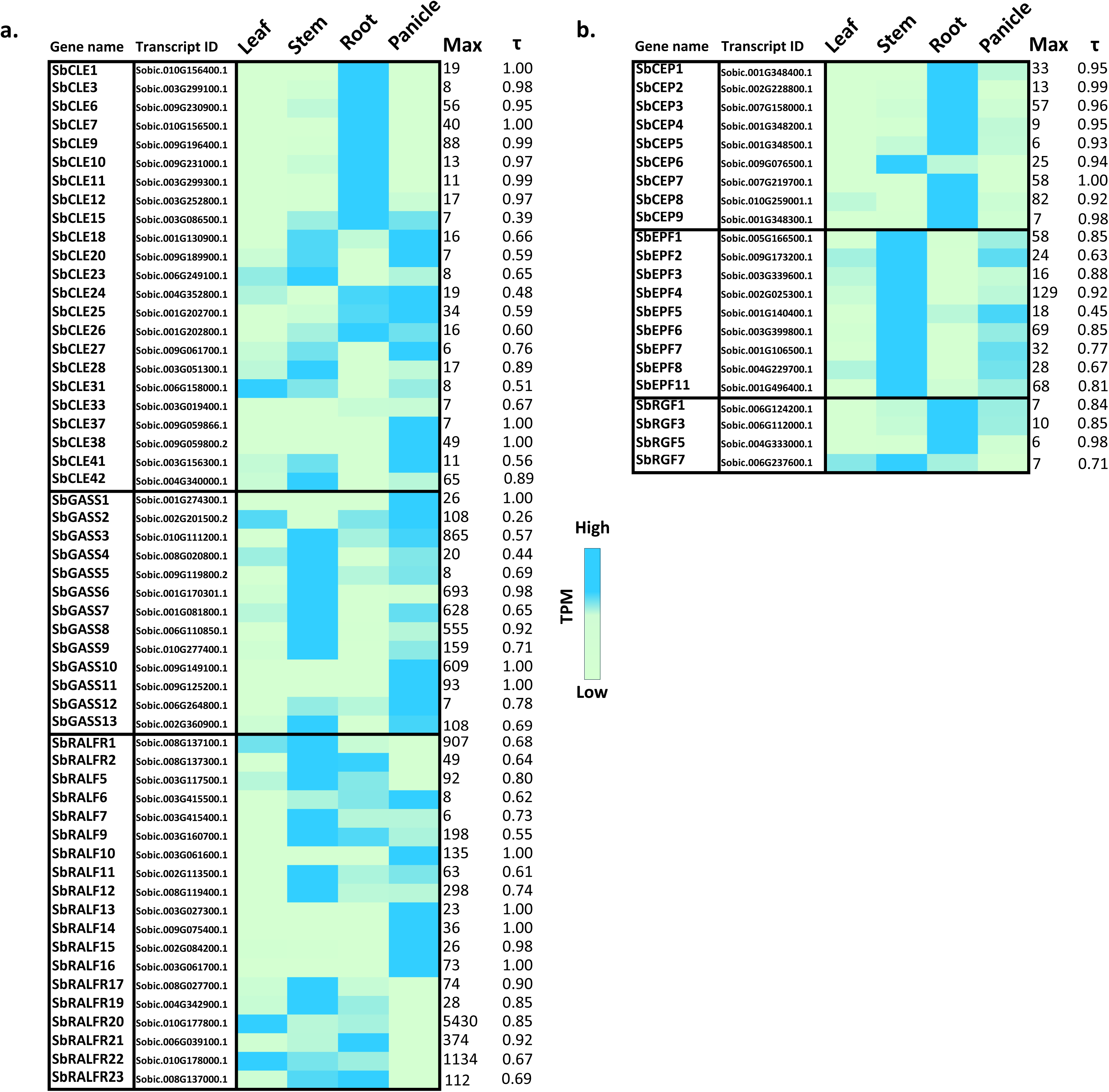
Expression of genes that encode SSPs in sorghum leaf, stem, root, and panicle tissues. Genes were ordered within families based on phylogenetic analysis. Gene encoding SSPs shown were expressed with an average >5 TPM in at least one organ. Expression values are reported in TPM, with blue indicating high expression and green indicating low expression. a) Expression of genes encoding CLE, GASS, and RALF peptide families, b) Expression of genes encoding CEP, EPF, and RGF peptide families that show elevated organ specific expression. The Max column is the highest TPM expression of the four organs. The τ column is of the gauge of organ specificity.

The *RGF* and *CEP* gene families were enriched in genes expressed in roots consistent with prior studies of RGF and CEP SSP function (Keerthana, Ramakrishnan et al. 2025). For example, *SbRGF1*, *3*, and *5* were more highly expressed in the roots than leaves, stems and panicles however, *SbRGF7* was expressed in stems and leaves (**Figure 1b**). Similarly, eight of the nine *SbCEP* genes were highly expressed in roots compared to leaves, stems and panicles (**Figure 1b**). In contrast, the *SbEPF* genes were highly expressed in stems, and some family members were also expressed at somewhat lower levels in panicles, possibly because panicles contain stem-like structures (i.e., rachis) (**Figure 1b**).

### Stem cell type expression of genes that encode SSPs

Expression of sorghum genes in stem internode cell types was characterized using data generated previously using laser capture microdissection (Fu, McKinley et al. 2024). The stem tissues/cell types targeted for transcriptome profiling included the epidermis, pith parenchyma cells, xylem fibers, vascular parenchyma, and phloem (**Figure 2**). SSP encoding genes expressed with high tissue/cell type specificity were identified by Tau analysis (<0.9) and sorted by cell type (**Figure 2a**). Tau analysis identified numerous genes with highly specific expression in the epidermis, pith parenchyma, vascular parenchyma or phloem (**Figure 2a**). Specific *CLE* and *PSK* gene family members were expressed at high levels and specificity in each of the four cell types (**Figure 2a**). In contrast, two IDA genes were expressed at high levels only in the epidermis, while *SbEPF1* was highly expressed in the epidermis and S*bEPF4* was highly expressed in stem pith parenchyma cells (**Figure 2a**). Three *SbRALF* genes were expressed with high cell type specificity in stem vascular parenchyma (**Figure 2a**).

**Figure 2.**
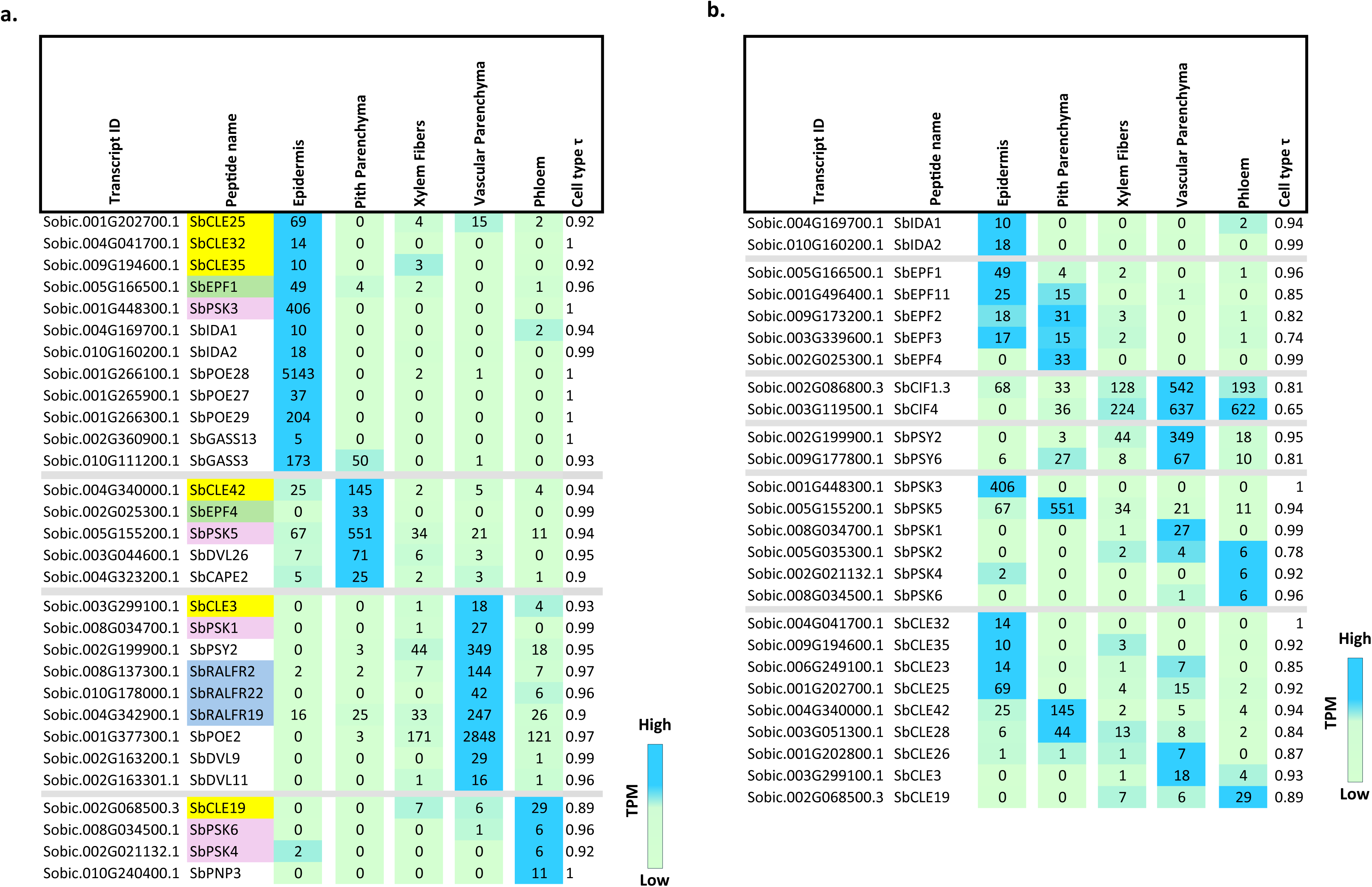
Expression of genes that encode SSPs in cell types of a fully elongated internode of sorghum genotype Wray. a) Peptide encoding genes that exhibit high cell type specific expression (τ>0.89), b) Expression of all SSP encoding genes expressed at >5 TPM in at least one stem internode cell type, sorted by peptide family.

Analysis of all SSP encoding genes expressed at >5 TPM in fully expanded stem internode tissue showed that five genes encoding EPFs were highly expressed in the epidermis, pith parenchyma or both the epidermis and pith (**Figure 2b**). Two *SbCIF* genes were more highly expressed in cells of vascular bundles (xylem fibers, vascular parenchyma, phloem) and two *SbPSY* genes were highly expressed in vascular parenchyma (**Figure 2b**). In contrast, the six members of the *SbPSK* gene family were expressed selectively in the epidermis, pith parenchyma, vascular parenchyma or phloem (**Figure 2b**). Several *SbCLE* genes were expressed in the epidermis and other members of that gene family were expressed in the epidermis, pith parenchyma, vascular parenchyma and phloem. For example, *SbCLE42*, a homolog of TDIF, was highly expressed in pith parenchyma. *SbCLE3* was highly expressed in vascular parenchyma, with additional expression in the phloem and xylem fibers. *SbCLE19* was most highly expressed in the phloem, with additional expression in vascular parenchyma and xylem fibers (**Figure 2b**).

Some SSP encoding genes were broadly expressed across multiple cell types. For example, *SbRALFR1* was expressed at 1499 TPM in the epidermis and 917 TPM in the core pith/parenchyma, also exhibited high expression in xylem fibers, vascular parenchyma, and phloem (**S-Table 13**). *SbRALF11* was also expressed in the epidermis, core pith/parenchyma, xylem fibers, vascular parenchyma, and phloem (**S-Table 13**). As noted above, *SbRALFR2*, *SbRALFR19 and SbRALF22* were predominantly expressed in the vascular parenchyma, with low expression in the epidermis, core pith/parenchyma, xylem fibers, and phloem. *SbRALFR17* was expressed in both the epidermis and pith/parenchyma (**S-Table 13**). *SbRALFR21* showed peak expression in the phloem and vascular parenchyma, with intermediate levels in the xylem fibers and low expression in the epidermis and core pith-parenchyma. Similarly, *SbRALFR20* was highly expressed in vascular parenchyma and phloem, with moderate expression in xylem fibers (**S-Table 13**).

With respect to the other genes encoding SSPs (**S-Table 13**), *SbGASS13* showed relatively specific expression in the epidermis whereas *SbGASS3* and *SbGASS8* showed relatively high expression levels in the epidermis and pith-parenchyma cells (**S-Table 13**). In addition, *SbPOE28*, *SbPOE29*, *SbPOE27*, and *SbPOE11* are expressed in the stem epidermis whereas *SbCAPE2*, *SbDVL22*, and *SbDVL26* are expressed in stem core pith/parenchyma. *SbCIF1.3*, *SbDVL9*, *SbDVL11*, *SbPOE14*, *SbPOE1*, *SbPOE2* have relatively high expression in the stem vascular parenchyma along with some expression in xylem fibers and phloem. *SbMEG5* showed high expression in the stem phloem cells. *SbPOE3* and *SbCIF4* are expressed in xylem fibers, stem vascular parenchyma, and stem phloem. *SbDVL25* and *SbPOE21* are expressed in each of the cell types analyzed.

### Expression of SSP encoding genes in developing stem internodes

The potential involvement of SSPs in stem growth regulation was investigated by analyzing the expression of SSP encoding genes during stem internode development. In a previous study, transcriptome profiles were collected from four stages of R07020 stem internode growth and development (Kebrom, McKinley et al. 2017). Genes involved in cell proliferation (i.e., *SbSDKB2*) were highly expressed in the youngest internode (INT1) and genes involved in lignin synthesis and secondary cell wall formation (i.e., *SbCESA4*) were highly expressed in a fully elongated internode (INT4) (Kebrom, McKinley et al. 2017). In the current study, expression of genes that encode SSPs in INT1-4 was investigated to identify genes that are differentially expressed during this phase of internode growth and development. The analysis identified genes that encode SSPs that were highly expressed early in internode development (INT1) that showed decreasing expression during development and in fully elongated internodes (INT4) (INT1>4), whereas other genes were expressed at low levels in INT1 and higher levels during development and in INT4 (INT4>1) suggesting they may play roles in internode differentiation following cessation of growth (**Figure 3**). For example, *SbGASS8*, *SbGASS9*, *SbGASS3*, *SbGASS13*, and *SbGASS2* showed a INT1>4 pattern of expression during internode development. Similarly, *SbRALF7, SbCAPE9*, *SbDVL24, SbEPF7/8, SbES1, SbPOE19* and *SbPSY3* exhibited an INT1>4 expression pattern, suggesting a potential role in regulating growth processes during early internode development. Conversely, several members of the RALF SSP family, including *SbRALF5*, *SbRALFR2*, *SbRALFR20*, *SbRALFR22* and *SbRALFR19* displayed an INT4>1 pattern of expression highlighting their possible involvement in processes associated with internode differentiation and secondary cell wall formation. Expression of *SbCIF1*, *SbPIP2*, *SbPOE2/14/27/28/29* and *SbPSK3/5* also increased during internode development (INT4>1) (**Figure 3**). SSPs which were expressed more than 5 TPM in at least one internode but did not have a FC > 5 are included in **S-Table 11** along with SSPs which were expressed less than 5 TPM.

**Figure 3.**
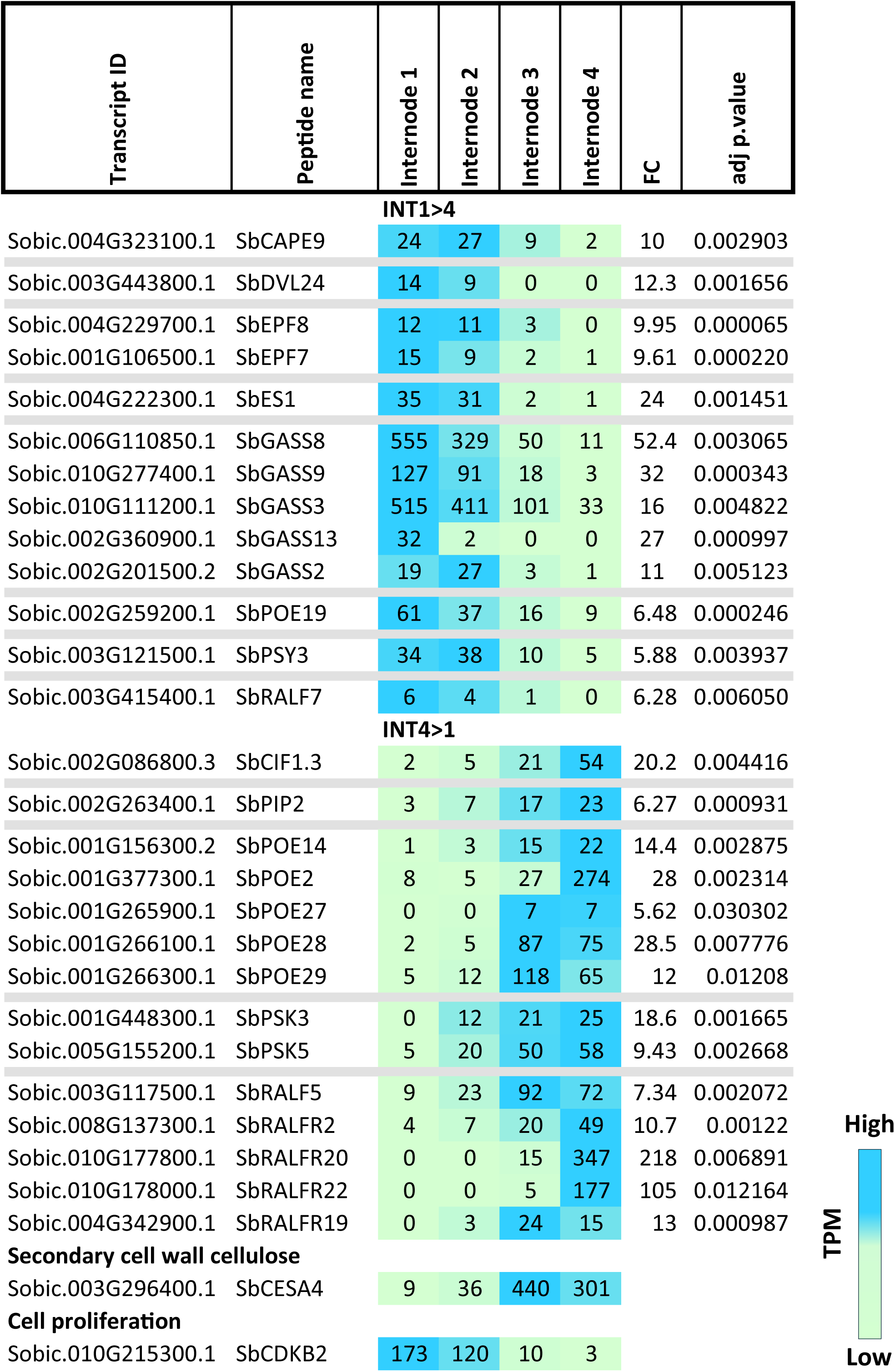
Expression of genes that encode SSPs during development of internodes of sorghum genotype R07020. The youngest internodes (INT1/2) showed elevated expression of *SbCDK2*, a marker gene for cell proliferation. Older internodes (INT3/4) showed elevated expression of *SbCESA4*. Genes were sorted based on their differential expression in young (INT1) versus older internodes (INT1>4 or INT4>1). Genes included in this figure had a fold change (FC) > 5 (adjusted p-value < 0.05).

### Expression of SSP encoding genes during TX08001 stem development

Sorghum stems are comprised of node-internode segments that part of phytomers initially formed below the stem apical meristem approximately every 3-4 days during vegetative growth. The node-internode segments of each phytomer are comprised of the nodal plexus, where the leaf sheath joins the stem, internode tissue produced by an intercalary meristem, and the pulvinus where the formation of tiller buds and nodal root buds takes place. An intercalary meristem (IM) zone of cell division (ZoD) is located at the base of a growing internode just above the pulvinus, with an internode zone of cell elongation (ZoE) located immediately above the IM and a zone of cell maturation (ZoM) above the ZoE (Yu, Oliver et al. 2022) (**Figure 4b, 5b**). To better understand the potential role of SSPs in stem growth and development, the expression of genes that encode SSPs in stem tissues during growth and development was investigated. Stem tissue associated with phytomers 1–4 (Phy1-4) corresponded to nascent nodes that were harvested without further dissection. Stem node-internode segments of phytomers 6 and 7 were dissected into the nodal plexus (NP), internode, intercalary meristem (IM), and pulvinus for RNA-seq analysis. Elongating/elongated internode tissue was divided into 5-, 10-, and 20-mm sections starting from the base of the internode (5 mm, IM) to the nodal plexus (20 mm) (**Figure 4b and 5b**). SSPs that were classified to be regulating growing internodes were sorted based on their similarity in expression to a cell proliferation gene Sobic.010G215300.1 (*SbCDKB2*) used as a biomarker. For SSPs that were classified for regulating stem tissue differentiation and secondary cell wall formation, Sobic.003G296400 (*SbCESA4*) served as a secondary cell wall biomarker for pattern grouping.

**Figure 4.**
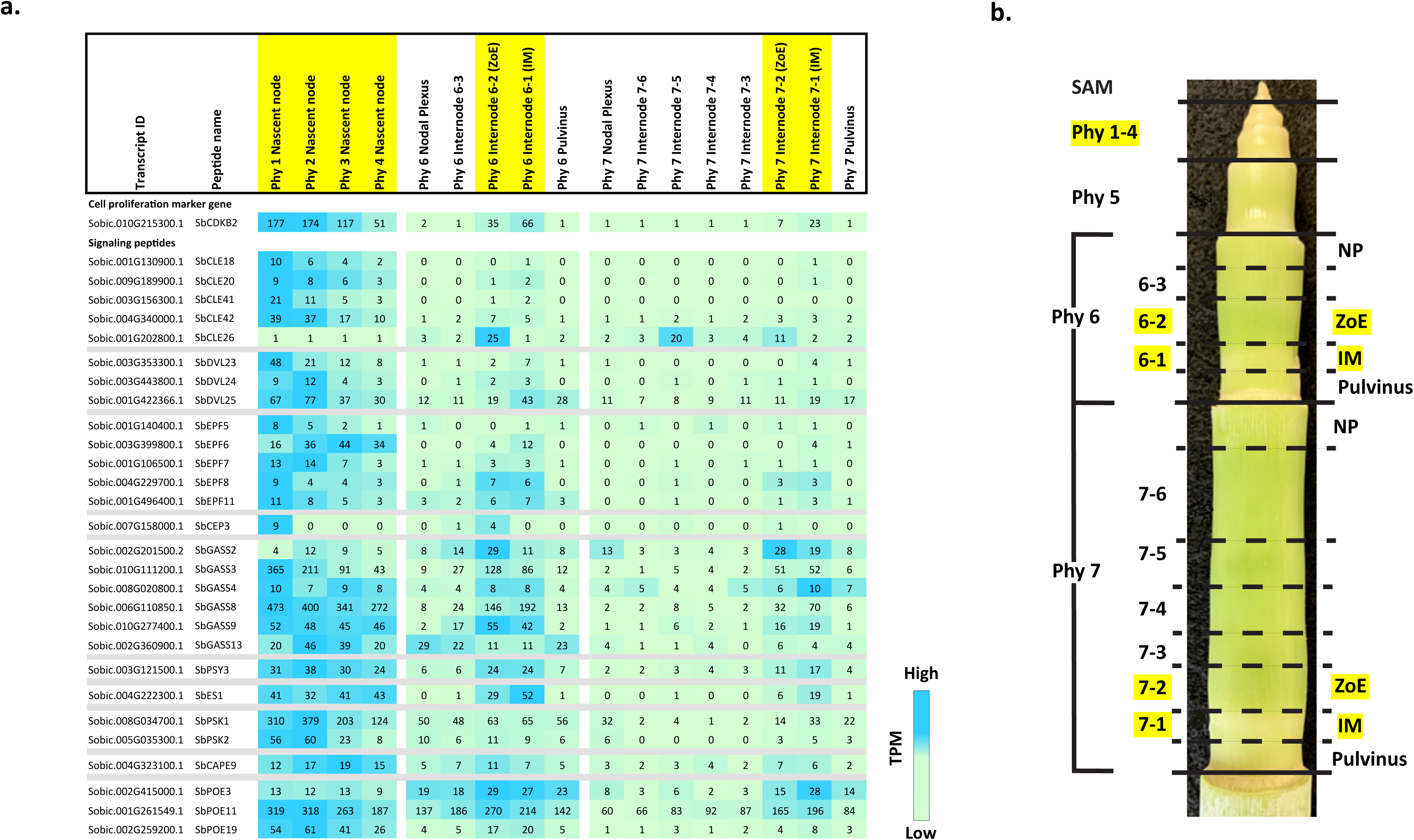
Expression of genes encoding SSPs that are highly expressed in sorghum stem growing zones. a) Heat map of peptide gene expression in stems of sorghum genotype TX08001. Stem tissues with active cell proliferation (highlighted in yellow) were identified using *SbCDKB2* as a cell proliferation biomarker. Nascent nodes of phytomers 1-4 (Phy1-4) and the internode intercalary meristem (IM) and adjacent zone of elongation (ZoE) of phytomers 6 and 7 were identified as regions of cell proliferation. b) Stem tissue identification based on the analysis of (Yu, Oliver et al. 2022). Stem tissues collected for RNA-seq analysis included apical tissue spanning the SAM (shoot apical meristem), IM (intercalary meristem), ZoE (Zone of Elongation), pulvinus and nodal plexus (NP). Nascent nodes were collected from Phy1-4. Stem internode sections 6-1 and 7-1 were 5 mm long, whereas 6-2 and 7-2 were 10 mm sections.

Expression of *SbCDK2,* a gene associated with high rates of cell proliferation and *SbCESA4*, a gene involved in secondary cell wall formation were used as a biomarkers for tissue/cells that are proliferating or differentiating, respectively (Yu, Oliver et al. 2022). *SbCDK2* was highly expressed in stem tissue associated with the youngest internodes of phytomers 1-4 and the intercalary meristem whereas *SbCESA4* expression was highest in the nodal plexus of phytomer 6 and nodes-internodes of phytomer 7 (**Figure 4a and 5a**). Expression of the 219 sorghum genes that encode SSPs was analyzed to identify those expressed at 5 TPM or greater levels in this set of stem tissues and those that show variation in expression during development. Genes encoding SSPs less than 5 TPM are included in (**S-Table 12**). A figure showing the relative expression of genes encoding all families of SSPs in this analysis was generated to illustrate the complexity and dynamics of SSP gene expression during stem development (**Figure 4a**). Genes encoding SSPs expressed in stem tissue from the youngest phytomers (1–4) but at low levels in older phytomers (6–7) suggest they play specific roles in early stages of stem node development (**Figure 4a**). Examples of such SSPs include *SbCLE18*, *SbCLE41*, and *SbCLE42*. Expression of *SbGASS8*, *SbGASS3*, *SbGASS9*, and *SbGASS4* was elevated in apical and intercalary meristem tissues that are characterized by high rates of cell proliferation (Yu, Oliver et al. 2022). SSP encoding genes such as *SbGASS7*, *SbGASS6*, and *SbGASS13* were also highly expressed in the stem apical phytomers and lower expression in stem tissues from phytomer 7. These SSP encoding genes were expressed several tissues indicating they may regulate broader developmental processes.

**Figure 5.**
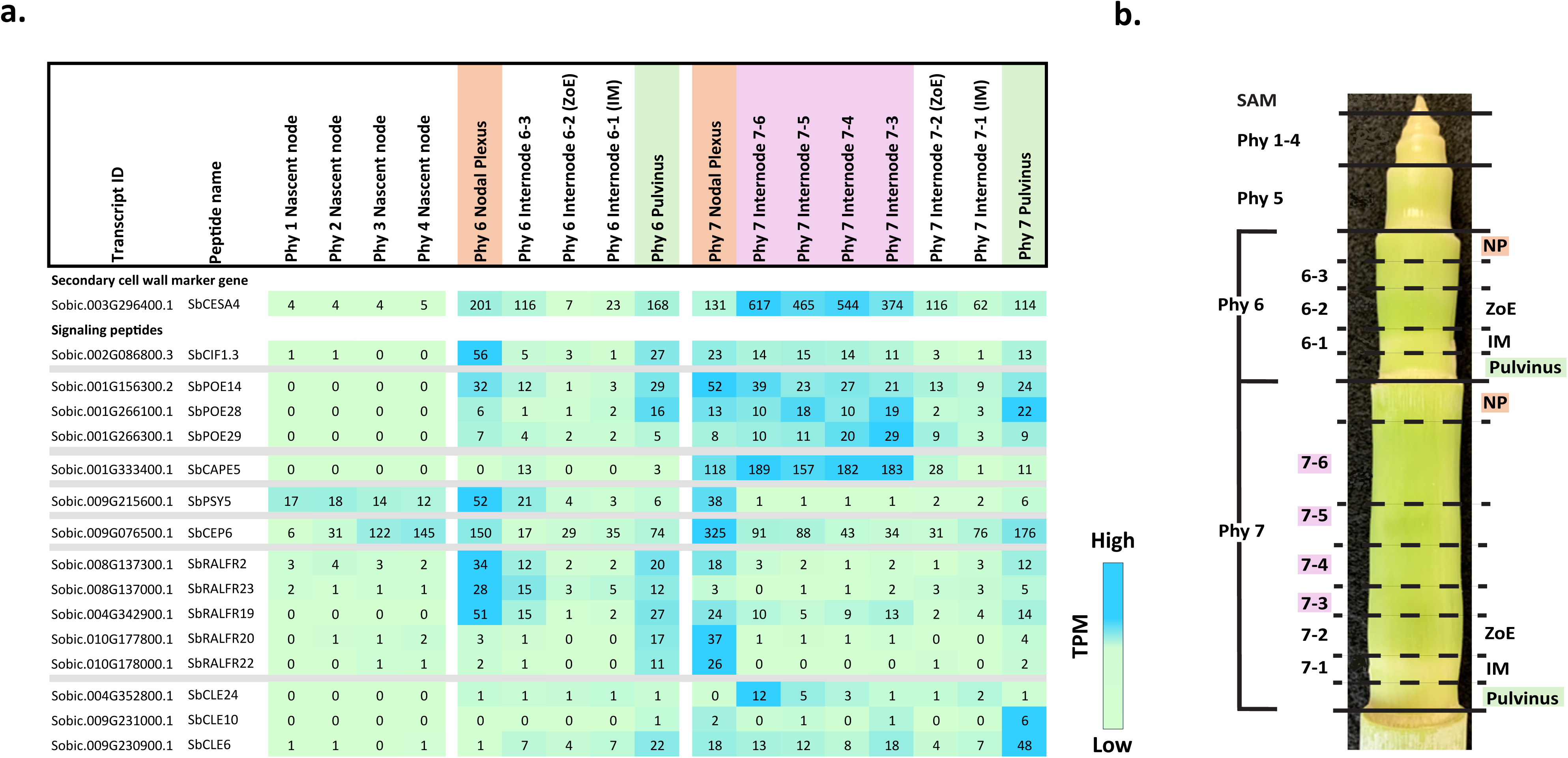
Expression of genes that encode SSPs in sorghum stem differentiating zones. a) Heat map of peptide encoding genes that show elevated expression during stem tissue differentiation of sorghum genotype TX08001. Regions of cell differentiation were identified using *SbCESA4* as a secondary cell wall formation biomarker. The orange highlight was used to denote stem nodal plexus (NP) tissue whereas green highlight was used to denote stem pulvinus tissue. The purple highlight corresponds to tissues within the internode that have completed the elongation process and expressing genes involved in secondary cell wall synthesis. b) Stem tissue identification based on the analysis of (Yu, Oliver et al. 2022). NP refers to Nodal Plexus. The sections corresponding to 6-3 and 7-3 were 5 mm long, 7-4 and 7-5 were 10 mm long, and section 7-6 was 20 mm in length.

*SbCLE26* and *SbGASS2* genes encoding SSPs presented an intriguing expression pattern, with elevated expression in a 10 mm section of the internode just above the IM of phytomer 6. At this stage of internode tissue development, rates of cell proliferation decrease coinciding with increased cell elongation followed by cessation of growth and synthesis of secondary cell walls. Expression of *SbCESA4*, a biomarker associated with secondary cell wall formation (McKinley, Rooney et al. 2016), was highest in the upper regions of later internodes that had completed elongation, suggesting the 10 mm section of phytomer 6 above the IM represents a transitional zone where cell elongation increases. This correlation suggests SbCLE26 and SbGASS2 may regulate the decrease in cell proliferation or increase in cell elongation.

*SbCLE24* was highly expressed in internodes undergoing secondary cell wall formation, while *SbCLE10* and *SbCLE6* were predominantly expressed in the pulvinus and nodal plexus of older phytomers (**Figure 5a**). *SbGASS12* was uniquely expressed in the internode and pulvinus of phytomer 6 (**S-Table 12**). Members of the SbRALF family, including *SbRALFR23*, *SbRALFR2*, *SbRALFR19*, *SbRALFR20*, and *SbRALFR23*, were expressed more highly in the nodal plexus and upper internodes of phytomers 6 and 7, suggesting a role in nodal plexus development (**Figure 5a**).

The other genes encoding SSPs like *SbDVL24*, *SbDVL23*, *SbEPF7*, and *SbEPF9* exhibited elevated expression in young stem tissue of phytomer 1-4 (**Figure 4a**). *SbEPF8*, *SbEPF11*, *SbEPF6*, *SbES1*, *SbPOE19*, *SbPSK2*, *SbPSY3* were expressed in meristematic tissues of the internode. *SbCAPE9*, one variant of *SbCIF1.3*, *SbDVL25*, *SbDVL1*, *SbEPF4*, *SbPOE11*, *SbPSK1* were expressed at higher levels during development in all the internode tissues. *SbCAPE5*, a variant of *SbCIF1.2*, *SbEPF1*, *SbIDA1*, *SbPOE3*, *SbPOE26*, *SbPOE6*, *SbPOE14*, *SbPOE28*, *SbPOE29*, and *SbPSY5* show a pattern of increased expression during stem development, with interesting patterns in the nodal plexus tissue. *SbCEP6*, *SbCIF4*, *SbDVL22*, *SbDVL26*, *SbDVL21*, *SbPOE23*, *SbPOE2*, *SbPOE21*, *SbPSK3*, *SbPSK5*, *SbPSK6*, *SbPSY2* showed elevated expression in many of the mature tissues (**Figure 4a**, **Figure 5a, and S-Table 12**).

## Discussion

SSPs regulate plant growth, differentiation, adaptation to abiotic stress, and responses to plant pathogens and pests (Czyzewicz, Yue et al. 2013, Matsubayashi 2014, Xie, Zhao et al. 2022, Xiao, Zhou et al. 2025). The action of SSPs is complementary to and integrated with regulation of similar processes by the plant hormones ABA, IAA, CK, GA, JA and ethylene. While the role of plant hormones in sorghum stem growth regulation has been investigated (i.e., (Multani, Briggs et al. 2003, Hilley, Truong et al. 2016, Kebrom, McKinley et al. 2017, Yu, McKinley et al. 2021) the function of SSPs is largely unexplored. Therefore, the goal of this study was to update the identification of sorghum genes that encode SSPs and analyze the expression of these genes in sorghum organs, stem cell types and during stem growth and development. The current study identified 219 sorghum genes that encode SSPs that were members of 19 SSP families. Some of the sorghum genes were identified previously in a study of rice SSP encoding genes (Muhammad, Li et al. 2019) based on sorghum genome sequence V1 (Paterson, Bowers et al. 2009). The current study annotated sorghum genes that encode SSPs encoded in the BTx623 reference genome sequence V3.1.1 (McCormick, Truong et al. 2018) based on homology to SSP encoding genes identified in other plant species, motif identification, phylogenetic analysis, and expression profiling.

### Organ and cell-specific expression

Sorghum genes that encode SSPs were expressed in leaves, stems, roots and panicles. Several gene families were mostly expressed in one organ. For example, genes encoding SbCEP and SbRGF SSPs were largely expressed in roots, whereas genes encoding EPF SSPs were primarily expressed in stems and panicles. Specific genes encoding members of other SSP families (i.e., *SbCLE, SbGASS* and *SbRALF*-genes) were expressed in different organs, or in some cases, more than one organ. Expression of sorghum genes that encode SSPs in stem cell types was examined using LCM-derived cell specific transcriptome data from a fully expanded stem internode (Fu, McKinley et al. 2024). That analysis showed that many of the sorghum SSP encoding genes are expressed in a cell type specific manner (Tau >0.9). One gene family was selectively expressed in cells of the stem epidermis (i.e., IDAs) while genes encoding EPF SSPs were expressed in the epidermis and pith parenchyma. Specific members of other gene families (i.e., *CLE, PSK*) were expressed in each of the stem cell types analyzed (epidermis, pith, vascular parenchyma, phloem). Overall, the analysis showed that expression of many of the SSP encoding genes in sorghum occurs in a highly organ and cell-type specific manner that would enable targeted spatial regulation of cell type growth and differentiation.

### Expression of SSP genes during stem development

Bioenergy sorghum stems account for ∼80% of harvested biomass at the end of the growing season (Olson, Ritter et al. 2012). Sorghum stem architecture has an impact on leaf distribution in the canopy, stems provide a conduit for water, sugars and nutrient exchange between roots and leaves, and stem anatomy and composition influences stalk lodging and conversion of biomass to bioproducts. Sorghum stem growth and development is highly regulated and influenced by genotype (Multani, Briggs et al. 2003, Hilley, Truong et al. 2016, Hilley, Weers et al. 2017), environmental factors (i.e., shading) (Yu, McKinley et al. 2021) and mechanical stress (Zargar, Li et al. 2022).

The current study found that numerous sorghum SSP encoding genes are differentially expressed in stem zones of cell proliferation (apical, IM) or at later stages of stem tissue development. For example, thirteen genes were differentially expressed early in internode development during the cell proliferation stage, and a different set of 14 genes were induced later in internode development when cells are differentiating (Fig 2). Analysis of stem development starting with nascent nodes located in stem apex (phytomers 1-4) and extending through stem tissues in phytomers 6 and 7 revealed additional complexity of SSP gene expression. For example, several *SbCLE*-genes were expressed at their highest levels in nascent nodes of phytomer 1 (*SbCLE18, 20, 41, 42*) followed by decreased in expression in stem tissue of older phytomers. Other genes were highly expressed in the apical nascent nodes and in the internode IM and adjacent ZoE (i.e., *SbGASS3, SbPSY3*) indicating a more general role in the regulation of cell proliferation and enlargement consistent with prior studies (Yu, Oliver et al. 2022, Zhang, Han et al. 2025). Other genes were expressed at low levels in nascent nodes and at higher levels in the nodal plexus and fully elongated portions of internodes in older phytomers (i.e., *SbPOE14, 28, 29, SbCAPE5, SbRALFR2,19, 20, 22, 23*).

Phylogenetic analysis was used to identify homologs of sorghum SSP encoding genes in Arabidopsis, rice and maize where many studies of SSPs have been carried out. Information derived from research in other plants was used to help interpret the patterns of expression observed in sorghum.

### SbGASS genes

Homologs of SSP encoding *GASA* (GA-Stimulated in Arabidopsis) genes are found in many plant species including sorghum (Muhammad, Li et al. 2019). *GASA-*homologs are involved in growth regulation (i.e., (Sun, Wang et al. 2013, Han, Jin et al. 2017, Lee, Kim et al. 2017), salt tolerance (Lee, Han et al. 2015), regulation of ROS levels (Rubinovich and Weiss 2010) and regulation of flowering and seed development (Roxrud, Lid et al. 2007). Three sorghum *GASS* genes were previously found to be differentially expressed in the sorghum stem intercalary meristem (Yu, Oliver et al. 2022) consistent with GA-regulation of cell proliferation in stem internodes (Yu, McKinley et al. 2021). The current study showed that five *SbGASS*-genes (*SbGASS2,3,4,8,9,13*) were expressed in nascent stem nodes of phytomers 1-4 and/or in the internode intercalary meristem consistent with a role in GA-modulated cell proliferation. *SbGASS3* and *SbGASS13* were also expressed in the epidermis of a fully elongated section of the internode of phytomer 7.

### SbCLE genes

In the current study, *SbCLE18, 20, 41*, and *42* were highly expressed in nascent stem nodes *SbCLE26* was expressed in the internode zone of cell elongation, and *SbCLE6, 10* and *24* were expressed in the pulvinus or internode tissue later in stem development. *SbCLE41* and *SbCLE42* are homologs of TDIFs that promote expression of *WOX4* and vascular cambium activity and inhibit xylem differentiation (Qiang, Wu et al. 2013). *SbCLE42* is a homolog of *ZmCLE2-1*, which interacts with WUS2 to regulate meristem activity (Dong, Shi et al. 2023). Further analysis of the *SbCLE*-genes that are expressed in apical nascent stem nodes may provide insight into the early stages of sorghum stem growth and vascular bundle cell development. Analysis of stem internode cell type expression showed that *SbCLE42* was selectively expressed in pith parenchyma of a fully elongated portion of the internode associated with phytomer 7. Continued expression of *SbCLE42* in pith cells that lack secondary cell walls is consistent with TDIF’s additional role in repressing xylem cell differentiation and secondary cell wall formation (Qiang, Wu et al. 2013).

### SbRALF genes

RALF SSPs have been found to regulate plant growth, root hair elongation, pollen development, and plant responses to adverse environmental conditions (Murphy and De Smet 2014, Blackburn, Haruta et al. 2020, Liu, Liu et al. 2024, Zhang, Han et al. 2025). RALF SSPs are essential for cell wall integrity, receptor activation, and stress adaptation. They interact with FERONIA receptor kinases to modulate calcium signaling, pH homeostasis, and endocytosis (Yu, Chakravorty et al. 2018, Zhu, Martinez Pacheco et al. 2020). They regulate pollen tube reception (AtRALF4;(Mecchia, Santos-Fernandez et al. 2017, Feng, Liu et al. 2019)), root elongation (AtRALF1; (Yu, Chakravorty et al. 2018, Zhu, Martinez Pacheco et al. 2020), and cell wall modification via pectin interactions (AtRALF4 and AtRALF22; (Moussu, Lee et al. 2023, Schoenaers, Lee et al. 2024)). Stress responses are mediated through FER receptor internalization (AtRALF1;(Liu, Liu et al. 2024), linking RALFs to environmental adaptation.

In the current study, *SbRALF/R 2,5,19,20,22* and *SbRALF23* were expressed during stem development at low levels in regions of cell proliferation/elongation and at higher levels in tissues that were undergoing differentiation. Elevated expression of these *SbRALF* genes in the nodal plexus and pulvinus of phytomers 6/7 was noteworthy. In addition, *SbRALFR2,19* and *22* were expressed in internode vascular parenchyma cells in fully elongated internode tissue of phytomer 7. The function of the SbRALF/R genes listed above in stem tissue differentiation is currently unknown.

### Conclusion, next steps, and study limitations

This research contributes to our understanding of bioenergy sorghum by identifying sorghum genes that encode SSPs known to regulate diverse functions in plants. Since biomass production is largely dependent on stem growth, optimizing SSPs that influence stem growth could improve biomass yield. Additionally, modifying the SSP pathways through overexpression or CRISPR knockouts could fine-tune internode elongation, biomass yield and composition. The specific functions of sorghum SSPs will require the generation of overexpression or knockout lines to validate or discover SSP roles in stem growth and development. Given their small size and bioactive properties, synthetic SSPs could also be applied exogenously to test their effects on stem growth. Furthermore, since many SSPs exhibit cell-type-specific expression, integrating developmental time-course data from laser capture microdissection (LCM) could refine our understanding of their roles. Establishing gene regulatory networks that incorporate these SSPs, growth regulators, and cell wall biosynthesis genes could also improve our understanding of stem growth regulation. This study expands knowledge of SSPs in sorghum, a drought tolerant C4 grass of great importance in agriculture. By integrating phylogenetics, transcriptomics, and expression profiling during stem development, this research provides a valuable dataset for understanding the roles of SSPs in stem development. Further characterization of SSPs involved in growth regulation could provide strategies to increase biomass yields through targeted genetic or agronomic interventions.

## Supporting information

Tree Files

## Data availability statement

The data supporting this study are available from multiple repositories. RNA-seq data from developing stem internodes (internodes 1–4) of R.07020, generated by Kebrom et al., 2017, are available at the NCBI under accession number GSE98817. Organ specificity data used for Tau analysis in the BTx623 Atlas dataset are accessible via the JGI Genome Portal under project IDs 1051037, 1051406, and 1053825. Laser capture microdissection (LCM) RNA-seq data from Fu et al., 2024, are available in the NCBI Sequence Read Archive under accession numbers SRA558272, SRA558514, and SRA558539. Additionally, data from the Sorghum bicolor TX08001 Compendium Set2 Gene Expression Profiling project can be accessed through the JGI Genome Portal (https://genome.jgi.doe.gov/portal) under JGI Project ID 1454838. To analyze the phylogenetic trees directly in iTOL via link https://itol.embl.de/shared/1gT26PPKrO3dq or can be accessed with a phylogenetic viewer of choice by downloading the .tree files within the supplementary files.

## Author contributions

EK and JM conceived the original research plan. EK performed the experiments. EK, JM, BAM analyzed the data. EK and JM wrote the manuscript.

## Acknowledgements

The authors thank Kerrie Barry and the Joint Genome Institute for contributing to RNA-seq analysis. The work (proposal: 10.46936/10.25585/60001026 and 10.46936/10.25585/60000987) conducted by the U.S. Department of Energy Joint Genome Institute (https://ror.org/04xm1d337), a DOE Office of Science User Facility, is supported by the Office of Science of the US Department of Energy operated under contract no. DE-AC02-05CH11231. The authors acknowledge Brock D. Weers for developing methods for building rhizotrons for bioenergy sorghum.

## Supplementary Figure Legends

**S-Figure 1. Maximum likelihood phylogenetic analysis of CLE SSP sequences in multiple plant species.** The phylogeny includes sequences from Arabidopsis *thaliana* TAIR10 (At), *Oryza sativa* v7.0 (Os), *Brachypodium distachyon* v3.2 (Bd), *Triticum aestivum* v2.2 (Ta), *Zea mays* RefGen_V4 (Zm), and *Sorghum bicolor* v3.1.1 (Sb). Sorghum genes encoding SSPs are highlighted in blue. Genes encoding Sorghum SSPs are named based on their phylogenetic order in **S-Figure 20**. Genes encoding SSPs in other species are named based on published nomenclature. The tree was midpoint rooted. The scale bar represents the average number of amino acid substitutions per site. Trees were annotated in Dendroscope3 and visualized using iTOL v7.0.

**S-Figure 2. Maximum likelihood phylogenetic analysis of GASA SSP sequences in multiple plant species.** The phylogeny includes sequences from Arabidopsis *thaliana* TAIR10 (At), *Oryza sativa* v7.0 (Os), *Brachypodium distachyon* v3.2 (Bd), *Triticum aestivum* v2.2 (Ta), *Zea mays* RefGen_V4 (Zm), and *Sorghum bicolor* v3.1.1 (Sb). Sorghum genes encoding SSPs are highlighted in blue. Genes encoding Sorghum SSPs are named based on their phylogenetic order in **S-Figure 21**. Genes encoding SSPs in other species are named based on published nomenclature. The tree was midpoint rooted. The scale bar represents the average number of amino acid substitutions per site. Trees were annotated in Dendroscope3 and visualized using iTOL v7.0.

**S-Figure 3. Maximum likelihood phylogenetic analysis of RALF SSP sequences in multiple plant species.** The phylogeny includes sequences from Arabidopsis *thaliana* TAIR10 (At), *Oryza sativa* v7.0 (Os), *Brachypodium distachyon* v3.2 (Bd), *Triticum aestivum* v2.2 (Ta), *Zea mays* RefGen_V4 (Zm), and *Sorghum bicolor* v3.1.1 (Sb). Sorghum genes encoding SSPs are highlighted in blue. Genes encoding Sorghum SSPs are named based on their phylogenetic order in **S-Figure 22**. Genes encoding SSPs in other species are named based on published nomenclature. The tree was rooted to AtRALF18. The scale bar represents the average number of amino acid substitutions per site. Trees were annotated in Dendroscope3 and visualized using iTOL v7.0.

**S-Figure 4. Maximum likelihood phylogenetic analysis of RGF SSP sequences in multiple plant species.** The phylogeny includes sequences from Arabidopsis *thaliana* TAIR10 (At), *Oryza sativa* v7.0 (Os), *Brachypodium distachyon* v3.2 (Bd), *Triticum aestivum* v2.2 (Ta), *Zea mays* RefGen_V4 (Zm), and *Sorghum bicolor* v3.1.1 (Sb). Sorghum genes encoding SSPs are highlighted in blue. Genes encoding Sorghum SSPs are named based on their phylogenetic order in **S-Figure 23**. Genes encoding SSPS in other species are named based on published nomenclature. The tree was midpoint rooted. The scale bar represents the average number of amino acid substitutions per site. Trees were annotated in Dendroscope3 and visualized using iTOL v7.0.

**S-Figure 5. Maximum likelihood phylogenetic analysis of CAPE SSP sequences in multiple plant species.** The phylogeny includes sequences from Arabidopsis *thaliana* TAIR10 (At), *Oryza sativa* v7.0 (Os), *Brachypodium distachyon* v3.2 (Bd), *Triticum aestivum* v2.2 (Ta), *Zea mays* RefGen_V4 (Zm), and *Sorghum bicolor* v3.1.1 (Sb). Sorghum genes encoding SSPs are highlighted in blue. Genes encoding Sorghum SSPs are named based on their phylogenetic order in **S-Figure 24**. Genes encoding SSPs in other species are named based on published nomenclature. The tree was midpoint rooted. The scale bar represents the average number of amino acid substitutions per site. Trees were annotated in Dendroscope3 and visualized using iTOL v7.0.

**S-Figure 6. Maximum likelihood phylogenetic analysis of CEP SSP sequences in multiple plant species.** The phylogeny includes sequences from Arabidopsis *thaliana* TAIR10 (At), *Oryza sativa* v7.0 (Os), *Brachypodium distachyon* v3.2 (Bd), *Triticum aestivum* v2.2 (Ta), *Zea mays* RefGen_V4 (Zm), and *Sorghum bicolor* v3.1.1 (Sb). Sorghum SSPs are highlighted in blue. Genes encoding Sorghum SSPs are named based on their phylogenetic order in **S-Figure 25**. Genes encoding SSPs in other species are named based on published nomenclature. The tree was midpoint rooted. The scale bar represents the average number of amino acid substitutions per site. Trees were annotated in Dendroscope3 and visualized using iTOL v7.0.

**S-Figure 7. Maximum likelihood phylogenetic analysis of CIF SSP sequences in multiple plant species.** The phylogeny includes sequences from Arabidopsis *thaliana* TAIR10 (At), *Oryza sativa* v7.0 (Os), *Brachypodium distachyon* v3.2 (Bd), *Triticum aestivum* v2.2 (Ta), *Zea mays* RefGen_V4 (Zm), and *Sorghum bicolor* v3.1.1 (Sb). Sorghum SSPs are highlighted in blue. Genes encoding Sorghum SSPs are named based on their phylogenetic order in **S-Figure 26**. Genes encoding SSPs in other species are named based on published nomenclature. The tree was midpoint rooted. The scale bar represents the average number of amino acid substitutions per site. Trees were annotated in Dendroscope3 and visualized using iTOL v7.0.

**S-Figure 8. Maximum likelihood phylogenetic analysis of DVL SSP sequences in multiple plant species.** The phylogeny includes sequences from Arabidopsis *thaliana* TAIR10 (At), *Oryza sativa* v7.0 (Os), *Brachypodium distachyon* v3.2 (Bd), *Triticum aestivum* v2.2 (Ta), *Zea mays* RefGen_V4 (Zm), and *Sorghum bicolor* v3.1.1 (Sb). Sorghum genes encoding SSPs are highlighted in blue. Genes encoding Sorghum SSPs are named based on their phylogenetic order in **S-Figure 27**. Genes encoding SSPs in other species are named based on published nomenclature. The tree was midpoint rooted. The scale bar represents the average number of amino acid substitutions per site. Trees were annotated in Dendroscope3 and visualized using iTOL v7.0.

**S-Figure 9. Maximum likelihood phylogenetic analysis of ECL SSP sequences in multiple plant species.** The phylogeny includes sequences from Arabidopsis *thaliana* TAIR10 (At), *Oryza sativa* v7.0 (Os), *Brachypodium distachyon* v3.2 (Bd), *Triticum aestivum* v2.2 (Ta), *Zea mays* RefGen_V4 (Zm), and *Sorghum bicolor* v3.1.1 (Sb). Sorghum genes encoding SSPs are highlighted in blue. Genes encoding SSPs in other species are named based on published nomenclature. The tree was midpoint rooted. The scale bar represents the average number of amino acid substitutions per site. Trees were annotated in Dendroscope3 and visualized using iTOL v7.0.

**S-Figure 10. Maximum likelihood phylogenetic analysis of EPF SSP sequences in multiple plant species.** The phylogeny includes sequences from Arabidopsis *thaliana* TAIR10 (At), *Oryza sativa* v7.0 (Os), *Brachypodium distachyon* v3.2 (Bd), *Triticum aestivum* v2.2 (Ta), *Zea mays* RefGen_V4 (Zm), and *Sorghum bicolor* v3.1.1 (Sb). Sorghum genes encoding SSPs are highlighted in blue. Genes encoding Sorghum SSPs are named based on their phylogenetic order in **S-Figure 28**. Genes encoding SSPs in other species are named based on published nomenclature. The tree was midpoint rooted. The scale bar represents the average number of amino acid substitutions per site. Trees were annotated in Dendroscope3 and visualized using iTOL v7.0.

**S-Figure 11. Maximum likelihood phylogenetic analysis of ES SSP sequences in multiple plant species.** The phylogeny includes sequences from Arabidopsis *thaliana* TAIR10 (At), *Oryza sativa* v7.0 (Os), *Brachypodium distachyon* v3.2 (Bd), *Triticum aestivum* v2.2 (Ta), *Zea mays* RefGen_V4 (Zm), and *Sorghum bicolor* v3.1.1 (Sb). Sorghum genes encoding SSPs are highlighted in blue. Genes encoding Sorghum SSPs are named based on their phylogenetic order in **S-Figure 29**. Genes encoding SSPs in other species are named based on published nomenclature. The tree was midpoint rooted. The scale bar represents the average number of amino acid substitutions per site. Trees were annotated in Dendroscope3 and visualized using iTOL v7.0.

**S-Figure 12. Maximum likelihood phylogenetic analysis of IDA SSP sequences in multiple plant species.** The phylogeny includes sequences from Arabidopsis *thaliana* TAIR10 (At), *Oryza sativa* v7.0 (Os), *Brachypodium distachyon* v3.2 (Bd), *Triticum aestivum* v2.2 (Ta), *Zea mays* RefGen_V4 (Zm), and *Sorghum bicolor* v3.1.1 (Sb). Sorghum genes encoding SSPs are highlighted in blue. Genes encoding Sorghum SSPs are named based on their phylogenetic order in **S-Figure 30**. Genes encoding SSPs in other species are named based on published nomenclature. The tree was midpoint rooted. The scale bar represents the average number of amino acid substitutions per site. Trees were annotated in Dendroscope3 and visualized using iTOL v7.0.

**S-Figure 13. Maximum likelihood phylogenetic analysis of MEG SSP sequences in multiple plant species.** The phylogeny includes sequences from Arabidopsis *thaliana* TAIR10 (At), *Oryza sativa* v7.0 (Os), *Brachypodium distachyon* v3.2 (Bd), *Triticum aestivum* v2.2 (Ta), *Zea mays* RefGen_V4 (Zm), and *Sorghum bicolor* v3.1.1 (Sb). Sorghum genes encoding SSPs are highlighted in blue. Genes encoding Sorghum SSPs are named based on their phylogenetic order in **S-Figure 31**. Genes encoding SSPs in other species are named based on published nomenclature. The tree was midpoint rooted. The scale bar represents the average number of amino acid substitutions per site. Trees were annotated in Dendroscope3 and visualized using iTOL v7.0.

**S-Figure 14. Maximum likelihood phylogenetic analysis of PIP SSP sequences in multiple plant species.** The phylogeny includes sequences from Arabidopsis *thaliana* TAIR10 (At), *Oryza sativa* v7.0 (Os), *Brachypodium distachyon* v3.2 (Bd), *Triticum aestivum* v2.2 (Ta), *Zea mays* RefGen_V4 (Zm), and *Sorghum bicolor* v3.1.1 (Sb). Sorghum genes encoding SSPs are highlighted in blue. Genes encoding Sorghum SSPs are named based on their phylogenetic order in **S-Figure 32**. Genes encoding SSPs in other species are named based on published nomenclature. The tree was midpoint rooted. The scale bar represents the average number of amino acid substitutions per site. Trees were annotated in Dendroscope3 and visualized using iTOL v7.0.

**S-Figure 15. Maximum likelihood phylogenetic analysis of PNP SSP sequences in multiple plant species.** The phylogeny includes sequences from Arabidopsis *thaliana* TAIR10 (At), *Oryza sativa* v7.0 (Os), *Brachypodium distachyon* v3.2 (Bd), *Triticum aestivum* v2.2 (Ta), *Zea mays* RefGen_V4 (Zm), and *Sorghum bicolor* v3.1.1 (Sb). Sorghum genes encoding SSPs are highlighted in blue. Genes encoding Sorghum SSPs are named based on their phylogenetic order in **S-Figure 33**. Genes encoding SSPs in other species are named based on published nomenclature. The tree was midpoint rooted. The scale bar represents the average number of amino acid substitutions per site. Trees were annotated in Dendroscope3 and visualized using iTOL v7.0.

**S-Figure 16. Maximum likelihood phylogenetic analysis of POE SSP sequences in multiple plant species.** The phylogeny includes sequences from Arabidopsis *thaliana* TAIR10 (At), *Oryza sativa* v7.0 (Os), *Brachypodium distachyon* v3.2 (Bd), *Triticum aestivum* v2.2 (Ta), *Zea mays* RefGen_V4 (Zm), and *Sorghum bicolor* v3.1.1 (Sb). Sorghum genes encoding SSPs are highlighted in blue. Genes encoding Sorghum SSPs are named based on their phylogenetic order in **S-Figure 34**. Genes encoding SSPs in other species are named based on published nomenclature. The tree was midpoint rooted. The scale bar represents the average number of amino acid substitutions per site. Trees were annotated in Dendroscope3 and visualized using iTOL v7.0.

**S-Figure 17. Maximum likelihood phylogenetic analysis of PSK SSP sequences in multiple plant species.** The phylogeny includes sequences from Arabidopsis *thaliana* TAIR10 (At), *Oryza sativa* v7.0 (Os), *Brachypodium distachyon* v3.2 (Bd), *Triticum aestivum* v2.2 (Ta), *Zea mays* RefGen_V4 (Zm), and *Sorghum bicolor* v3.1.1 (Sb). Sorghum genes encoding SSPs are highlighted in blue. Genes encoding Sorghum SSPs are named based on their phylogenetic order in **S-Figure 35**. Genes encoding SSPs in other species are named based on published nomenclature. The tree was midpoint rooted. The scale bar represents the average number of amino acid substitutions per site. Trees were annotated in Dendroscope3 and visualized using iTOL v7.0.

**S-Figure 18. Maximum likelihood phylogenetic analysis of PSY SSP sequences in multiple plant species.** The phylogeny includes sequences from Arabidopsis *thaliana* TAIR10 (At), *Oryza sativa* v7.0 (Os), *Brachypodium distachyon* v3.2 (Bd), *Triticum aestivum* v2.2 (Ta), *Zea mays* RefGen_V4 (Zm), and *Sorghum bicolor* v3.1.1 (Sb). Sorghum genes encoding SSPs are highlighted in blue. Genes encoding Sorghum SSPs are named based on their phylogenetic order in **S-Figure 36**. Genes encoding SSPs in other species are named based on published nomenclature. The tree was midpoint rooted. The scale bar represents the average number of amino acid substitutions per site. Trees were annotated in Dendroscope3 and visualized using iTOL v7.0.

**S-Figure 19. Maximum likelihood phylogenetic analysis of TPD SSP sequences in multiple plant species.** The phylogeny includes sequences from Arabidopsis *thaliana* TAIR10 (At), *Oryza sativa* v7.0 (Os), *Brachypodium distachyon* v3.2 (Bd), *Triticum aestivum* v2.2 (Ta), *Zea mays* RefGen_V4 (Zm), and *Sorghum bicolor* v3.1.1 (Sb). Sorghum genes encoding SSPs are highlighted in blue. Genes encoding Sorghum SSPs are named based on their phylogenetic order in **S-Figure 37**. Genes encoding SSPs in other species are named based on published nomenclature. The tree was midpoint rooted. The scale bar represents the average number of amino acid substitutions per site. Trees were annotated in Dendroscope3 and visualized using iTOL v7.0.

**S-Figure 20. Maximum likelihood phylogenetic analysis of CLE SSP sequences for *Sorghum bicolor*.** The scale bar represents the average number of amino acid substitutions per site. Tree was annotated and visualized in iTOL v7.0.

**S-Figure 21. Maximum likelihood phylogenetic analysis of GASS SSP sequences for *Sorghum bicolor*.** The scale bar represents the average number of amino acid substitutions per site. Tree was annotated and visualized in iTOL v7.0.

**S-Figure 22. Maximum likelihood phylogenetic analysis of RALF SSP sequences for *Sorghum bicolor*.** The scale bar represents the average number of amino acid substitutions per site. Tree was annotated and visualized in iTOL v7.0.

**S-Figure 23. Maximum likelihood phylogenetic analysis of RGF SSP sequences for *Sorghum bicolor*.** The scale bar represents the average number of amino acid substitutions per site. Tree was annotated and visualized in iTOL v7.0.

**S-Figure 24. Maximum likelihood phylogenetic analysis of CAPE SSP sequences for *Sorghum bicolor*.** The scale bar represents the average number of amino acid substitutions per site. Tree was annotated and visualized in iTOL v7.0.

**S-Figure 25. Maximum likelihood phylogenetic analysis of CEP SSP sequences for *Sorghum bicolor*.** The scale bar represents the average number of amino acid substitutions per site. Tree was annotated and visualized in iTOL v7.0.

**S-Figure 26. Maximum likelihood phylogenetic analysis of CIF SSP sequences for *Sorghum bicolor*.** The scale bar represents the average number of amino acid substitutions per site. Tree was annotated and visualized in iTOL v7.0.

**S-Figure 27. Maximum likelihood phylogenetic analysis of DVL SSP sequences for *Sorghum bicolor*.** The scale bar represents the average number of amino acid substitutions per site. Tree was annotated and visualized in iTOL v7.0.

**S-Figure 28. Maximum likelihood phylogenetic analysis of EPF SSP sequences for *Sorghum bicolor*.** The scale bar represents the average number of amino acid substitutions per site. Tree was annotated and visualized in iTOL v7.0.

**S-Figure 29. Maximum likelihood phylogenetic analysis of ES SSP sequences for *Sorghum bicolor*.** The scale bar represents the average number of amino acid substitutions per site. Tree was annotated and visualized in iTOL v7.0.

**S-Figure 30. Maximum likelihood phylogenetic analysis of IDA SSP sequences for *Sorghum bicolor*.** The scale bar represents the average number of amino acid substitutions per site. Tree was annotated and visualized in iTOL v7.0.

**S-Figure 31. Maximum likelihood phylogenetic analysis of MEG SSP sequences for *Sorghum bicolor*.** The scale bar represents the average number of amino acid substitutions per site. Tree was annotated and visualized in iTOL v7.0.

**S-Figure 32. Maximum likelihood phylogenetic analysis of PIP SSP sequences for *Sorghum bicolor*.** The scale bar represents the average number of amino acid substitutions per site. Tree was annotated and visualized in iTOL v7.0.

**S-Figure 33. Maximum likelihood phylogenetic analysis of PNP SSP sequences for *Sorghum bicolor*.** The scale bar represents the average number of amino acid substitutions per site. Tree was annotated and visualized in iTOL v7.0.

**S-Figure 34. Maximum likelihood phylogenetic analysis of POE SSP sequences for *Sorghum bicolor*.** The scale bar represents the average number of amino acid substitutions per site. Tree was annotated and visualized in iTOL v7.0.

**S-Figure 35. Maximum likelihood phylogenetic analysis of PSK SSP sequences for *Sorghum bicolor*.** The scale bar represents the average number of amino acid substitutions per site. Tree was annotated and visualized in iTOL v7.0.

**S-Figure 36. Maximum likelihood phylogenetic analysis of PSY SSP sequences for *Sorghum bicolor*.** The scale bar represents the average number of amino acid substitutions per site. Tree was annotated and visualized in iTOL v7.0.

**S-Figure 37. Maximum likelihood phylogenetic analysis of TPD SSP sequences for *Sorghum bicolor*.** The scale bar represents the average number of amino acid substitutions per site. Tree was annotated and visualized in iTOL v7.0.

**S-Figure 38. Expression of 158 genes encoding small signaling SSPs in 19 small signaling peptide families in leaf, stem, root, and panicle tissues.** SSPs were sorted based on phylogenetic order and 158 of the 219 were selected using a >5 TPM threshold across the averaged expression values within each organ. SbECL2 has a max TPM of 3 but was included to not remove the family. Expression values are reported in TPM, with blue indicating high expression and green indicating low expression.

## Supplementary Table Legends

**S-Table 1. List of SSP encoding genes in sorghum with their gene names based on phylogenetic order in their respective sorghum phylogenetic analysis.**

**S-Table 2. CLE SSPs in Sorghum identified from homology and literature sources.** This table also includes any annotations provided by Phytozome v13 for a given peptide. Provided is the best e-value to a respective published homolog query from another species to Sorghum. Each pre-processed peptide length is provided along with the peptide structure visual provided from the MEME analysis. Previously published conserved sequences for the respective mature peptide sequences were analyzed and their presence or absence is provided in the tabular format. The results for the motif consensus are also provided as an image along with a color-coded key to show which sequence is present in the peptide of interest. The settings used for the MEME search can also be located here.

**S-Table 3. GASS SSPs in Sorghum identified from homology and literature sources.** This table also includes any annotations provided by Phytozome v13 for a given peptide. Provided is the best e-value to a respective published homolog query from another species to Sorghum. Each pre-processed peptide length is provided along with the peptide structure visual provided from the MEME analysis. Previously published conserved sequences for the respective mature peptide sequences were analyzed and their presence or absence is provided in the tabular format. The results for the motif consensus are also provided as an image along with a color-coded key to show which sequence is present in the peptide of interest. The settings used for the MEME search can also be located here.

**S-Table 4. RALF and RALFR SSPs in Sorghum identified from homology and literature sources.** This table also includes any annotations provided by Phytozome v13 for a given peptide. Provided is the best e-value to a respective published homolog query from another species to Sorghum. Each pre-processed peptide length is provided along with the peptide structure visual provided from the MEME analysis. Previously published conserved sequences for the respective mature peptide sequences were analyzed and their presence or absence is provided in the tabular format. The results for the motif consensus are also provided as an image along with a color-coded key to show which sequence is present in the peptide of interest. The settings used for the MEME search can also be located here.

**S-Table 5. RGF SSPs in Sorghum identified from homology and literature sources.** This table also includes any annotations provided by Phytozome v13 for a given peptide. Provided is the best e-value to a respective published homolog query from another species to Sorghum. Each pre-processed peptide length is provided along with the peptide structure visual provided from the MEME analysis. Previously published conserved sequences for the respective mature peptide sequences were analyzed and their presence or absence is provided in the tabular format. The results for the motif consensus are also provided as an image along with a color-coded key to show which sequence is present in the peptide of interest. The settings used for the MEME search can also be located here.

**S-Table 6. Best homolog of a *Sorghum bicolor* gene encoding peptide from the maximum likelihood analysis of CLE SSPs.** Best homolog was determined by topological proximity. The best homolog for a given Sorghum bicolor is provided for *Zea mays*, *Oryza sativa*, and *Arabidopsis thaliana* along with their bootstrap values.

**S-Table 7. Best homolog of a *Sorghum bicolor* gene encoding peptide from the maximum likelihood analysis of GASS SSPs.** Best homolog was determined by topological proximity. The best homolog for a given Sorghum bicolor is provided for *Zea mays*, *Oryza sativa*, and *Arabidopsis thaliana* along with their bootstrap values.

**S-Table 8. Best homolog of a *Sorghum bicolor* gene encoding peptide from the maximum likelihood analysis of RALF SSPs.** Best homolog was determined by topological proximity. The best homolog for a given Sorghum bicolor is provided for *Zea mays*, *Oryza sativa*, and *Arabidopsis thaliana* along with their bootstrap values.

**S-Table 9. Best hits of a Sorghum bicolor gene for homology from the maximum likelihood analysis of RGF SSPs determined by topological proximity.** The best homolog for a given Sorghum bicolor is provided for *Zea mays*, *Oryza sativa*, and *Arabidopsis thaliana* along with their bootstrap values.

**S-Table 10. All 219 SSPs in *Sorghum bicolor* expression throughout different organ samples.** All SSPs from this analysis are included along with their respective max TPM across the dataset and TAU values.

**S-Table 11. Genes encoding SSPs expressed in each internode 1-4.** This table includes genes that were not differentially expressed more than 5 FC and the rest that had max expression across the 4 tissues analyzed less than 5 from the analysis in Figure 2.

**S-Table 12. Expression of genes encoding SSPs that increase in expression during stem differentiation or throughout stem differentiation.** Genes included are expressed throughout P1-6 stem development from Figure 4a) and Figure 5a). SSPs that had TPM max <5 were also included.

**S-Table 13. Table of all 219 SSPs in the LCM analysis from Figure 2.** All SSPs are expressed with max TPM>5 across the 5 cell types are included along with their TAU values. Included are all the other SSPs that were expressed <5 TPM in the data set.

