## Supplementary material for "Analysis of Small Signaling Peptides in *Sorghum bicolor*: Integrating Phylogeny and Gene Expression to Characterize Roles in Stem Development": S-Figures

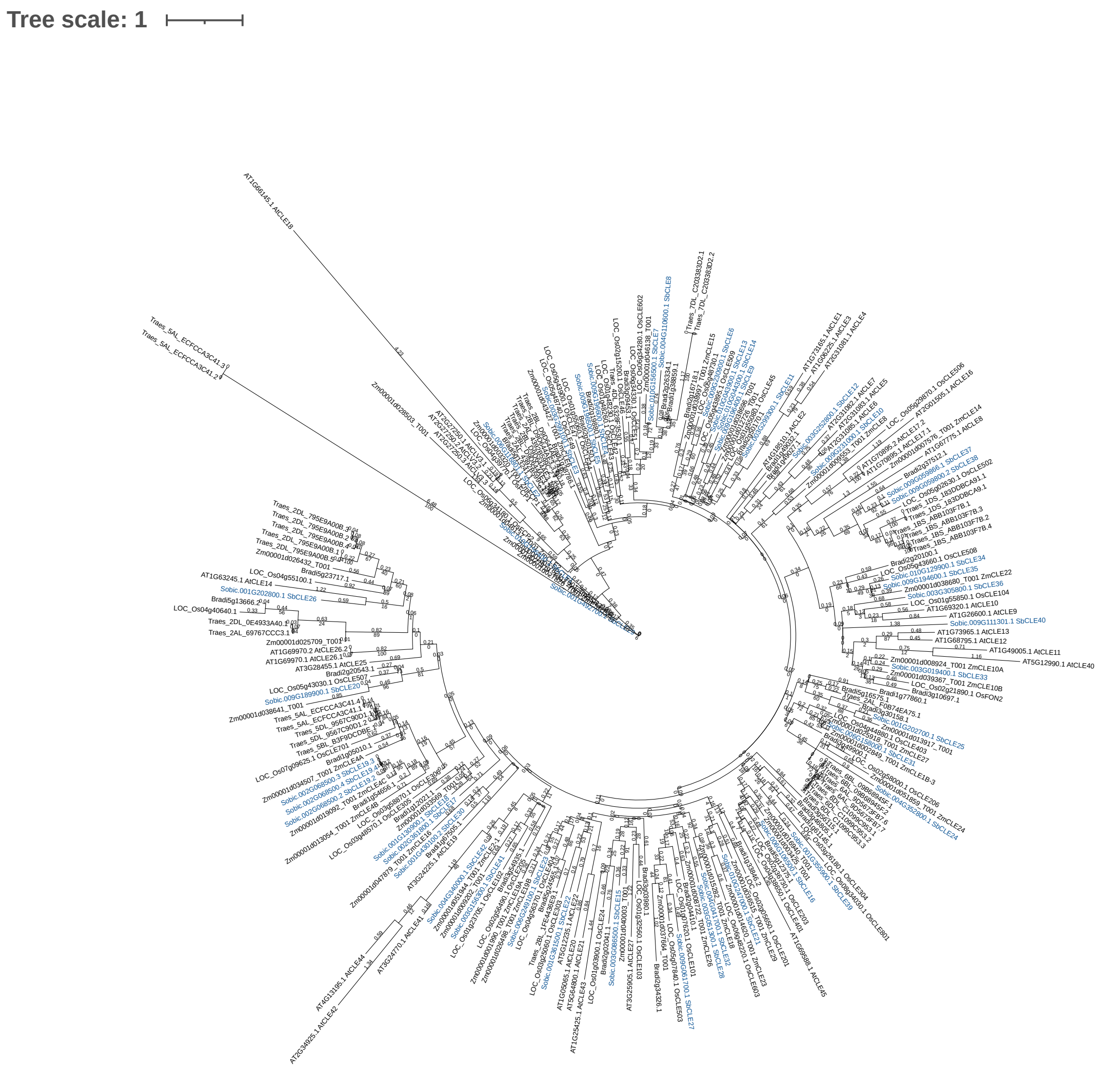


S-Figure 1. Maximum likelihood phylogenetic analysis of CLE SSP sequences in multiple plant species.
The phylogeny includes sequences from *Arabidopsis thaliana* TAIR10 (At), *Oryza sativa* v7.0 (Os), *Brachypodium distachyon* v3.2 (Bd), *Triticum aestivum* v2.2 (Ta), *Zea mays* RefGen_V4 (Zm), and *Sorghum bicolor* v3.1.1 (Sb). Sorghum genes encoding SSPs are highlighted in blue. Genes encoding Sorghum SSPs are named based on their phylogenetic order in S-Figure 20. Genes encoding SSPs in other species are named based on published nomenclature. The tree was midpoint rooted. The scale bar represents the average number of amino acid substitutions per site. Trees were annotated in Dendroscope3 and visualized using iTOL v7.0.

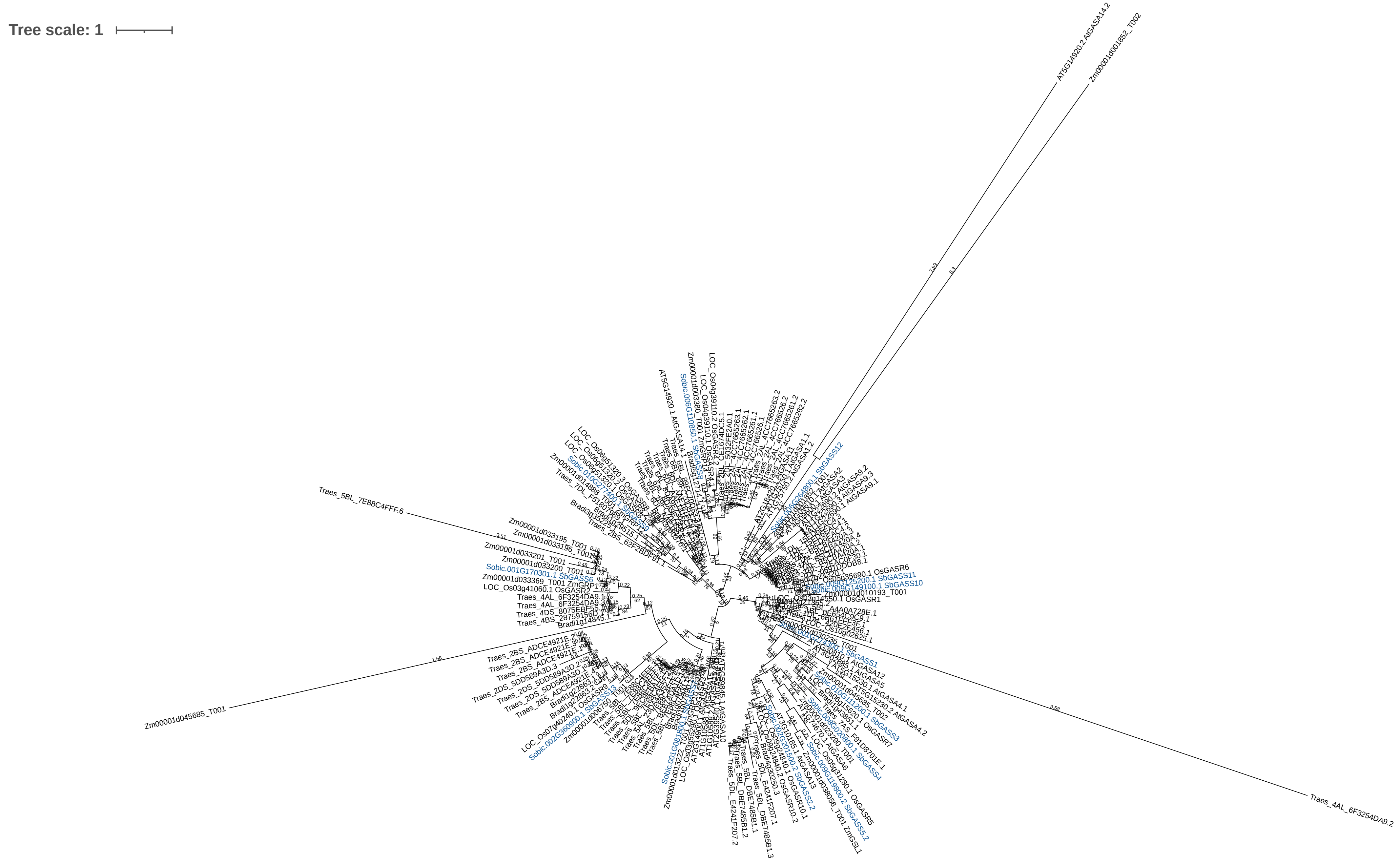


S-Figure 2. Maximum likelihood phylogenetic analysis of GASA SSP sequences in multiple plant species.
The phylogeny includes sequences from *Arabidopsis thaliana* TAIR10 (At), *Oryza sativa* v7.0 (Os), *Brachypodium distachyon* v3.2 (Bd), *Triticum aestivum* v2.2 (Ta), *Zea mays* RefGen_V4 (Zm), and *Sorghum bicolor* v3.1.1 (Sb). Sorghum genes encoding SSPs are highlighted in blue. Genes encoding Sorghum SSPs are named based on their phylogenetic order in S-Figure 21. Genes encoding SSPs in other species are named based on published nomenclature. The tree was midpoint rooted. The scale bar represents the average number of amino acid substitutions per site. Trees were annotated in Dendroscope3 and visualized using iTOL v7.0.

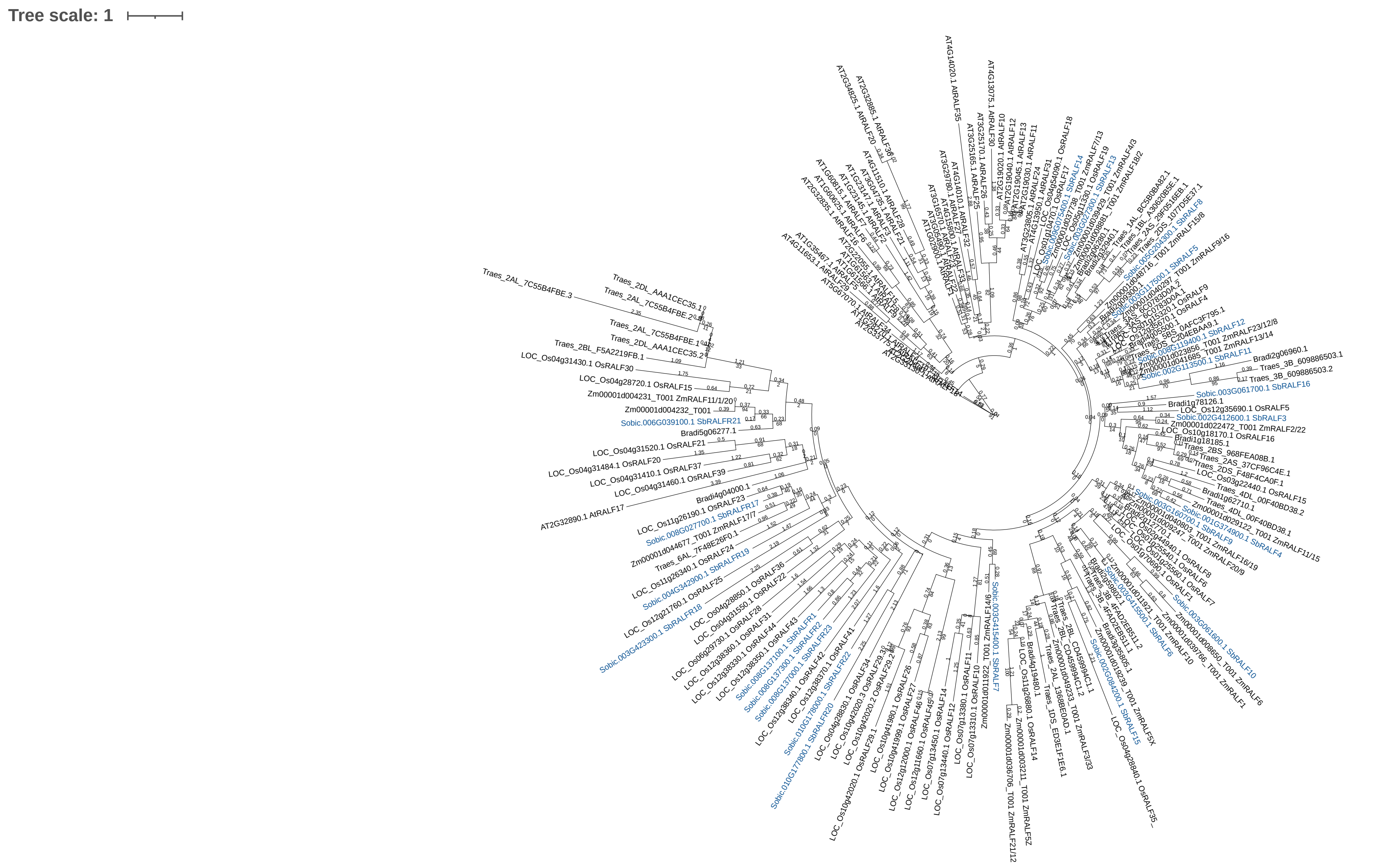


S-Figure 3. Maximum likelihood phylogenetic analysis of RALF SSP sequences in multiple plant species. The phylogeny includes sequences from *Arabidopsis thaliana* TAIR10 (At), *Oryza sativa* v7.0 (Os), *Brachypodium distachyon* v3.2 (Bd), *Triticum aestivum* v2.2 (Ta), *Zea mays* RefGen_V4 (Zm), and *Sorghum bicolor* v3.1.1 (Sb). Sorghum genes encoding SSPs are highlighted in blue. Genes encoding Sorghum SSPs are named based on their phylogenetic order in S-Figure 22. Genes encoding SSPs in other species are named based on published nomenclature.
The tree was rooted to AtRALF18. The scale bar represents the average number of amino acid substitutions per site. Trees were annotated in Dendroscope3 and visualized using iTOL v7.0.

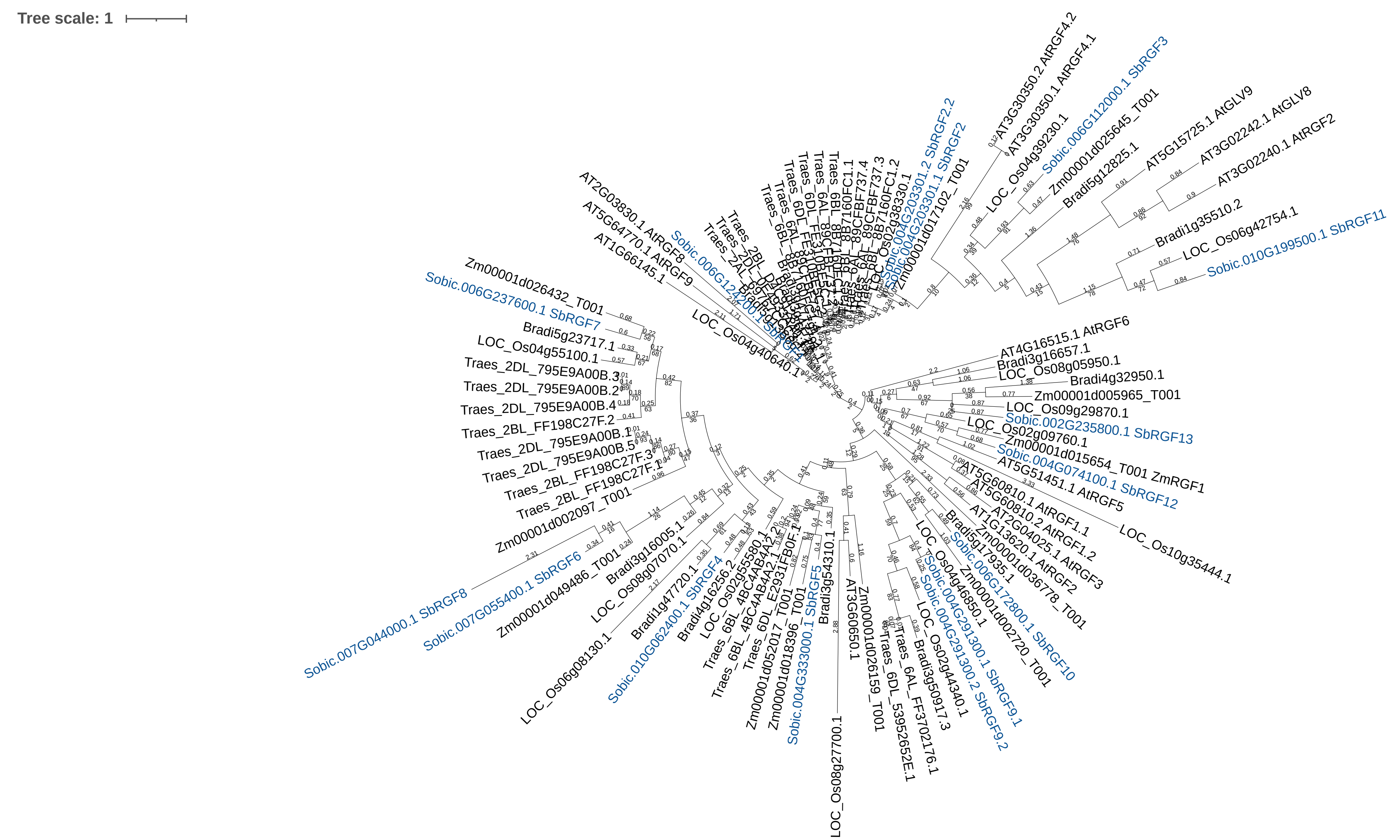


S-Figure 4. Maximum likelihood phylogenetic analysis of RGF SSP sequences in multiple plant species. The phylogeny includes sequences from *Arabidopsis thaliana* TAIR10 (At), *Oryza sativa* v7.0 (Os), *Brachypodium distachyon* v3.2 (Bd), *Triticum aestivum* v2.2 (Ta), *Zea mays* RefGen_V4 (Zm), and *Sorghum bicolor* v3.1.1 (Sb). Sorghum genes encoding SSPs are highlighted in blue. Genes encoding Sorghum SSPs are named based on their phylogenetic order in S-Figure 23. Genes encoding SSPs in other species are named based on published nomenclature. The tree was midpoint rooted. The scale bar represents the average number of amino acid substitutions per site. Trees were annotated in Dendroscope3 and visualized using iTOL v7.0.

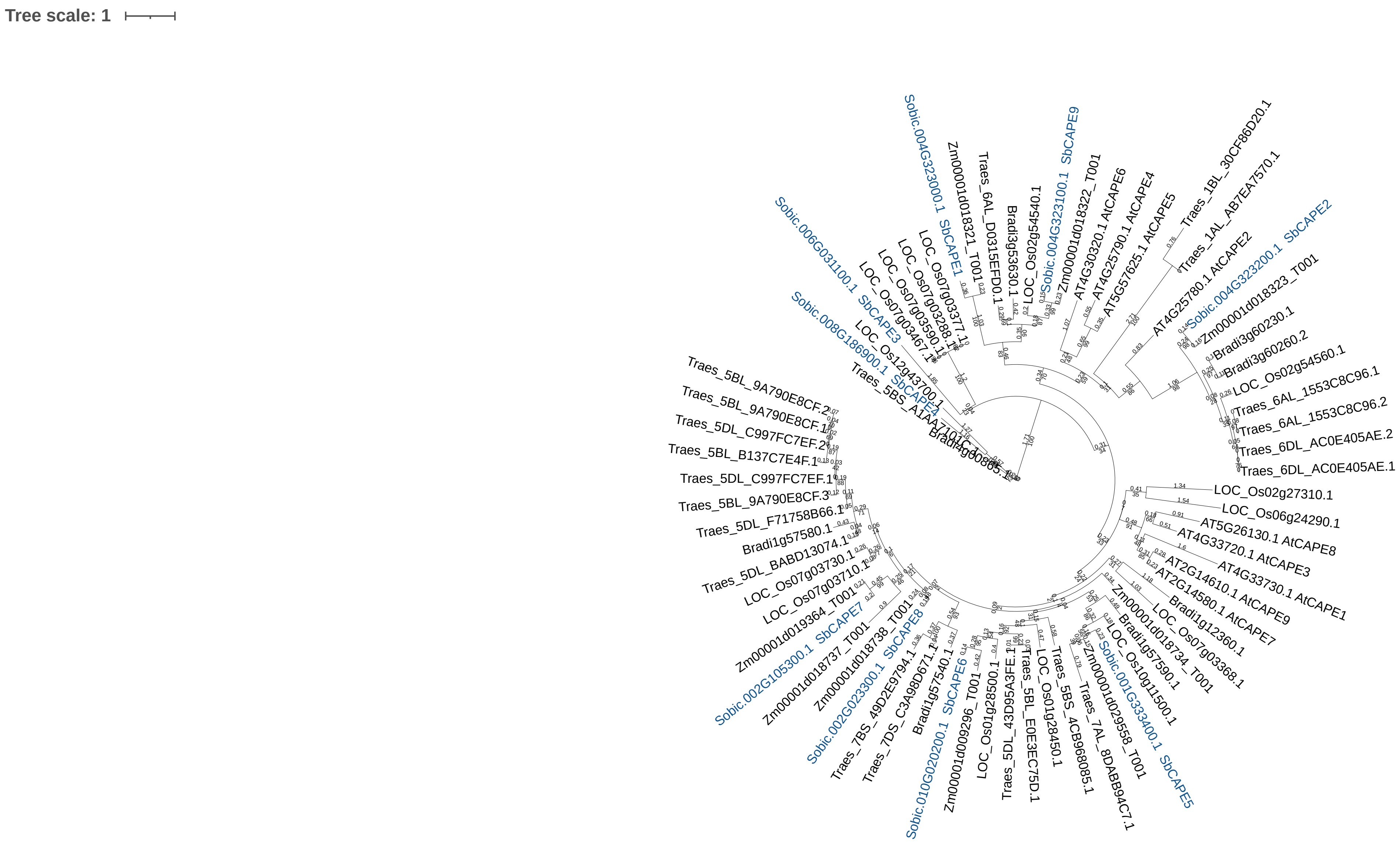


S-Figure 5. Maximum likelihood phylogenetic analysis of CAPE SSP sequences in multiple plant species. The phylogeny includes sequences from *Arabidopsis thaliana* TAIR10 (At), *Oryza sativa* v7.0 (Os), *Brachypodium distachyon* v3.2 (Bd), *Triticum aestivum* v2.2 (Ta), *Zea mays* RefGen_V4 (Zm), and *Sorghum bicolor* v3.1.1 (Sb). Sorghum genes encoding SSPs are highlighted in blue. Genes encoding Sorghum SSPs are named based on their phylogenetic order in S-Figure 24. Genes encoding SSPs in other species are named based on published nomenclature.
The tree was midpoint rooted. The scale bar represents the average number of amino acid substitutions per site.
Trees were annotated in Dendroscope3 and visualized using iTOL v7.0.

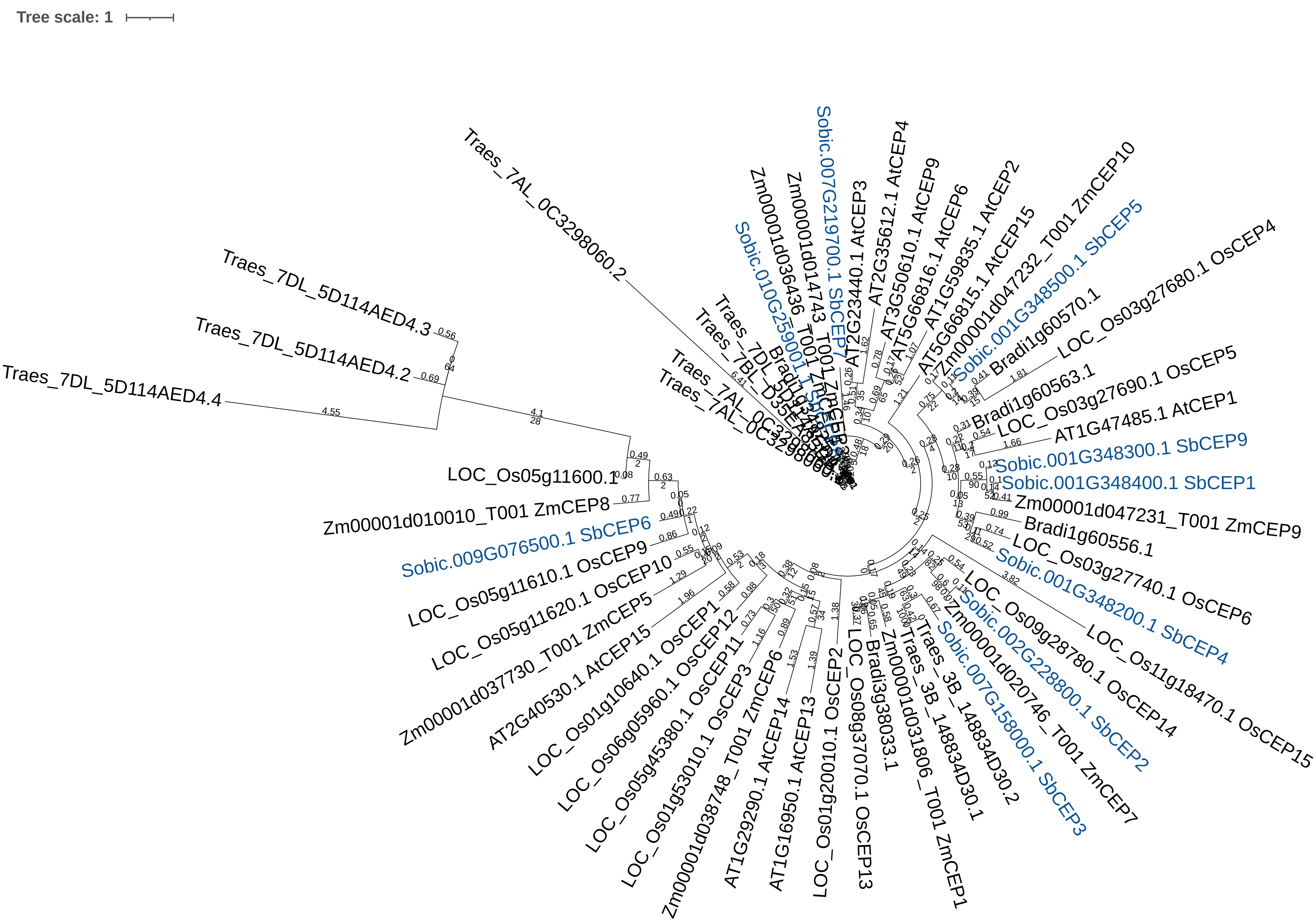


S-Figure 6. Maximum likelihood phylogenetic analysis of CEP SSP sequences in multiple plant species. The phylogeny includes sequences from *Arabidopsis thaliana* TAIR10 (At), *Oryza sativa* v7.0 (Os), *Brachypodium distachyon* v3.2 (Bd), *Triticum aestivum* v2.2 (Ta), *Zea mays* RefGen_V4 (Zm), and *Sorghum bicolor* v3.1.1 (Sb). Sorghum SSPs are highlighted in blue. Genes encoding Sorghum SSPs are named based on their phylogenetic order in S-Figure 25. Genes encoding SSPs in other species are named based on published nomenclature. The tree was midpoint rooted. The scale bar represents the average number of amino acid substitutions per site. Trees were annotated in Dendroscope3 and visualized using iTOL v7.0.

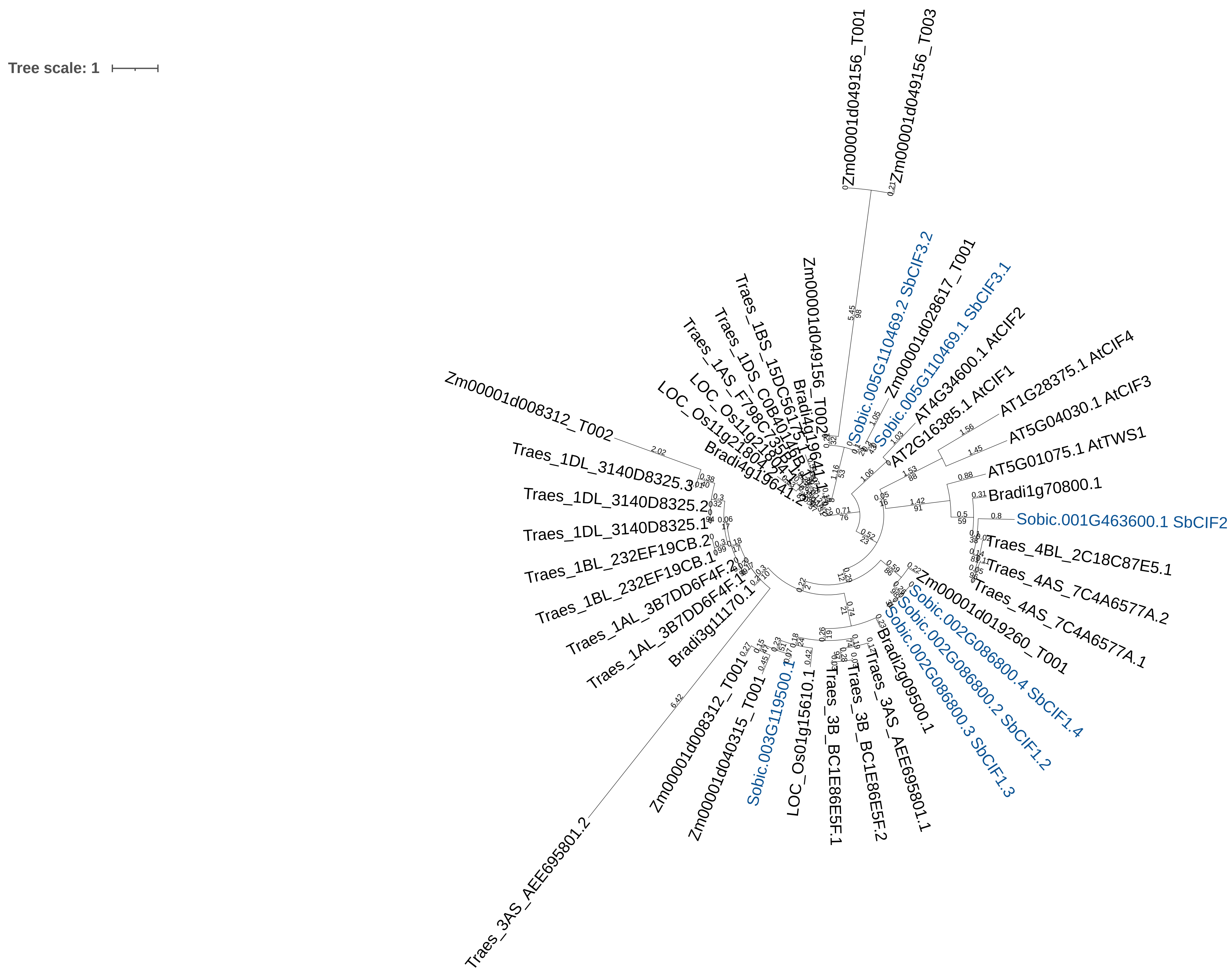


S-Figure 7. Maximum likelihood phylogenetic analysis of CIF SSP sequences in multiple plant species. The phylogeny includes sequences from *Arabidopsis thaliana* TAIR10 (At), *Oryza sativa* v7.0 (Os), *Brachypodium distachyon* v3.2 (Bd), *Triticum aestivum* v2.2 (Ta), *Zea mays* RefGen_V4 (Zm), and *Sorghum bicolor* v3.1.1 (Sb). Sorghum SSPs are highlighted in blue. Genes encoding Sorghum SSPs are named based on their phylogenetic order in S-Figure 26. Genes encoding SSPs in other species are named based on published nomenclature. The tree was midpoint rooted. The scale bar represents the average number of amino acid substitutions per site. Trees were annotated in Dendroscope3 and visualized using iTOL v7.0.

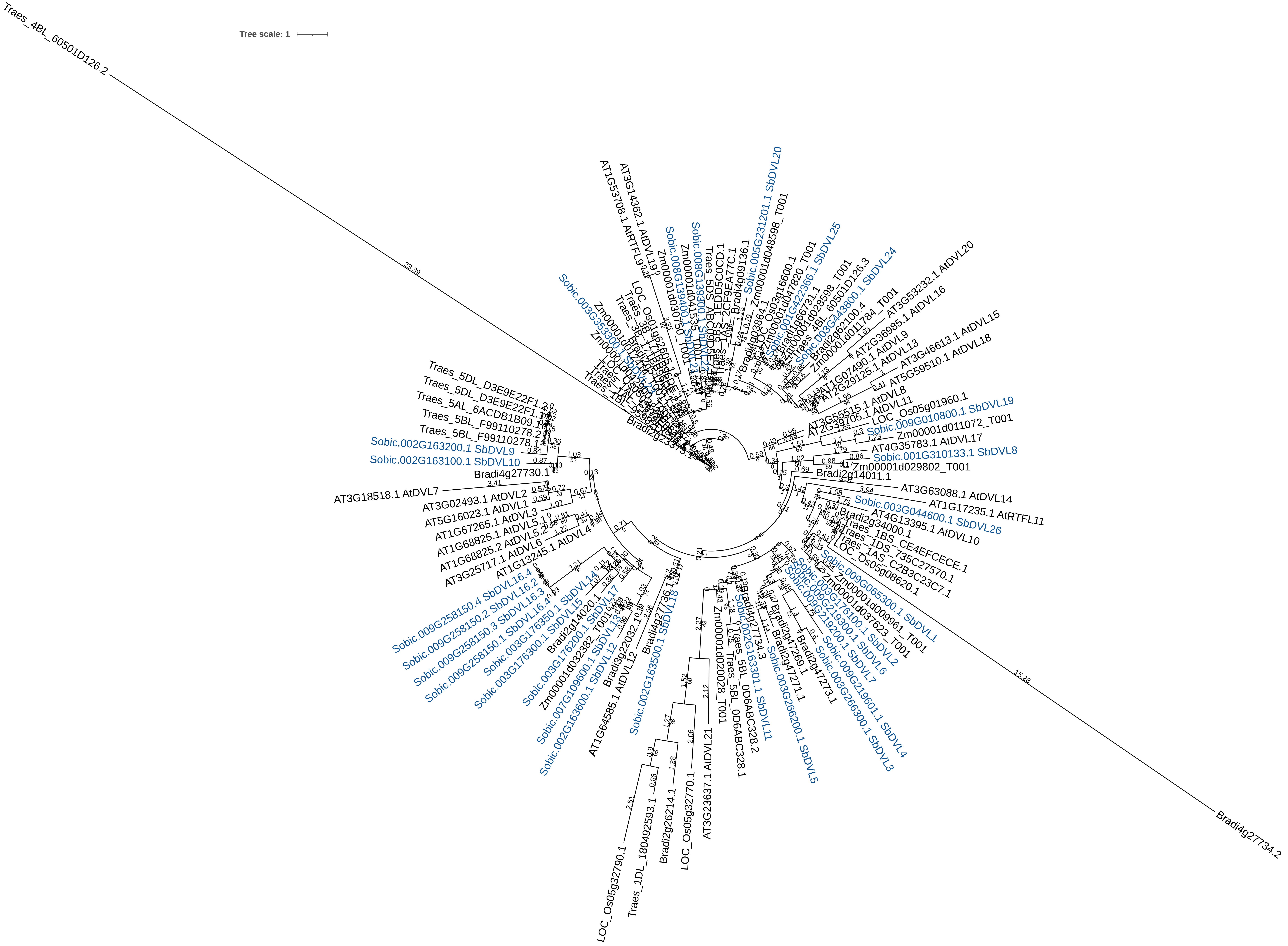


S-Figure 8. Maximum likelihood phylogenetic analysis of DVL SSP sequences in multiple plant species. The phylogeny includes sequences from *Arabidopsis thaliana* TAIR10 (At), *Oryza sativa* v7.0 (Os), *Brachypodium distachyon* v3.2 (Bd), *Triticum aestivum* v2.2 (Ta), *Zea mays* RefGen_V4 (Zm), and *Sorghum bicolor* v3.1.1 (Sb). Sorghum genes encoding SSPs are highlighted in blue. Genes encoding Sorghum SSPs are named based on their phylogenetic order in S-Figure 27. Genes encoding SSPs in other species are named based on published nomenclature. The tree was midpoint rooted. The scale bar represents the average number of amino acid substitutions per site. Trees were annotated in Dendroscope3 and visualized using iTOL v7.0.

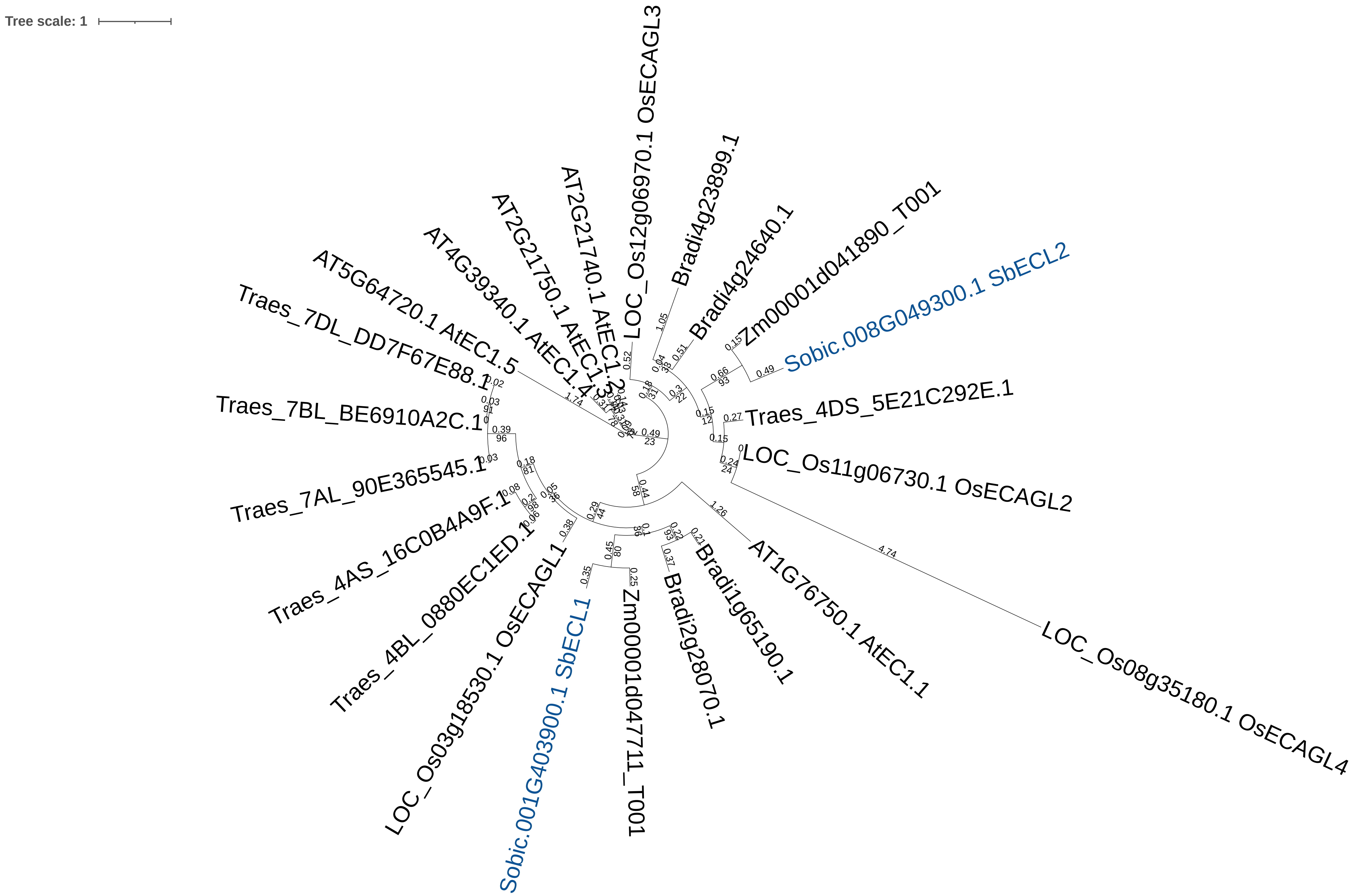


S-Figure 9. Maximum likelihood phylogenetic analysis of ECL SSP sequences in multiple plant species. The phylogeny includes sequences from *Arabidopsis thaliana* TAIR10 (At), *Oryza sativa* v7.0 (Os), *Brachypodium distachyon* v3.2 (Bd), *Triticum aestivum* v2.2 (Ta), *Zea mays* RefGen_V4 (Zm), and *Sorghum bicolor* v3.1.1 (Sb). Sorghum genes encoding SSPs are highlighted in blue. Genes encoding SSPs in other species are named based on published nomenclature. The tree was midpoint rooted. The scale bar represents the average number of amino acid substitutions per site. Trees were annotated in Dendroscope3 and visualized using iTOL v7.0.

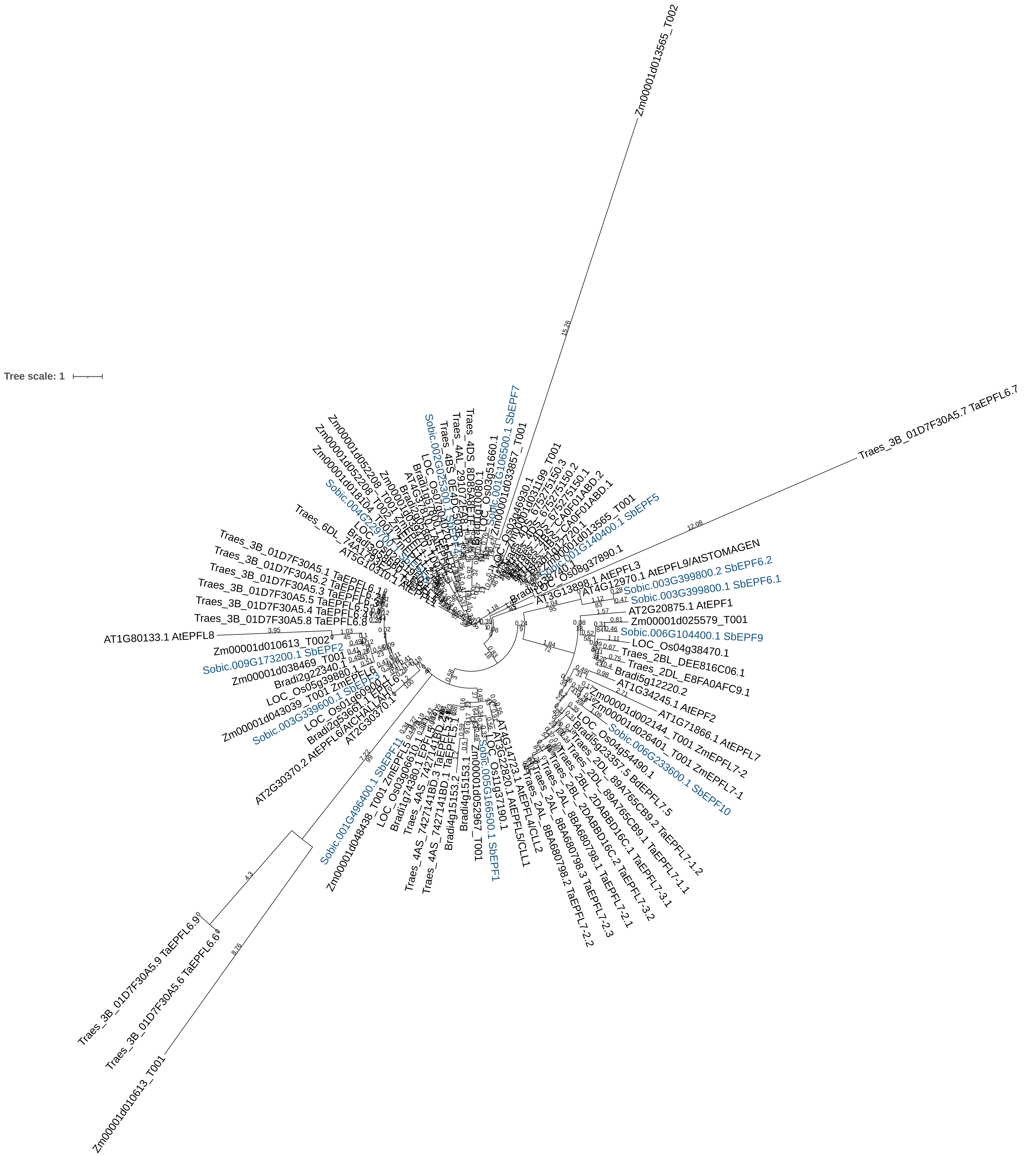


S-Figure 10. Maximum likelihood phylogenetic analysis of EPF SSP sequences in multiple plant species. The phylogeny includes sequences from *Arabidopsis thaliana* TAIR10 (At), *Oryza sativa* v7.0 (Os), *Brachypodium distachyon* v3.2 (Bd), *Triticum aestivum* v2.2 (Ta), *Zea mays* RefGen_V4 (Zm), and *Sorghum bicolor* v3.1.1 (Sb). Sorghum genes encoding SSPs are highlighted in blue. Genes encoding Sorghum SSPs are named based on their phylogenetic order in S-Figure 28. Genes encoding SSPs in other species are named based on published nomenclature. The tree was midpoint rooted. The scale bar represents the average number of amino acid substitutions per site. Trees were annotated in Dendroscope3 and visualized using iTOL v7.0.

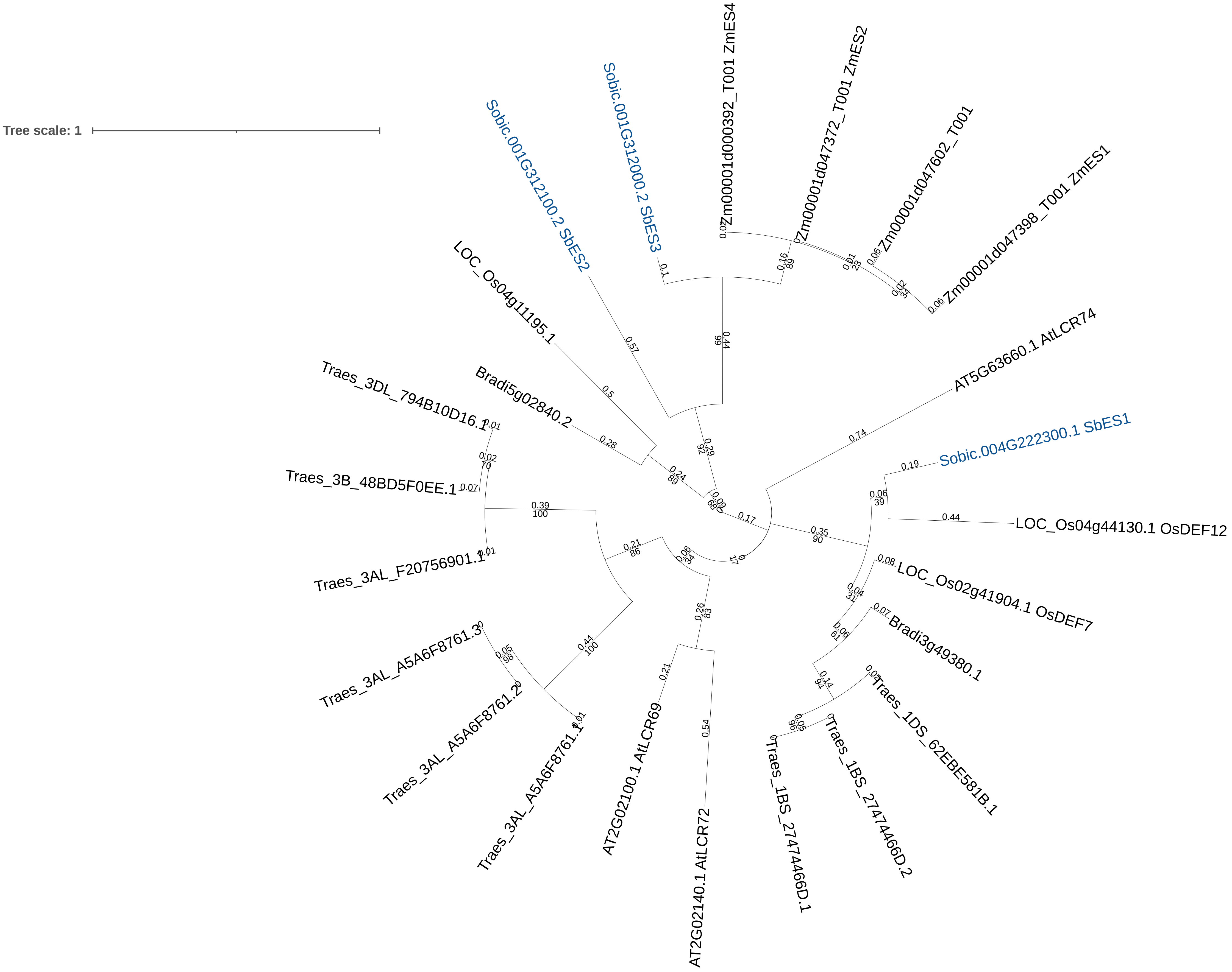


S-Figure 11. Maximum likelihood phylogenetic analysis of ES SSP sequences in multiple plant species. The phylogeny includes sequences from *Arabidopsis thaliana* TAIR10 (At), *Oryza sativa* v7.0 (Os), *Brachypodium distachyon* v3.2 (Bd), *Triticum aestivum* v2.2 (Ta), *Zea mays* RefGen_V4 (Zm), and *Sorghum bicolor* v3.1.1 (Sb). Sorghum genes encoding SSPs are highlighted in blue. Genes encoding Sorghum SSPs are named based on their phylogenetic order in S-Figure 29. Genes encoding SSPs in other species are named based on published nomenclature. The tree was midpoint rooted. The scale bar represents the average number of amino acid substitutions per site.
Trees were annotated in Dendroscope3 and visualized using iTOL v7.0.

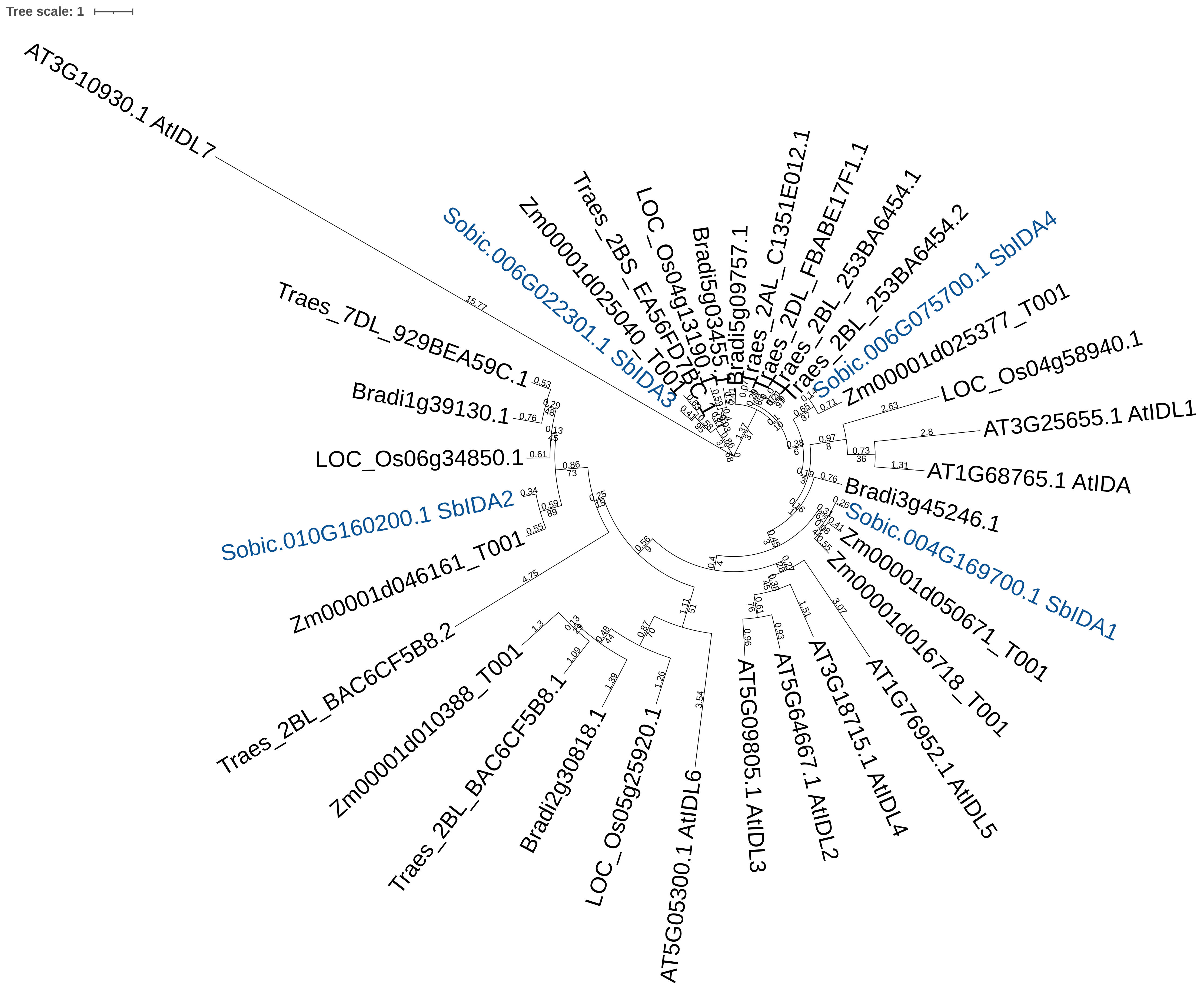


S-Figure 12. Maximum likelihood phylogenetic analysis of IDA SSP sequences in multiple plant species. The phylogeny includes sequences from *Arabidopsis thaliana* TAIR10 (At), *Oryza sativa* v7.0 (Os), *Brachypodium distachyon* v3.2 (Bd), *Triticum aestivum* v2.2 (Ta), *Zea mays* RefGen_V4 (Zm), and *Sorghum bicolor* v3.1.1 (Sb). Sorghum genes encoding SSPs are highlighted in blue. Genes encoding Sorghum SSPs are named based on their phylogenetic order in S-Figure 30. Genes encoding SSPs in other species are named based on published nomenclature. The tree was midpoint rooted. The scale bar represents the average number of amino acid substitutions per site.
Trees were annotated in Dendroscope3 and visualized using iTOL v7.0.

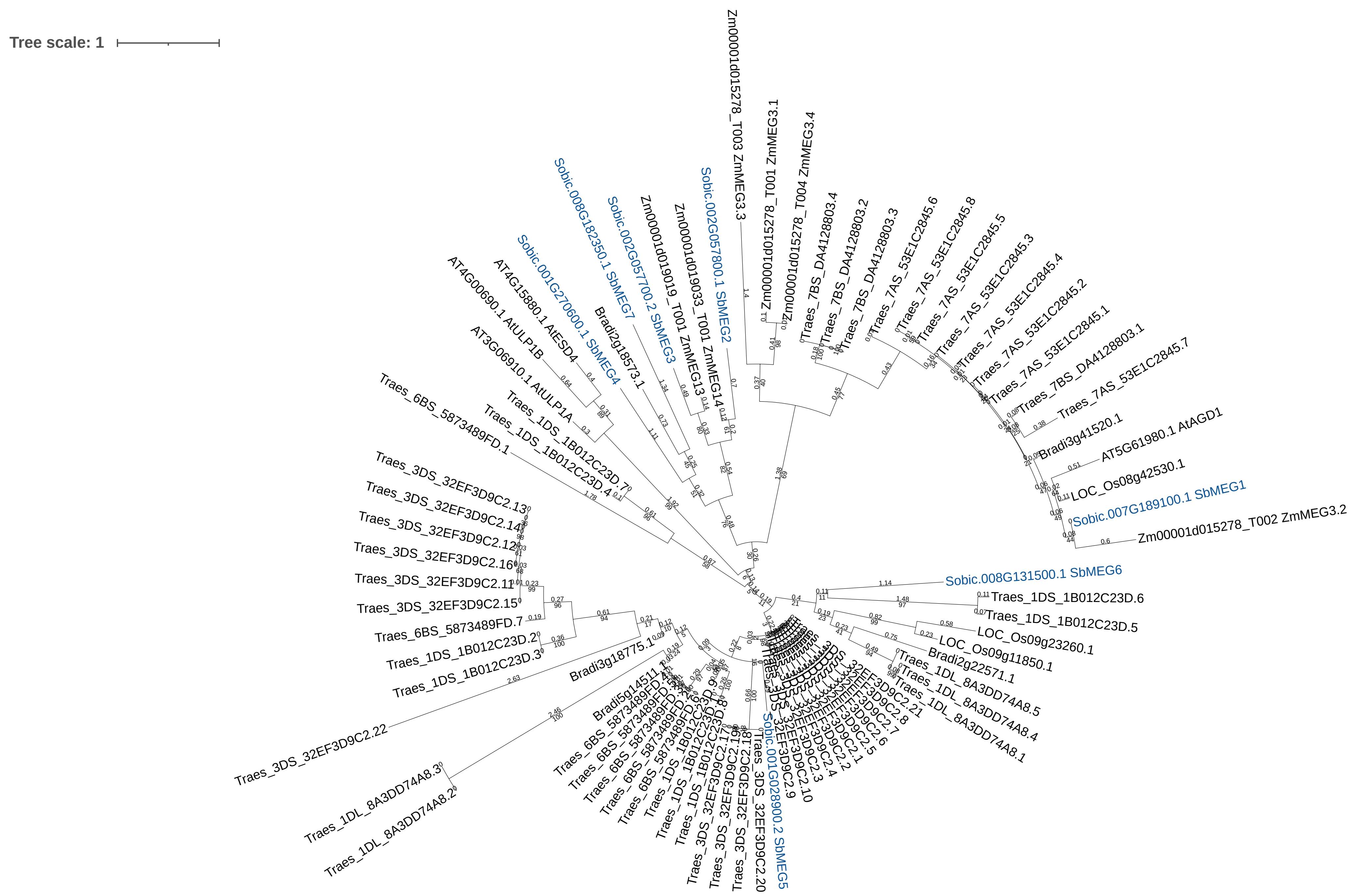


S-Figure 13. Maximum likelihood phylogenetic analysis of MEG SSP sequences in multiple plant species. The phylogeny includes sequences from *Arabidopsis thaliana* TAIR10 (At), *Oryza sativa* v7.0 (Os), *Brachypodium distachyon* v3.2 (Bd), *Triticum aestivum* v2.2 (Ta), *Zea mays* RefGen_V4 (Zm), and *Sorghum bicolor* v3.1.1 (Sb). Sorghum genes encoding SSPs are highlighted in blue. Genes encoding Sorghum SSPs are named based on their phylogenetic order in S-Figure 31. Genes encoding SSPs in other species are named based on published nomenclature. The tree was midpoint rooted. The scale bar represents the average number of amino acid substitutions per site. Trees were annotated in Dendroscope3 and visualized using iTOL v7.0.

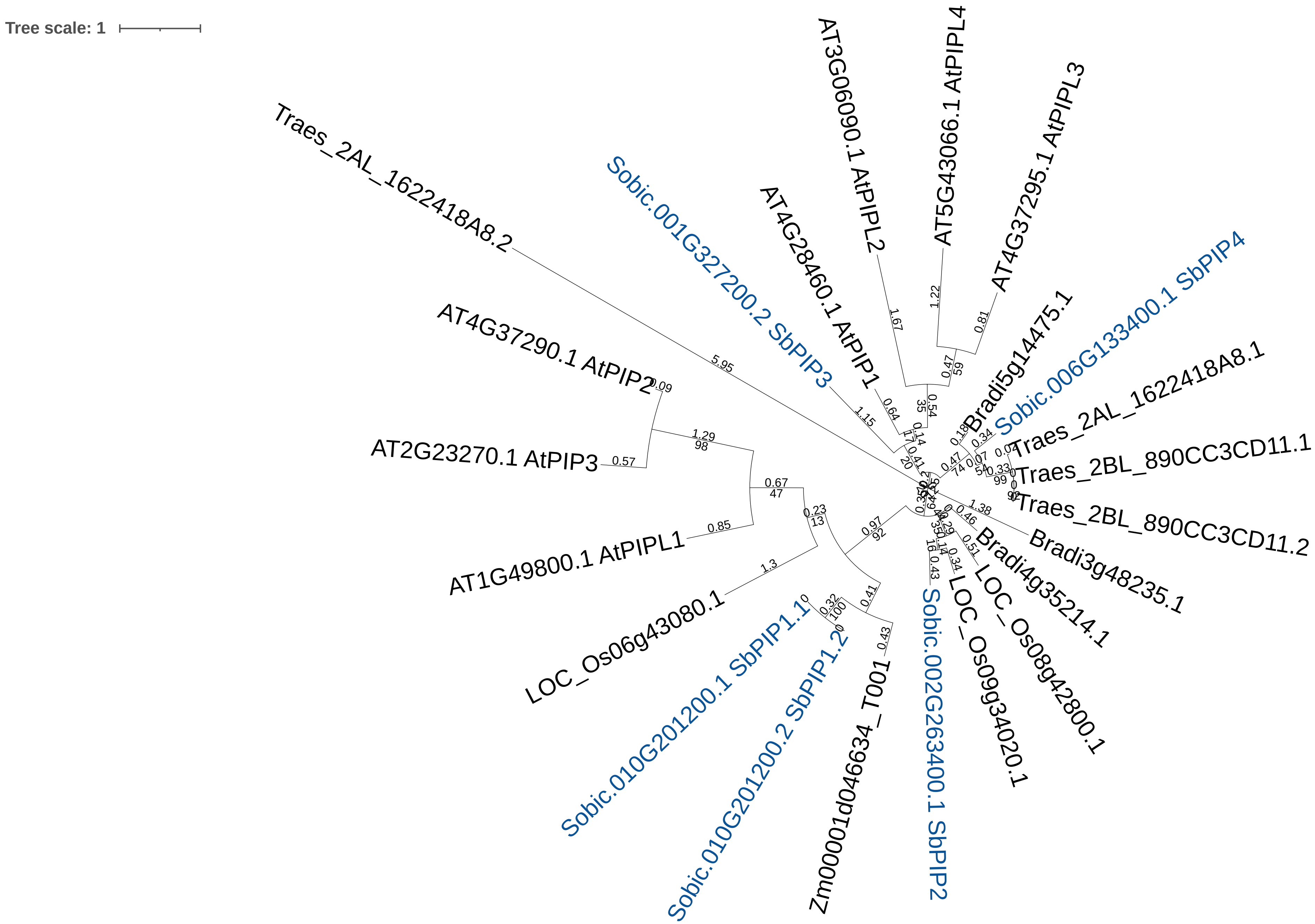


S-Figure 14. Maximum likelihood phylogenetic analysis of PIP SSP sequences in multiple plant species. The phylogeny includes sequences from *Arabidopsis thaliana* TAIR10 (At), *Oryza sativa* v7.0 (Os), *Brachypodium distachyon* v3.2 (Bd), *Triticum aestivum* v2.2 (Ta), *Zea mays* RefGen_V4 (Zm), and *Sorghum bicolor* v3.1.1 (Sb). Sorghum genes encoding SSPs are highlighted in blue. Genes encoding Sorghum SSPs are named based on their phylogenetic order in S-Figure 32. Genes encoding SSPs in other species are named based on published nomenclature. The tree was midpoint rooted. The scale bar represents the average number of amino acid substitutions per site. Trees were annotated in Dendroscope3 and visualized using iTOL v7.0.

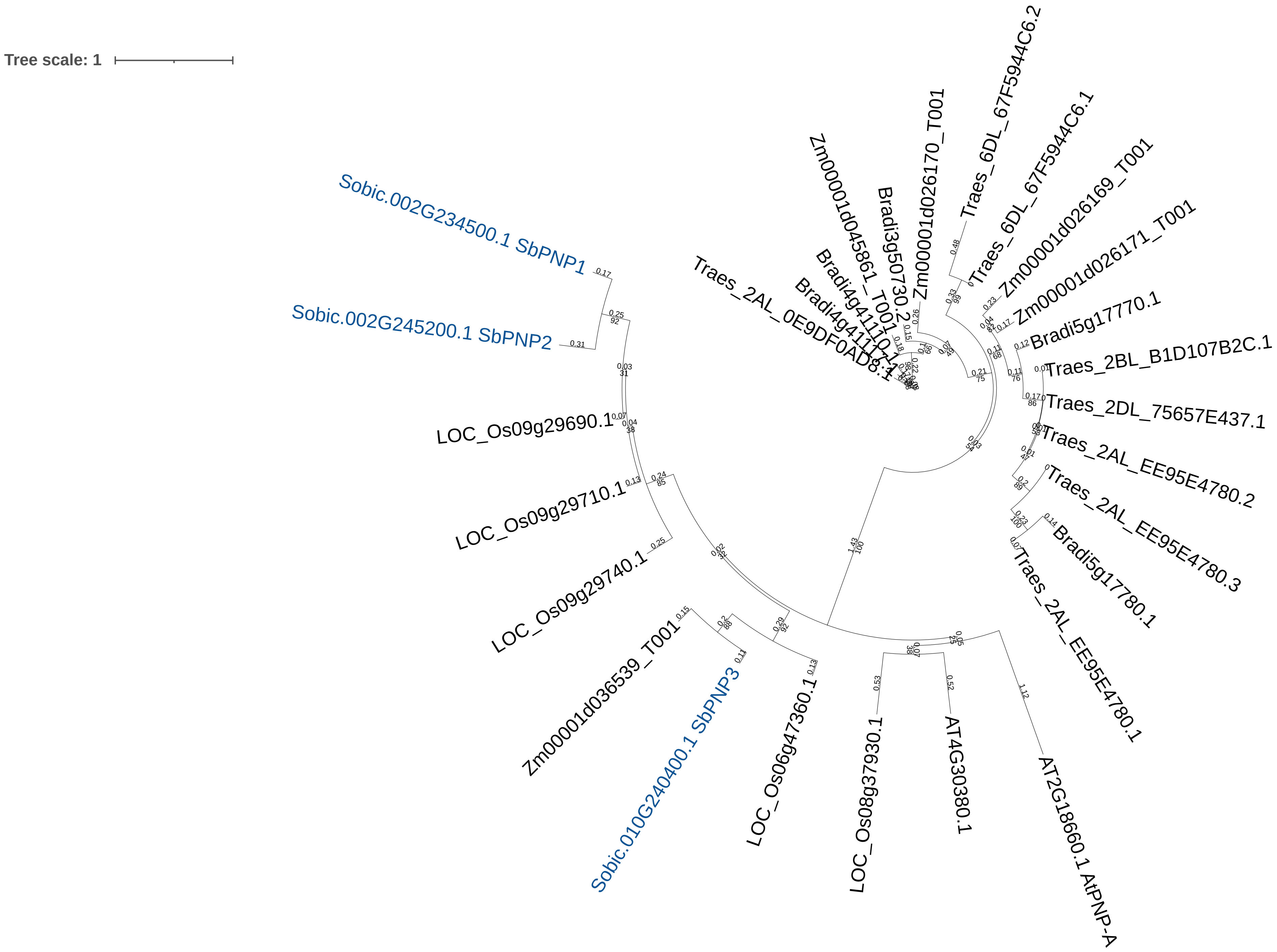


S-Figure 15. Maximum likelihood phylogenetic analysis of PNP SSP sequences in multiple plant species. The phylogeny includes sequences from *Arabidopsis thaliana* TAIR10 (At), *Oryza sativa* v7.0 (Os), *Brachypodium distachyon* v3.2 (Bd), *Triticum aestivum* v2.2 (Ta), *Zea mays* RefGen_V4 (Zm), and *Sorghum bicolor* v3.1.1 (Sb). Sorghum genes encoding SSPs are highlighted in blue. Genes encoding Sorghum SSPs are named based on their phylogenetic order in S-Figure 33. Genes encoding SSPs in other species are named based on published nomenclature.
The tree was midpoint rooted. The scale bar represents the average number of amino acid substitutions per site.
Trees were annotated in Dendroscope3 and visualized using iTOL v7.0.

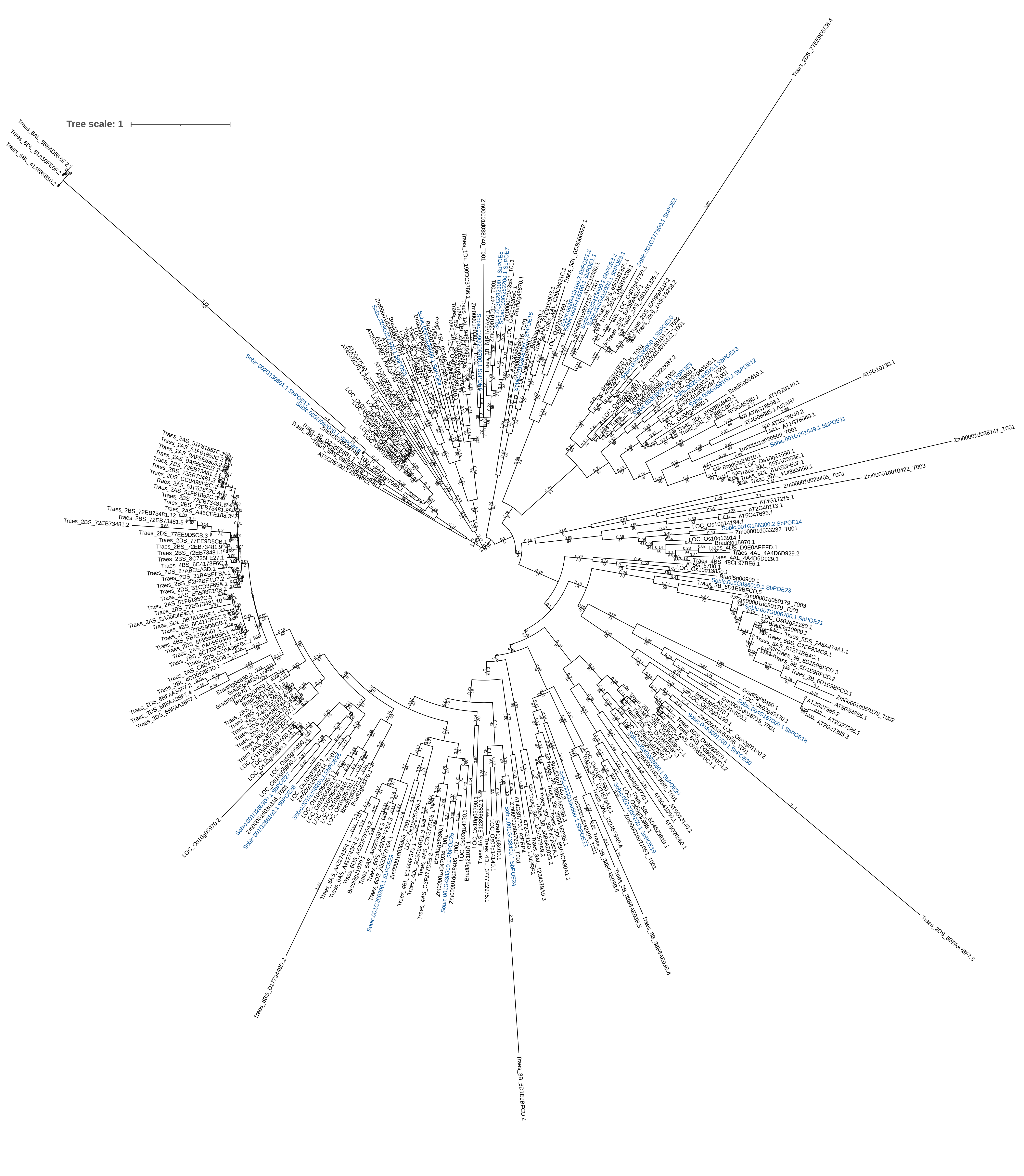


S-Figure 16. Maximum likelihood phylogenetic analysis of POE SSP sequences in multiple plant species. The phylogeny includes sequences from *Arabidopsis thaliana* TAIR10 (At), *Oryza sativa* v7.0 (Os), *Brachypodium distachyon* v3.2 (Bd), *Triticum aestivum* v2.2 (Ta), *Zea mays* RefGen_V4 (Zm), and *Sorghum bicolor* v3.1.1 (Sb). Sorghum genes encoding SSPs are highlighted in blue. Genes encoding Sorghum SSPs are named based on their phylogenetic order in S-Figure 34. Genes encoding SSPs in other species are named based on published nomenclature. The tree was midpoint rooted. The scale bar represents the average number of amino acid substitutions per site. Trees were annotated in Dendroscope3 and visualized using iTOL v7.0.

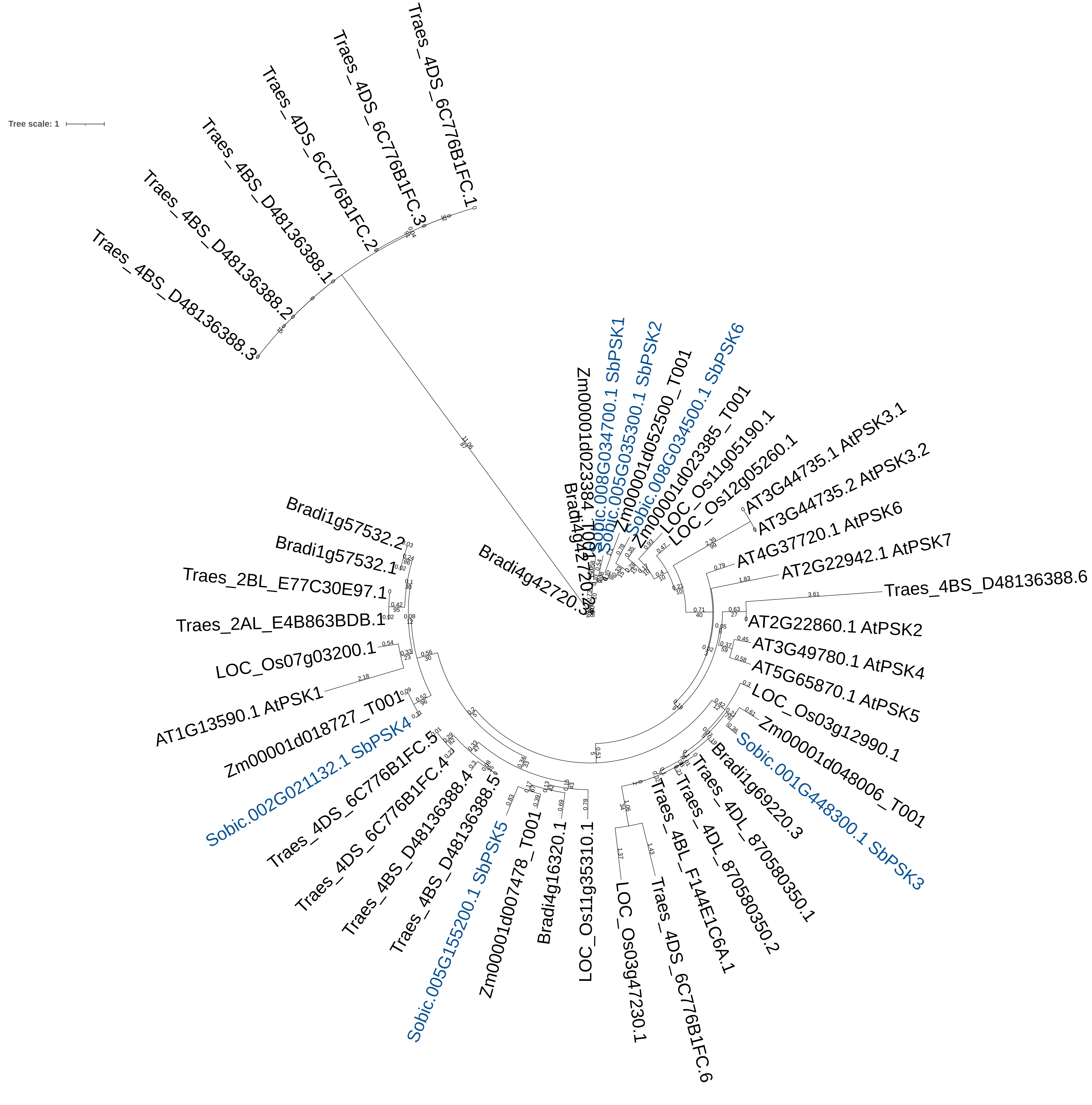


S-Figure 17. Maximum likelihood phylogenetic analysis of PSK SSP sequences in multiple plant species. The phylogeny includes sequences from *Arabidopsis thaliana* TAIR10 (At), *Oryza sativa* v7.0 (Os), *Brachypodium distachyon* v3.2 (Bd), *Triticum aestivum* v2.2 (Ta), *Zea mays* RefGen_V4 (Zm), and *Sorghum bicolor* v3.1.1 (Sb). Sorghum genes encoding SSPs are highlighted in blue. Genes encoding Sorghum SSPs are named based on their phylogenetic order in S-Figure 35. Genes encoding SSPs in other species are named based on published nomenclature. The tree was midpoint rooted. The scale bar represents the average number of amino acid substitutions per site.
Trees were annotated in Dendroscope3 and visualized using iTOL v7.0.

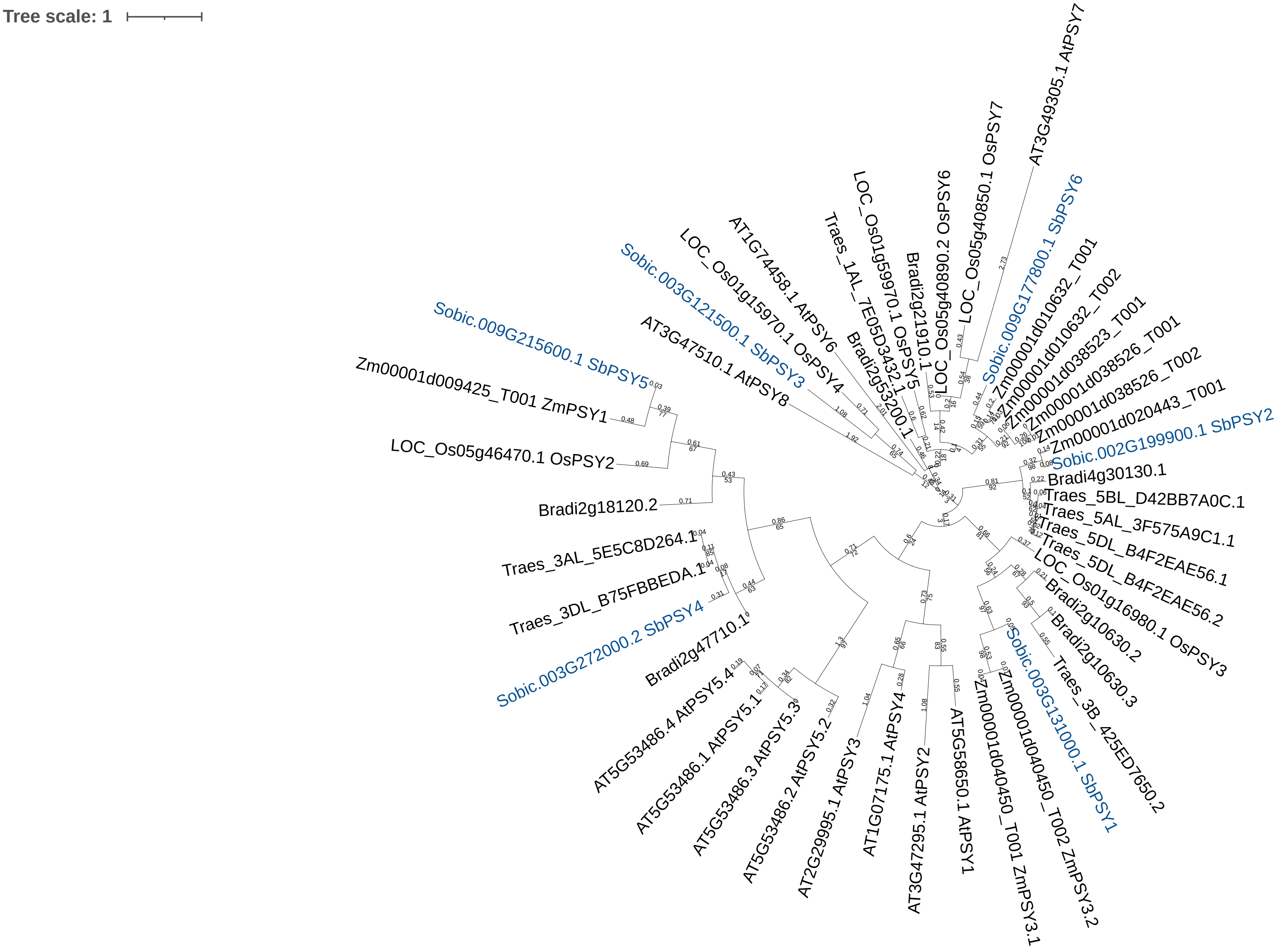


S-Figure 18. Maximum likelihood phylogenetic analysis of PSY SSP sequences in multiple plant species. The phylogeny includes sequences from *Arabidopsis thaliana* TAIR10 (At), *Oryza sativa* v7.0 (Os), *Brachypodium distachyon* v3.2 (Bd), *Triticum aestivum* v2.2 (Ta), *Zea mays* RefGen_V4 (Zm), and *Sorghum bicolor* v3.1.1 (Sb). Sorghum genes encoding SSPs are highlighted in blue. Genes encoding Sorghum SSPs are named based on their phylogenetic order in S-Figure 36. Genes encoding SSPs in other species are named based on published nomenclature. The tree was midpoint rooted. The scale bar represents the average number of amino acid substitutions per site. Trees were annotated in Dendroscope3 and visualized using iTOL v7.0.

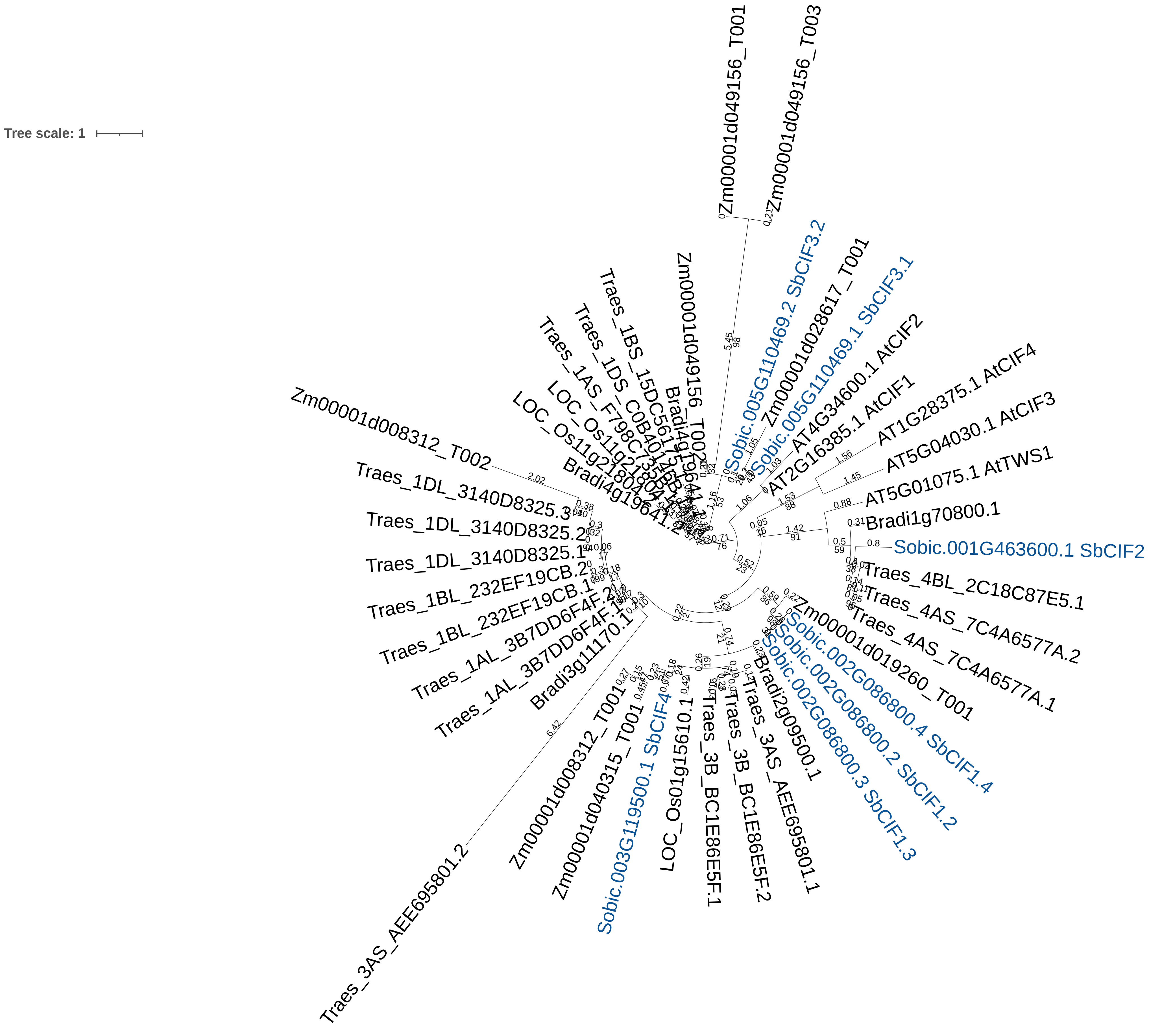


S-Figure 19. Maximum likelihood phylogenetic analysis of TPD SSP sequences in multiple plant species. The phylogeny includes sequences from *Arabidopsis thaliana* TAIR10 (At), *Oryza sativa* v7.0 (Os), *Brachypodium distachyon* v3.2 (Bd), *Triticum aestivum* v2.2 (Ta), *Zea mays* RefGen_V4 (Zm), and *Sorghum bicolor* v3.1.1 (Sb). Sorghum genes encoding SSPs are highlighted in blue. Genes encoding Sorghum SSPs are named based on their phylogenetic order in S-Figure 37. Genes encoding SSPs in other species are named based on published nomenclature. The tree was midpoint rooted. The scale bar represents the average number of amino acid substitutions per site. Trees were annotated in Dendroscope3 and visualized using iTOL v7.0.

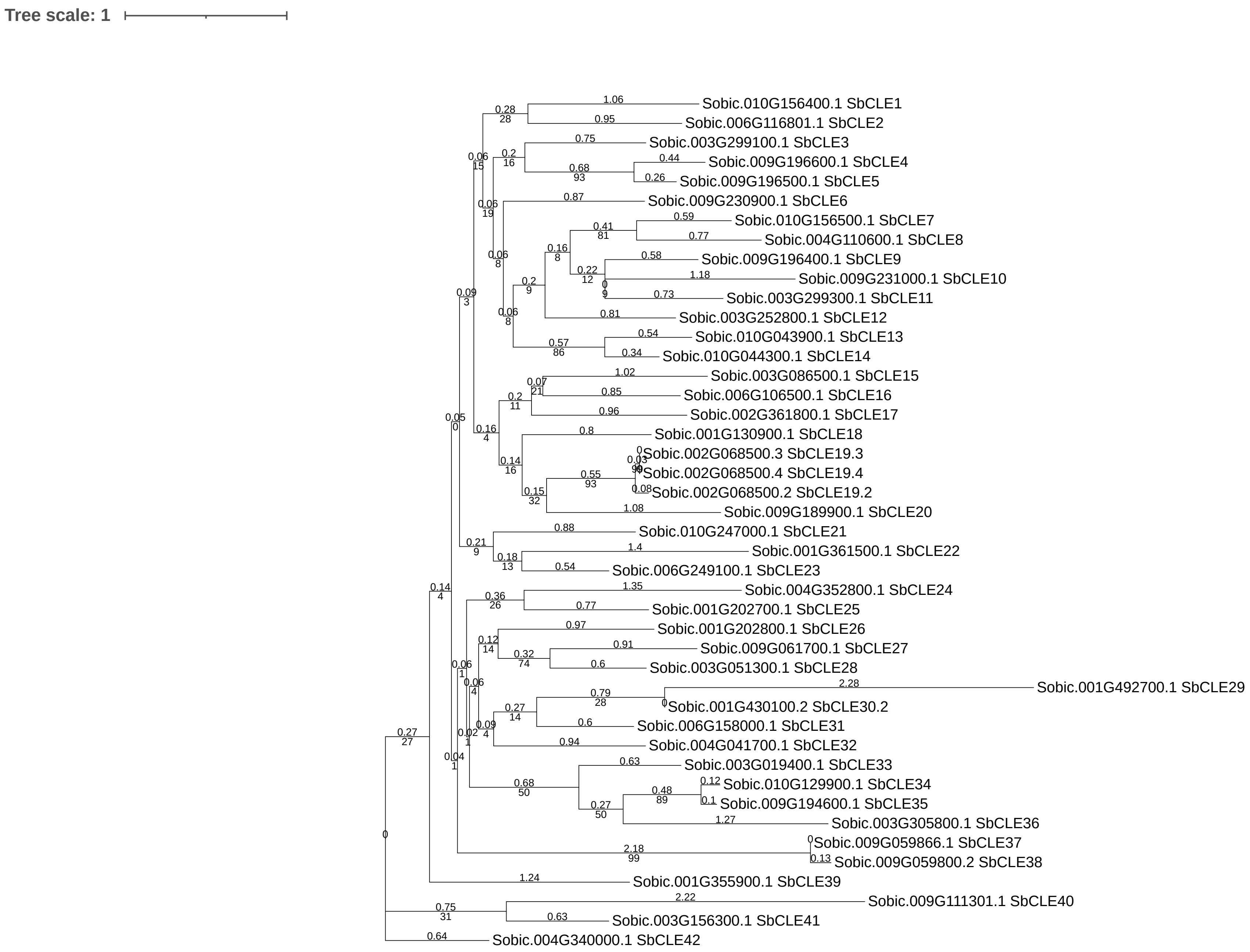


S-Figure 20. Maximum likelihood phylogenetic analysis of CLE SSP sequences for *Sorghum bicolor*. The scale bar represents the average number of amino acid substitutions per site. Tree was annotated and visualized in Dendroscope3 and iTOL v7.0.

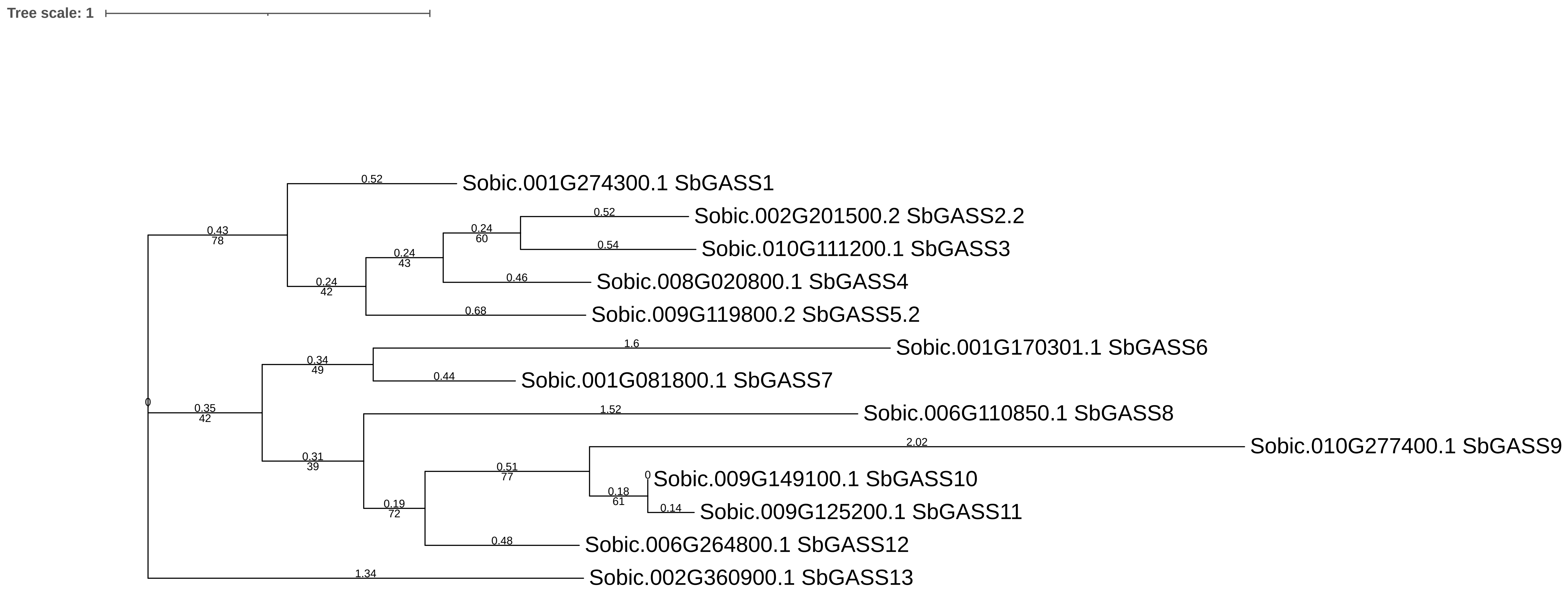


S-Figure 21. Maximum likelihood phylogenetic analysis of GASS SSP sequences for *Sorghum bicolor*. The scale bar represents the average number of amino acid substitutions per site. Tree was annotated and visualized in Dendroscope3 and iTOL v7.0.

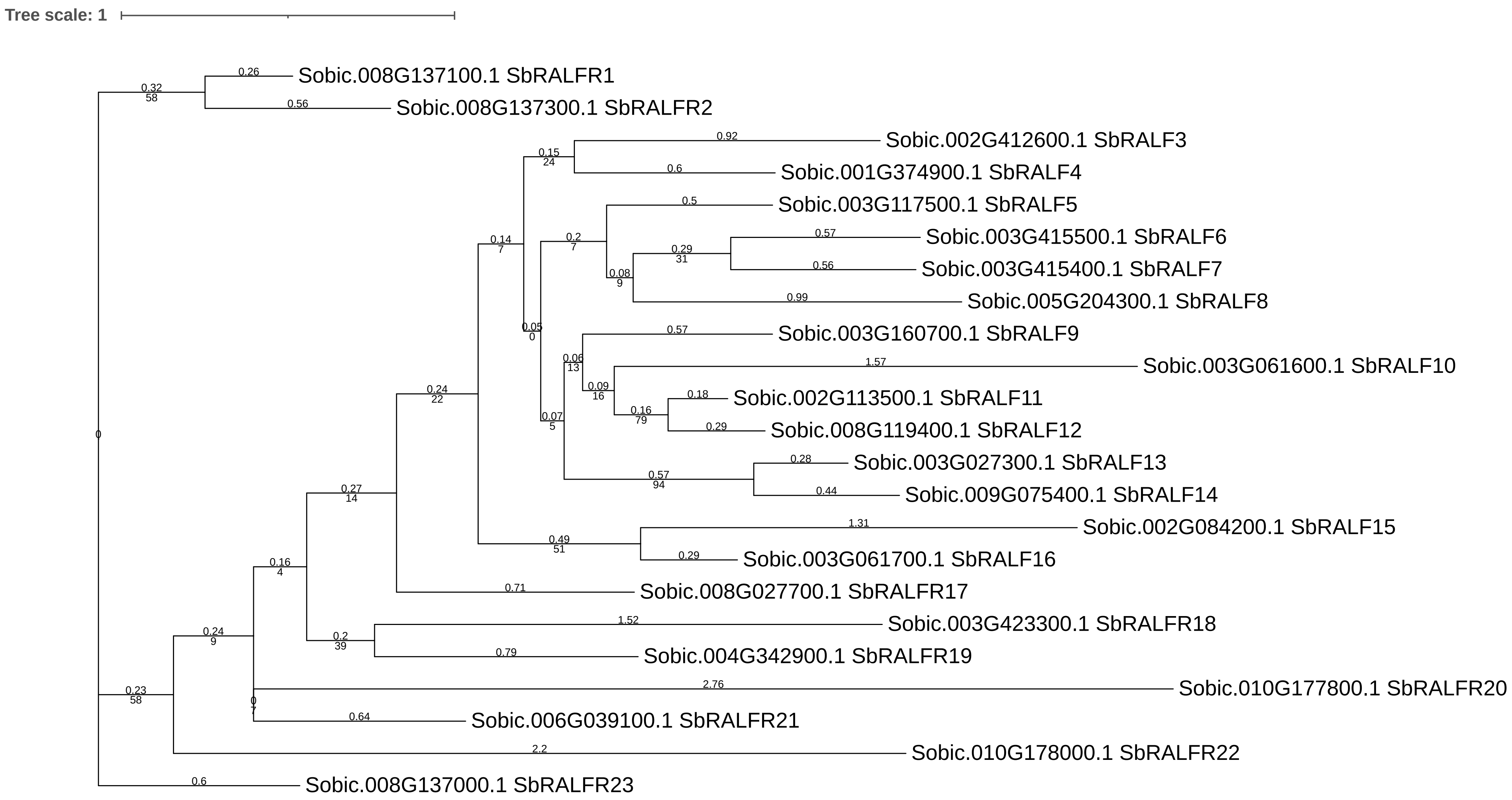


S-Figure 22. Maximum likelihood phylogenetic analysis of RALF SSP sequences for *Sorghum bicolor*. The scale bar represents the average number of amino acid substitutions per site. Tree was annotated and visualized in Dendroscope3 and iTOL v7.0.

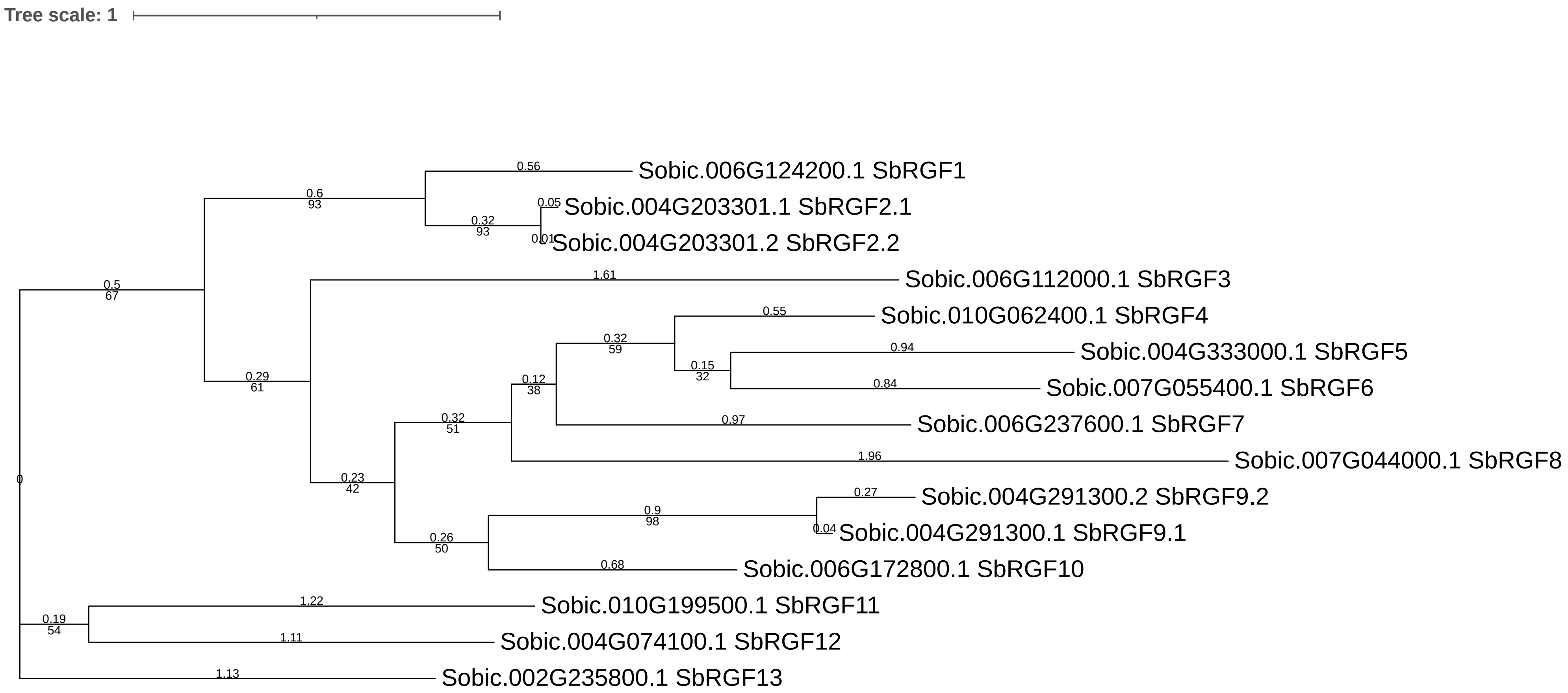


S-Figure 23. Maximum likelihood phylogenetic analysis of RGF SSP sequences for *Sorghum bicolor*. The scale bar represents the average number of amino acid substitutions per site. Tree was annotated and visualized in Dendroscope3 and iTOL v7.0.

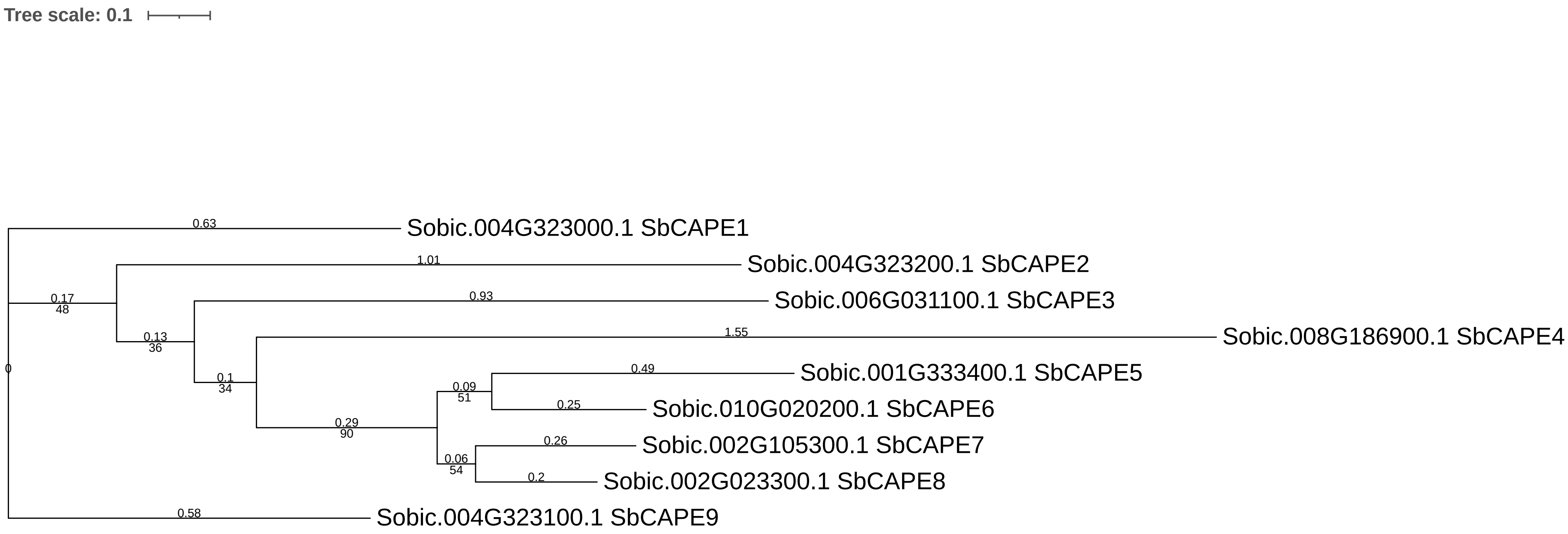


S-Figure 24. Maximum likelihood phylogenetic analysis of CAPE SSP sequences for *Sorghum bicolor*. The scale bar represents the average number of amino acid substitutions per site. Tree was annotated and visualized in Dendroscope3 and iTol v7.0.

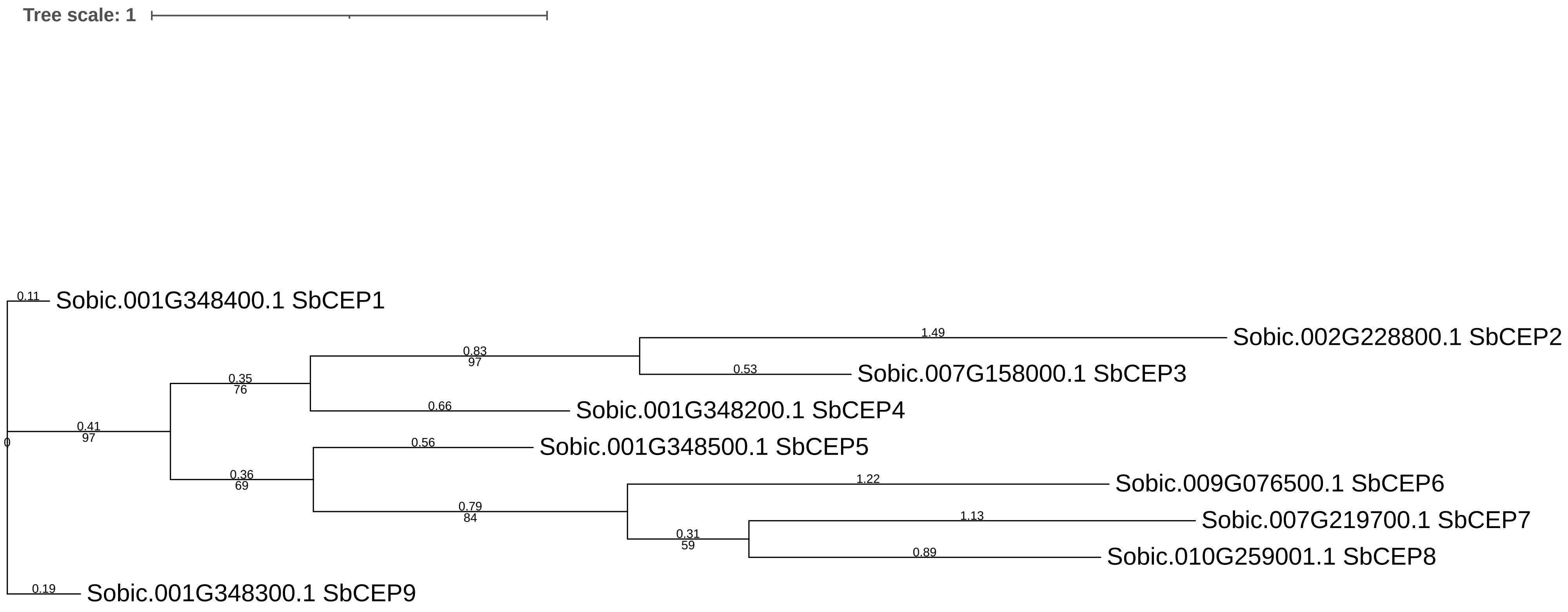


S-Figure 25. Maximum likelihood phylogenetic analysis of CEP SSP sequences for *Sorghum bicolor*. The scale bar represents the average number of amino acid substitutions per site. Tree was annotated and visualized in Dendroscope3 and iTOL v7.0.

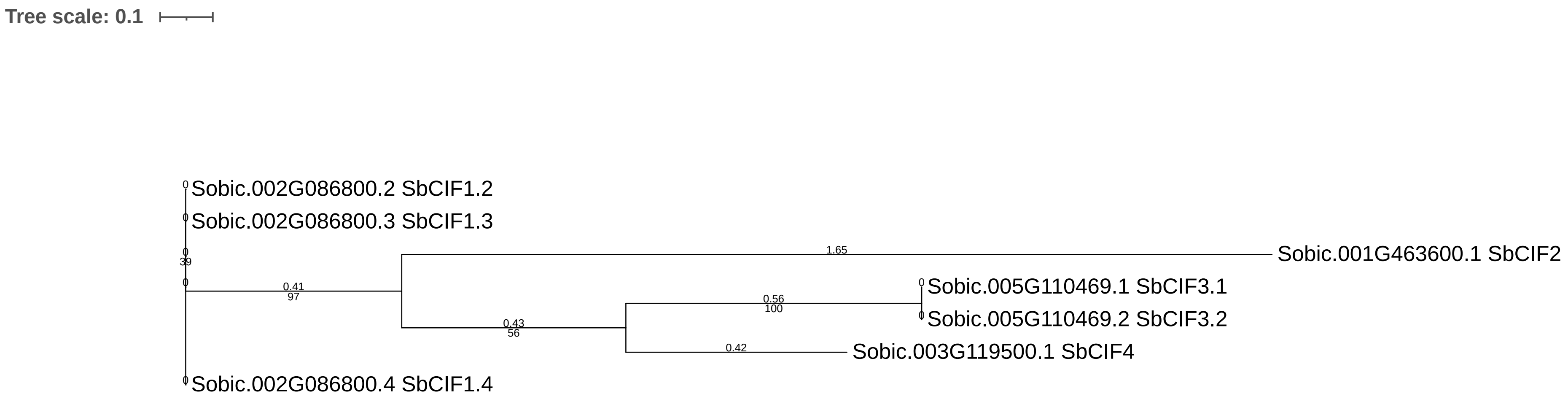


S-Figure 26. Maximum likelihood phylogenetic analysis of CIF SSP sequences for *Sorghum bicolor*. The scale bar represents the average number of amino acid substitutions per site. Tree was annotated and visualized in Dendroscope3 and iTol v7.0.

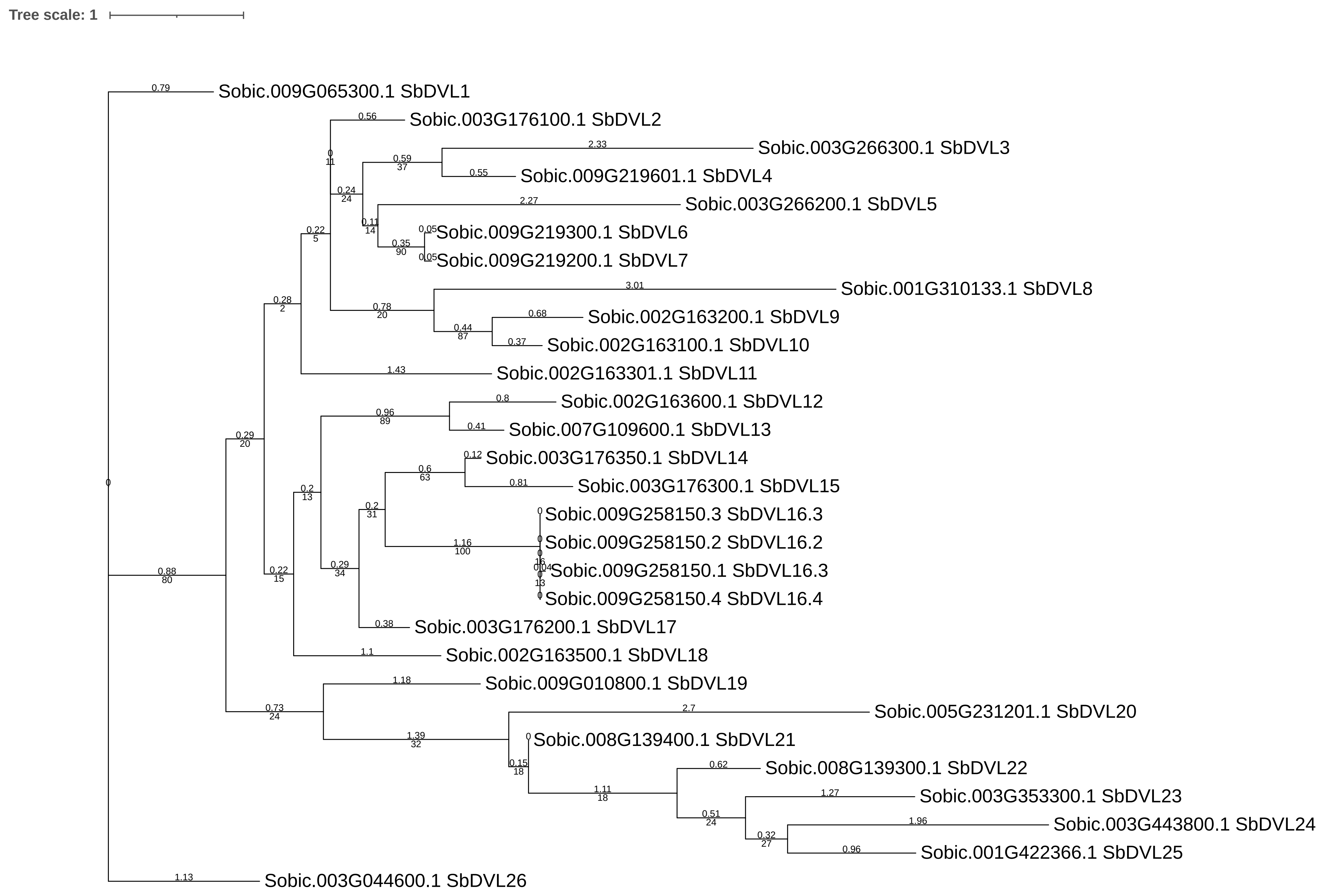


S-Figure 27. Maximum likelihood phylogenetic analysis of DVL SSP sequences for Sorghum bicolor. The scale bar represents the average number of amino acid substitutions per site. Tree was annotated and visualized in Dendroscope3 and iTol v7.0.

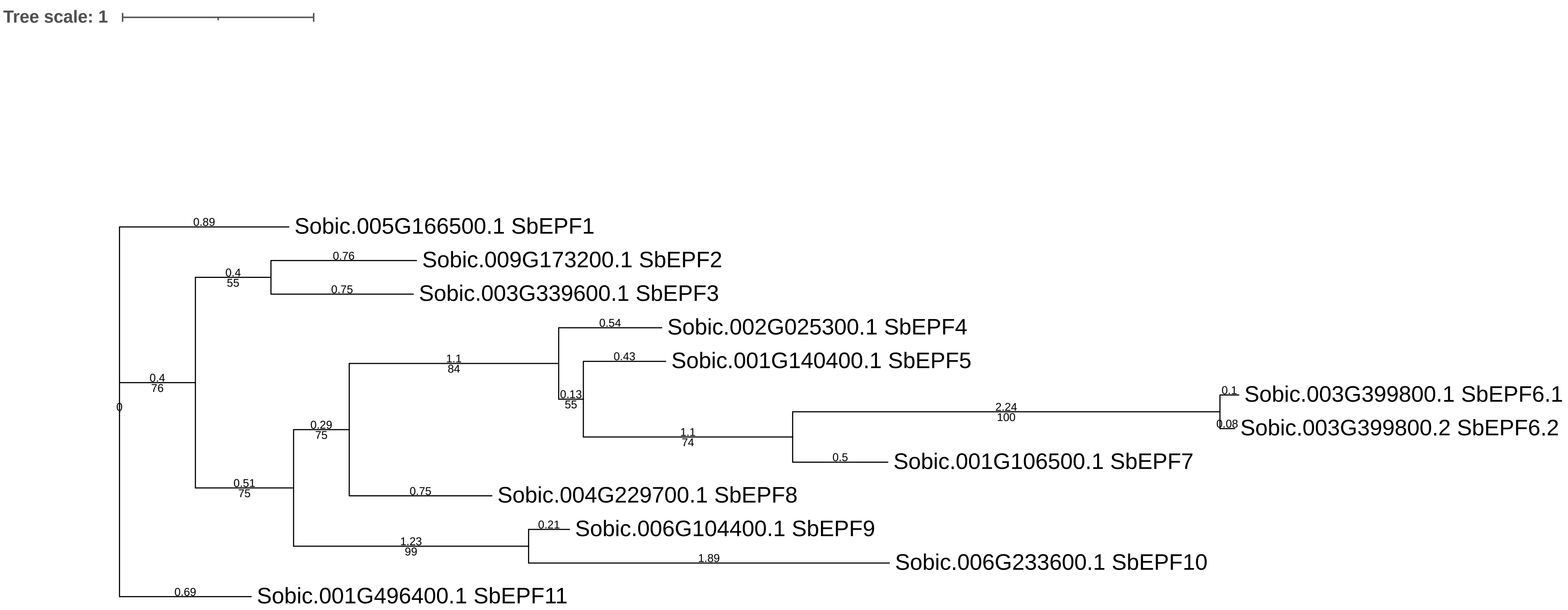


S-Figure 28. Maximum likelihood phylogenetic analysis of EPF SSP sequences for *Sorghum bicolor*. The scale bar represents the average number of amino acid substitutions per site. Tree was annotated and visualized in Dendroscope3 and iTol v7.0.

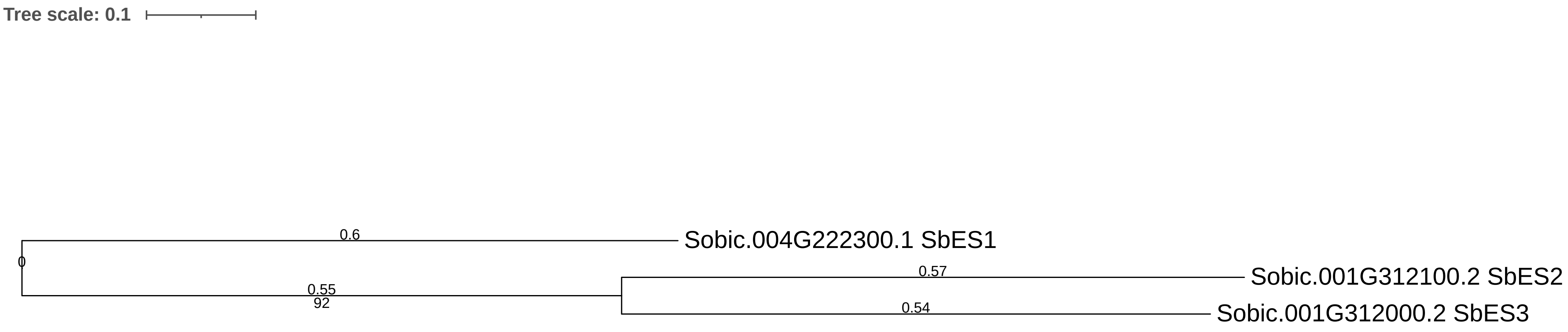


S-Figure 29. Maximum likelihood phylogenetic analysis of ES SSP sequences for *Sorghum bicolor*. The scale bar represents the average number of amino acid substitutions per site. Tree was annotated and visualized in Dendroscope3 and iTOL v7.0.

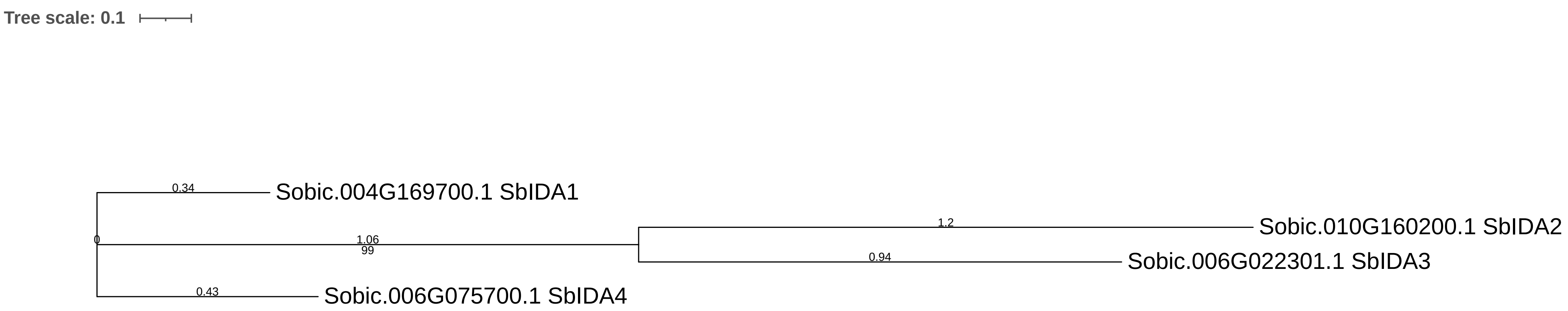


S-Figure 30. Maximum likelihood phylogenetic analysis of IDA SSP sequences for *Sorghum bicolor*. The scale bar represents the average number of amino acid substitutions per site. Tree was annotated and visualized in Dendroscope3 and iTOL v7.0.

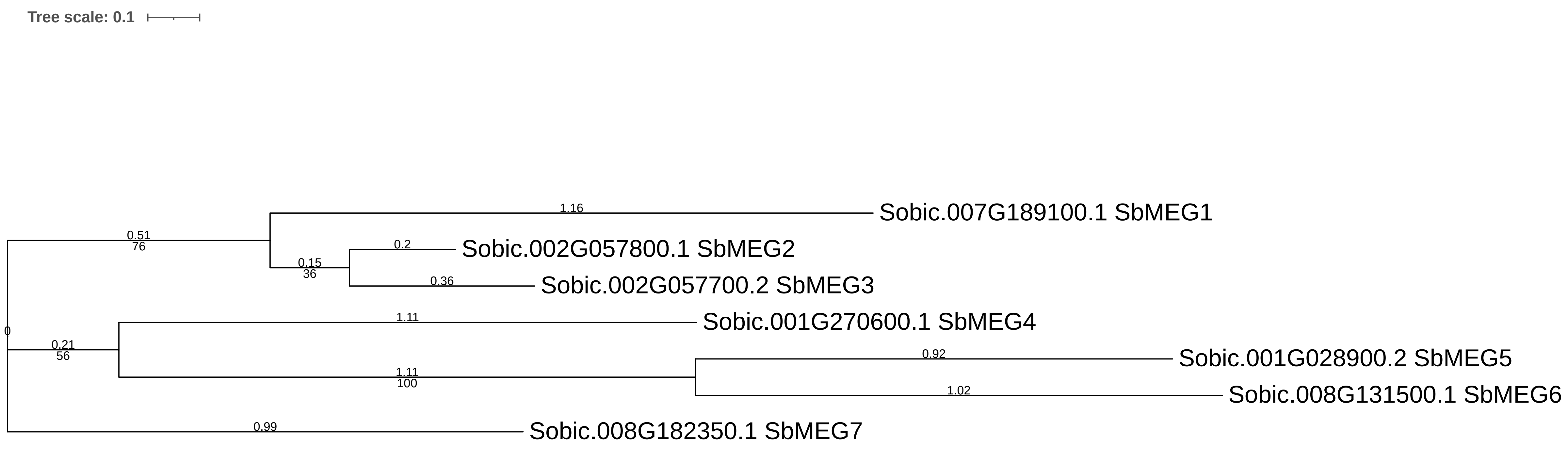


S-Figure 31. Maximum likelihood phylogenetic analysis of MEG SSP sequences for *Sorghum bicolor*. The scale bar represents the average number of amino acid substitutions per site. Tree was annotated and visualized in Dendroscope3 and iTOL v7.0.

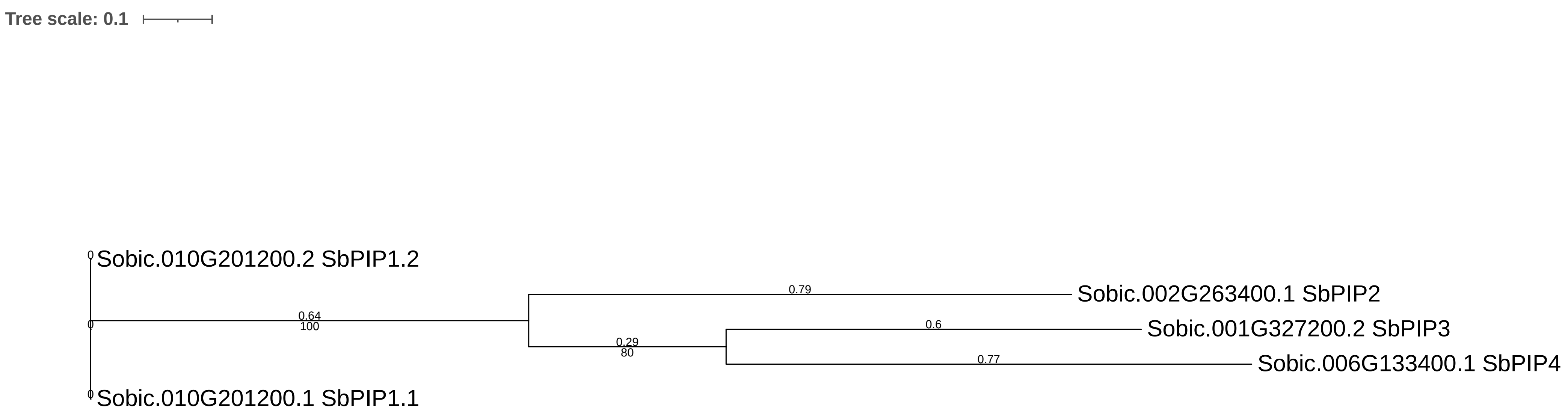


S-Figure 32. Maximum likelihood phylogenetic analysis of PIP SSP sequences for *Sorghum bicolor*. The scale bar represents the average number of amino acid substitutions per site. Tree was annotated and visualized in Dendroscope3 and iTOL v7.0.

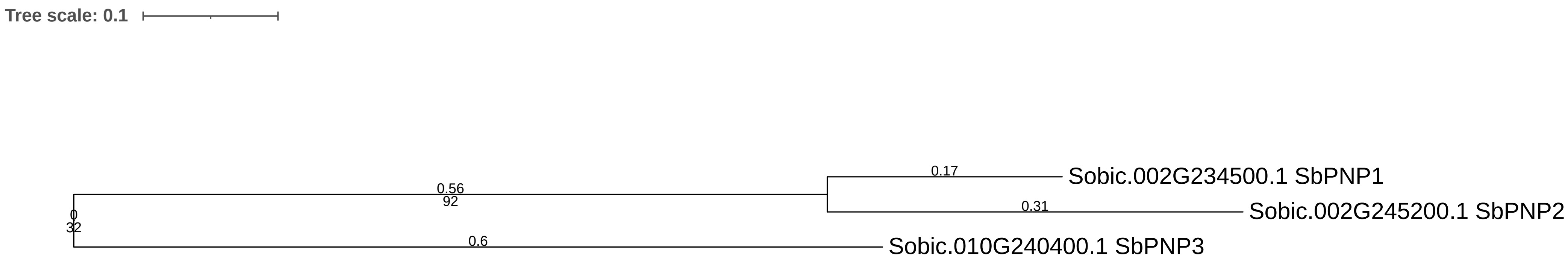


S-Figure 33. Maximum likelihood phylogenetic analysis of PNP SSP sequences for *Sorghum bicolor*. The scale bar represents the average number of amino acid substitutions per site. Tree was annotated and visualized in Dendroscope3 and iTOL v7.0.

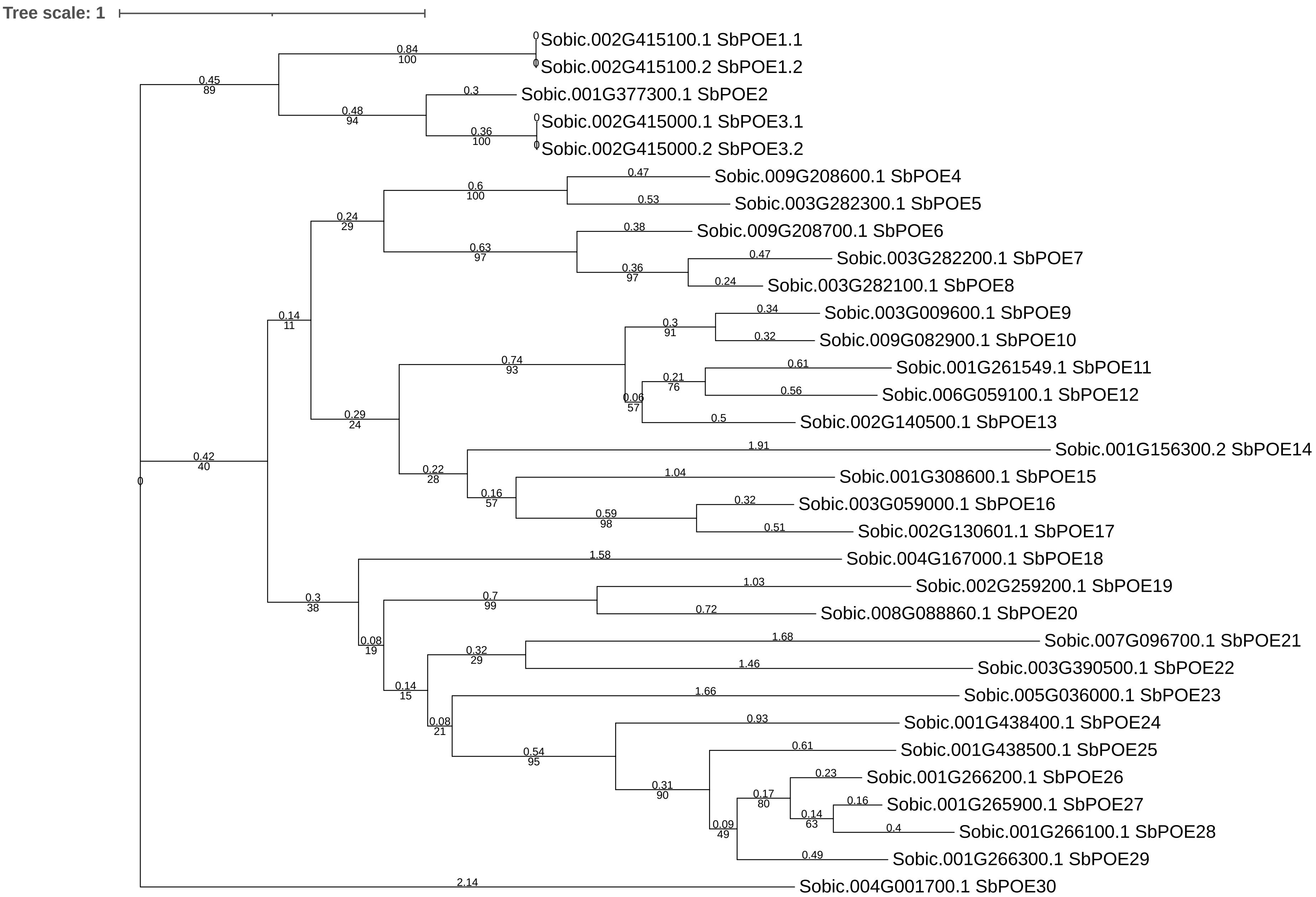


S-Figure 34. Maximum likelihood phylogenetic analysis of POE SSP sequences for *Sorghum bicolor*. The scale bar represents the average number of amino acid substitutions per site. Tree was annotated and visualized in Dendroscope3 and iTOL v7.0.

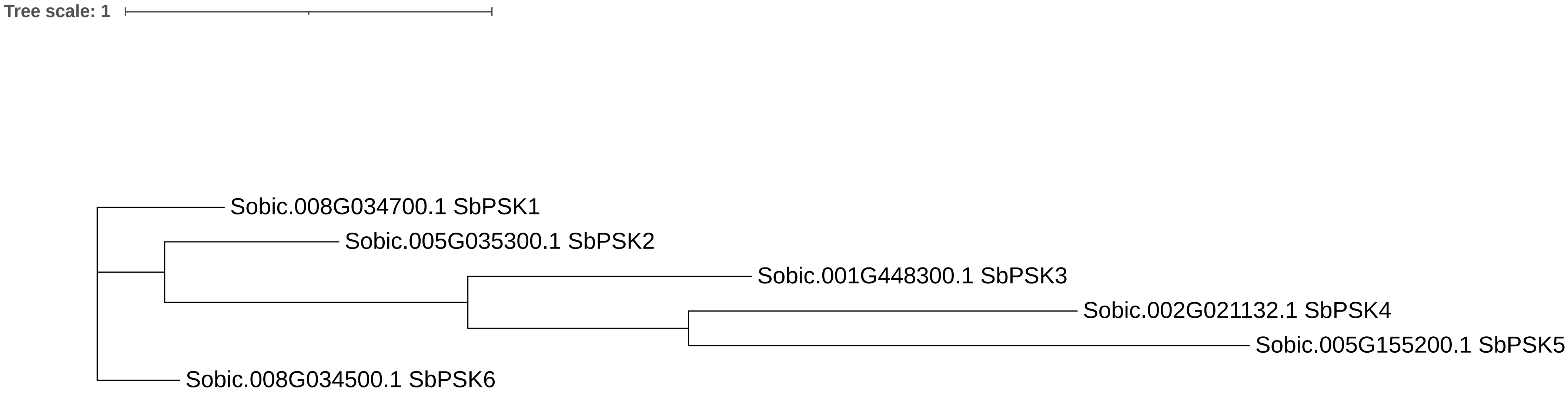


S-Figure 35. Maximum likelihood phylogenetic analysis of PSK SSP sequences for *Sorghum bicolor*. The scale bar represents the average number of amino acid substitutions per site. Tree was annotated and visualized in Dendroscope3 and iTOL v7.0.

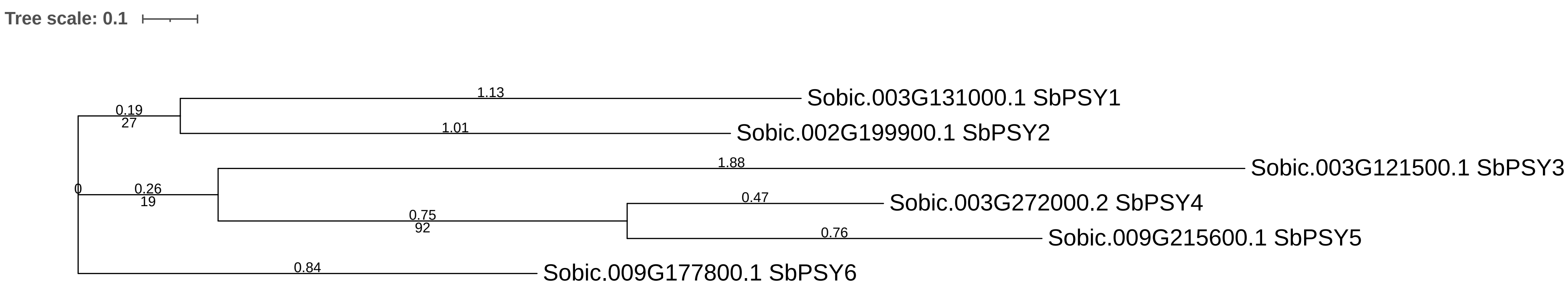


S-Figure 36. Maximum likelihood phylogenetic analysis of PSY SSP sequences for *Sorghum bicolor*. The scale bar represents the average number of amino acid substitutions per site. Tree was annotated and visualized in Dendroscope3 and iTOL v7.0.

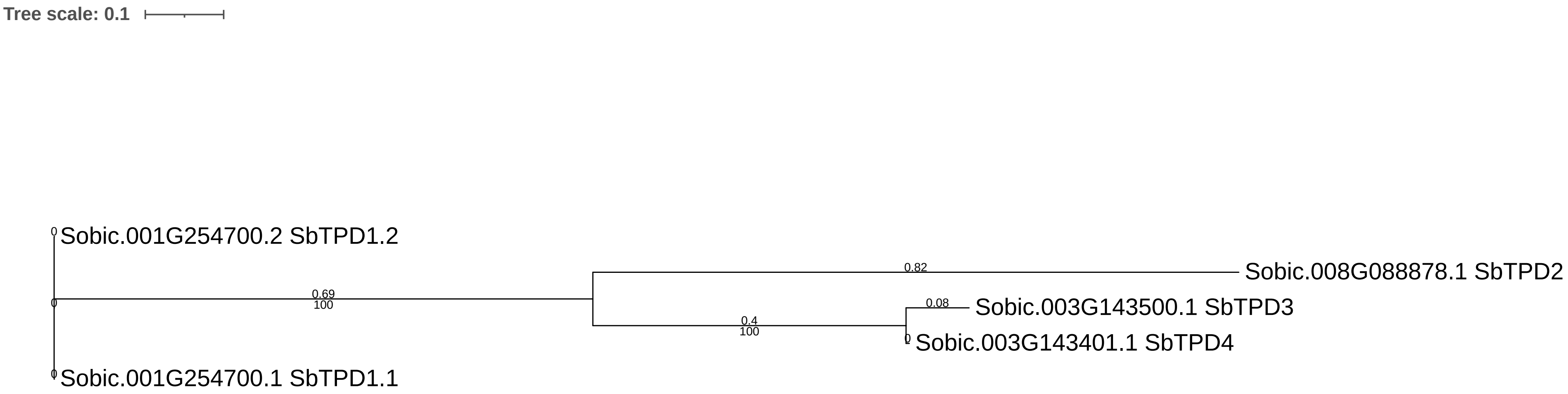


S-Figure 37. Maximum likelihood phylogenetic analysis of TPD SSP sequences for *Sorghum bicolor*. The scale bar represents the average number of amino acid substitutions per site. Tree was annotated and visualized in Dendroscope3 and iTOL v7.0.

| Gene name | Transcript ID | Max | τ |
|---|---|---|---|
| SbCAPE2 | Sobic.004G323200.1 | 39 | 0.83 |
| SbCAPE3 | Sobic.006G031100.1 | 27 | 0.69 |
| SbCAPE4 | Sobic.008G186900.1 | 155 | 1.00 |
| SbCAPE5 | Sobic.001G333400.1 | 113 | 0.98 |
| SbCAPE6 | Sobic.010G020200.1 | 306 | 0.83 |
| SbCAPE8 | Sobic.002G023300.1 | 803 | 0.97 |
| SbCAPE9 | Sobic.004G323100.1 | 27 | 0.70 |
| SbCEP1 | Sobic.001G348400.1 | 33 | 0.95 |
| SbCEP2 | Sobic.002G228800.1 | 13 | 0.99 |
| SbCEP3 | Sobic.007G158000.1 | 57 | 0.96 |
| SbCEP4 | Sobic.001G348200.1 | 9 | 0.95 |
| SbCEP5 | Sobic.001G348500.1 | 6 | 0.93 |
| SbCEP6 | Sobic.009G076500.1 | 25 | 0.94 |
| SbCEP7 | Sobic.007G219700.1 | 58 | 1.00 |
| SbCEP8 | Sobic.010G259001.1 | 82 | 0.92 |
| SbCEP9 | Sobic.001G348300.1 | 7 | 0.98 |
| SbCIF1 | Sobic.002G086800.4 | 228 | 0.88 |
| SbCIF3 | Sobic.005G110469.1 | 395 | 0.88 |
| SbCIF4 | Sobic.003G119500.1 | 927 | 0.86 |
| SbCLE1 | Sobic.010G156400.1 | 19 | 1.00 |
| SbCLE3 | Sobic.003G299100.1 | 8 | 0.98 |
| SbCLE6 | Sobic.009G230900.1 | 56 | 0.95 |
| SbCLE7 | Sobic.010G156500.1 | 40 | 1.00 |
| SbCLE9 | Sobic.009G196400.1 | 88 | 0.99 |
| SbCLE10 | Sobic.009G231000.1 | 13 | 0.97 |
| SbCLE11 | Sobic.003G299300.1 | 11 | 0.99 |
| SbCLE12 | Sobic.003G252800.1 | 17 | 0.97 |
| SbCLE15 | Sobic.003G086500.1 | 7 | 0.39 |
| SbCLE18 | Sobic.001G130900.1 | 16 | 0.66 |
| SbCLE20 | Sobic.009G189900.1 | 7 | 0.59 |
| SbCLE23 | Sobic.006G249100.1 | 8 | 0.65 |
| SbCLE24 | Sobic.004G352800.1 | 19 | 0.48 |
| SbCLE25 | Sobic.001G202700.1 | 34 | 0.59 |
| SbCLE26 | Sobic.001G202800.1 | 16 | 0.60 |
| SbCLE27 | Sobic.009G061700.1 | 6 | 0.76 |
| SbCLE28 | Sobic.003G051300.1 | 17 | 0.89 |
| SbCLE31 | Sobic.006G158000.1 | 8 | 0.51 |
| SbCLE33 | Sobic.003G019400.1 | 7 | 0.67 |
| SbCLE37 | Sobic.009G059866.1 | 7 | 1.00 |
| SbCLE38 | Sobic.009G059800.2 | 49 | 1.00 |
| SbCLE41 | Sobic.003G156300.1 | 11 | 0.56 |
| SbCLE42 | Sobic.004G340000.1 | 65 | 0.89 |

| Gene name | Transcript ID | Max | τ |
|---|---|---|---|
| SbDVL1 | Sobic.009G065300.1 | 69 | 0.81 |
| SbDVL2 | Sobic.003G176100.1 | 35 | 0.85 |
| SbDVL3 | Sobic.003G266300.1 | 11 | 0.85 |
| SbDVL6 | Sobic.009G219300.1 | 6 | 0.98 |
| SbDVL7 | Sobic.009G219200.1 | 33 | 0.94 |
| SbDVL9 | Sobic.002G163200.1 | 31 | 0.96 |
| SbDVL11 | Sobic.002G163301.1 | 21 | 0.87 |
| SbDVL12 | Sobic.002G163600.1 | 7 | 0.34 |
| SbDVL13 | Sobic.007G109600.1 | 15 | 0.87 |
| SbDVL14 | Sobic.003G176350.1 | 7 | 0.70 |
| SbDVL16 | Sobic.009G258150.2 | 26 | 0.89 |
| SbDVL17 | Sobic.003G176200.1 | 865 | 0.85 |
| SbDVL18 | Sobic.002G163500.1 | 17 | 0.84 |
| SbDVL21 | Sobic.008G139400.1 | 36 | 0.59 |
| SbDVL23 | Sobic.003G353300.1 | 54 | 0.77 |
| SbDVL24 | Sobic.003G443800.1 | 34 | 0.92 |
| SbDVL25 | Sobic.001G422366.1 | 222 | 0.70 |
| SbDVL26 | Sobic.003G044600.1 | 64 | 0.76 |
| SbECL2 | Sobic.008G049300.1 | 3 | 0.90 |
| SbEPF1 | Sobic.005G166500.1 | 58 | 0.85 |
| SbEPF2 | Sobic.009G173200.1 | 24 | 0.63 |
| SbEPF3 | Sobic.003G339600.1 | 16 | 0.88 |
| SbEPF4 | Sobic.002G025300.1 | 129 | 0.92 |
| SbEPF5 | Sobic.001G140400.1 | 18 | 0.45 |
| SbEPF6 | Sobic.003G399800.1 | 69 | 0.85 |
| SbEPF7 | Sobic.001G106500.1 | 32 | 0.77 |
| SbEPF8 | Sobic.004G229700.1 | 28 | 0.67 |
| SbEPF11 | Sobic.001G496400.1 | 68 | 0.81 |
| SbES1 | Sobic.004G222300.1 | 120 | 0.83 |
| SbIDA1 | Sobic.004G169700.1 | 26 | 0.31 |
| SbIDA4 | Sobic.006G075700.1 | 108 | 0.47 |
| SbGASS1 | Sobic.001G274300.1 | 26 | 1.00 |
| SbGASS2 | Sobic.002G201500.2 | 108 | 0.26 |
| SbGASS3 | Sobic.010G111200.1 | 865 | 0.57 |
| SbGASS4 | Sobic.008G020800.1 | 20 | 0.44 |
| SbGASS5 | Sobic.009G119800.2 | 8 | 0.69 |
| SbGASS6 | Sobic.001G170301.1 | 693 | 0.98 |
| SbGASS7 | Sobic.001G081800.1 | 628 | 0.65 |
| SbGASS8 | Sobic.006G110850.1 | 555 | 0.92 |
| SbGASS9 | Sobic.010G277400.1 | 159 | 0.71 |
| SbGASS10 | Sobic.009G149100.1 | 609 | 1.00 |
| SbGASS11 | Sobic.009G125200.1 | 93 | 1.00 |
| SbGASS12 | Sobic.006G264800.1 | 7 | 0.78 |
| SbGASS13 | Sobic.002G360900.1 | 108 | 0.69 |

| Gene name | Transcript ID | Max | τ |
|---|---|---|---|
| SbMEG1 | Sobic.007G189100.1 | 142 | 0.58 |
| SbMEG2 | Sobic.002G057800.1 | 6 | 0.99 |
| SbMEG5 | Sobic.001G028900.2 | 6 | 0.37 |
| SbMEG6 | Sobic.008G131500.1 | 25 | 0.57 |
| SbPIP1 | Sobic.010G201200.1 | 18 | 0.86 |
| SbPIP2 | Sobic.002G263400.1 | 23 | 0.49 |
| SbPIP4 | Sobic.006G133400.1 | 5 | 0.93 |
| SbPNP1 | Sobic.002G234500.1 | 1160 | 1.00 |
| SbPNP2 | Sobic.002G245200.1 | 1 | 0.94 |
| SbPNP3 | Sobic.010G240400.1 | 546 | 0.98 |
| SbPOE1 | Sobic.002G415100.1 | 17 | 0.94 |
| SbPOE2 | Sobic.001G377300.1 | 530 | 0.74 |
| SbPOE3 | Sobic.002G415000.1 | 69 | 0.66 |
| SbPOE4 | Sobic.009G208600.1 | 13 | 0.87 |
| SbPOE5 | Sobic.003G282300.1 | 35 | 0.99 |
| SbPOE6 | Sobic.009G208700.1 | 18 | 0.81 |
| SbPOE7 | Sobic.003G282200.1 | 9 | 1.00 |
| SbPOE9 | Sobic.003G009600.1 | 55 | 1.00 |
| SbPOE10 | Sobic.009G082900.1 | 49 | 1.00 |
| SbPOE11 | Sobic.001G261549.1 | 495 | 0.49 |
| SbPOE12 | Sobic.006G059100.1 | 335 | 1.00 |
| SbPOE13 | Sobic.002G140500.1 | 18 | 0.98 |
| SbPOE14 | Sobic.001G156300.2 | 147 | 0.57 |
| SbPOE15 | Sobic.001G308600.1 | 11 | 0.98 |
| SbPOE16 | Sobic.003G059000.1 | 25 | 1.00 |
| SbPOE19 | Sobic.002G259200.1 | 70 | 0.82 |
| SbPOE20 | Sobic.008G088860.1 | 60 | 0.59 |
| SbPOE21 | Sobic.007G096700.1 | 263 | 0.19 |
| SbPOE22 | Sobic.003G390500.1 | 139 | 0.97 |
| SbPOE23 | Sobic.005G036000.1 | 129 | 0.54 |
| SbPOE24 | Sobic.001G438400.1 | 11 | 0.78 |
| SbPOE25 | Sobic.001G438500.1 | 286 | 0.90 |
| SbPOE26 | Sobic.001G266200.1 | 648 | 0.71 |
| SbPOE27 | Sobic.001G265900.1 | 1373 | 0.98 |
| SbPOE28 | Sobic.001G266100.1 | 1152 | 0.92 |
| SbPOE29 | Sobic.001G266300.1 | 563 | 0.93 |
| SbPSK1 | Sobic.008G034700.1 | 292 | 0.46 |
| SbPSK2 | Sobic.005G035300.1 | 111 | 0.49 |
| SbPSK3 | Sobic.001G448300.1 | 342 | 0.82 |
| SbPSK4 | Sobic.002G021132.1 | 42 | 0.78 |
| SbPSK5 | Sobic.005G155200.1 | 251 | 0.78 |
| SbPSK6 | Sobic.008G034500.1 | 620 | 0.90 |
| SbPSY1 | Sobic.003G131000.1 | 17 | 0.87 |
| SbPSY2 | Sobic.002G199900.1 | 554 | 0.34 |
| SbPSY3 | Sobic.003G121500.1 | 50 | 0.70 |
| SbPSY4 | Sobic.003G272000.2 | 190 | 0.94 |
| SbPSY5 | Sobic.009G215600.1 | 369 | 0.55 |
| SbPSY6 | Sobic.009G177800.1 | 182 | 0.31 |

| Gene name | Transcript ID | Max | τ |
|---|---|---|---|
| SbRALFR1 | Sobic.008G137100.1 | 907 | 0.68 |
| SbRALFR2 | Sobic.008G137300.1 | 49 | 0.64 |
| SbRALF5 | Sobic.003G117500.1 | 92 | 0.80 |
| SbRALF6 | Sobic.003G415500.1 | 8 | 0.62 |
| SbRALF7 | Sobic.003G415400.1 | 6 | 0.73 |
| SbRALF9 | Sobic.003G160700.1 | 198 | 0.55 |
| SbRALF10 | Sobic.003G061600.1 | 135 | 1.00 |
| SbRALF11 | Sobic.002G113500.1 | 63 | 0.61 |
| SbRALF12 | Sobic.008G119400.1 | 298 | 0.74 |
| SbRALF13 | Sobic.003G027300.1 | 23 | 1.00 |
| SbRALF14 | Sobic.009G075400.1 | 36 | 1.00 |
| SbRALF15 | Sobic.002G084200.1 | 26 | 0.98 |
| SbRALF16 | Sobic.003G061700.1 | 73 | 1.00 |
| SbRALFR17 | Sobic.008G027700.1 | 74 | 0.90 |
| SbRALFR19 | Sobic.004G342900.1 | 28 | 0.85 |
| SbRALFR20 | Sobic.010G177800.1 | 5430 | 0.85 |
| SbRALFR21 | Sobic.006G039100.1 | 374 | 0.92 |
| SbRALFR22 | Sobic.010G178000.1 | 1134 | 0.67 |
| SbRALFR23 | Sobic.008G137000.1 | 112 | 0.69 |
| SbRGF1 | Sobic.006G124200.1 | 7 | 0.84 |
| SbRGF3 | Sobic.006G112000.1 | 10 | 0.85 |
| SbRGF5 | Sobic.004G333000.1 | 6 | 0.98 |
| SbRGF7 | Sobic.006G237600.1 | 7 | 0.71 |
| SbTPD1 | Sobic.001G254700.1 | 98 | 1.00 |

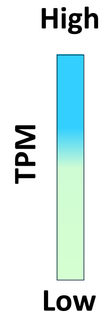


S-Figure 38. Expression of 158 genes encoding small signaling SSPs in 19 small signaling peptide families in leaf, stem, root, and panicle tissues. SSPs were sorted based on phylogenetic order and 158 of the 219 were selected using a >5 TPM threshold across the averaged expression values within each organ. SbECL2 has a max TPM of 3 but was included to not remove the family. Expression values are reported in TPM, with blue indicating high expression and green indicating low expression.
