## Supplementary material for "Analysis of Small Signaling Peptides in *Sorghum bicolor*: Integrating Phylogeny and Gene Expression to Characterize Roles in Stem Development": Tree Files: README.txt

These are standard Newick-format phylogenetic trees made with RAxML.
The trees must be loaded manually into any Newick-compatable viewer.
Examples include iTOL, FigTree(v1.4), or Dendroscope.
If file does not open by double-clicking, please load it manually into the
Newick-compatable viewer of choice.

Note: Some operating systems attempt to preview .tree files and corrupt the
format. Extracting from the ZIP and opening directly from the software avoids
this issue.
